# A potential pathway that links intron retention with the physiological recovery by a Japanese herbal medicine

**DOI:** 10.1101/2023.12.02.569734

**Authors:** Norihiro Okada, Kenshiro Oshima, Akiko Maruko, Trieu-Duc Vu, Ming-Tzu Chiu, Mitsue Nishiyama, Masahiro Yamamoto, Akinori Nishi, Yoshinori Kobayashi

## Abstract

Our previous experiments with klotho mice (i.e., klotho knockout mice) showed that aberrant intron retention (IR) occurs in a pre-symptomatic state of aging and can be restored to the normal state by administration of Juzentaihoto (JTT), a formula of the Japanese herbal medicine (referred to as kampo; Okada et al., 2021). To explain that this phenomenon is not just a specific example observed in klotho mice, but is due to a universal cause that may be common in a pre-symptomatic state, we re-analyzed the data for genes that were differentially expressed in the klotho liver in response to JTT. Our data showed that AMPK signaling was activated in these mice, resulting in inhibition of anabolic pathways including protein synthesis and activation of catabolic pathways including lipolysis in various organs. These data were consistent with the observation that klotho mice appeared to be in a perpetual state of pseudo-starvation, i.e., based on their overall metabolic profile. The abnormally increased incidence of IR (IncIR) in the liver of klotho mice could be classified into three types according to their transcriptional status, revealing that each of all the three types of IncIR was caused by AMPK activation, a consequence of the pseudo-starvation state exhibited by these mice. Furthermore, we showed that both the transcriptional drift variance and the IR drift variance can be useful for visually assessing recovery from the pseudo-starvation state of klotho mice after administration of JTT. With these data, we propose a novel pathway linking intron retention to changes in the physiological state such as starvation, resulting in the maintenance of cellular homeostasis.

## Introduction

Our previous experiments with klotho knockout mice (referred to as klotho mice; Okada et al., 2021) showed that, by 7 weeks of age, intron retention (IR) increases to aberrant levels in several organs including the liver. Interestingly, the retention of a subset of these introns in the liver could be restored to that of the wild-type (WT) levels by administration of a traditional Japanese herbal medicine, i.e., kampo (here we used a formula Juzentaihoto; JTT). Genes whose IR increase was normalized by JTT were found to be mostly involved in liver-specific metabolism. Liver metabolomics data revealed that 3-hydroxybutyric acid, one of the ketone bodies, accumulated in the liver of klotho mice, suggesting that these mice experienced a metabolic state similar to that of starvation (referred to here as ‘pseudo-starvation’). In addition, our analysis of genes that were differentially expressed in klotho mice revealed that liver metabolism did indeed recover to WT levels after JTT administration. In analogy to the widespread accumulation of intron-retained pre-mRNAs induced by heat shock (Shalgi et al., 2014), we proposed a model in which IR in klotho mice is a consequence of age-related stress and JTT-mediated recovery of normal IR actually reflects recovery of liver-specific metabolic functions (Okada et al., 2021).

### Characteristics of klotho mice

Before explaining why we re-analyzed our published data, let us explain the characteristics of the klotho gene, klotho mice, and Japanese herbal medicine. The klotho gene was first identified in 1997 as a putative anti-aging gene in mice (Kuro-o et al., 1997). α-Klotho is expressed only in certain tissues, i.e., kidney, brain, and reproductive organs, and functions primarily to regulate the activation of various signaling pathways, such as the insulin/IGF-1 and Wnt pathways (Kuro-o, 2009). Indeed, mice with increased klotho gene expression tend to have an extended lifespan, whereas mice with decreased klotho expression exhibit premature aging characteristics (Kurosu et al., 2005). *α*-Klotho helps protect neurons from oxidative stress, inflammation, and other aging-related factors, potentially affecting brain health and promoting cognitive decline (Chen et al., 2013; Dubal et al., 2014). Klotho is also involved in regulating the uptake and utilization of minerals, particularly calcium and phosphate, whose proper homeostasis is essential for bone health and the prevention of age-related bone diseases (Kuro-o, 2009, 2010).

Klotho mice exhibit hallmarks of accelerated aging, including premature development of aging-related pathologies like osteoporosis, cognitive decline, cardiovascular disease, and metabolic disorders. Although WT mice typically live for ∼2.5 years, klotho mice live for only ∼100 days (approximately 14 weeks; Kurosu et al., 2005). Klotho mice at ∼7 weeks of age (i.e., half the average lifespan) do not exhibit significant symptoms of abnormal physical aging, and thus this stage represents a pre-symptomatic state (Nagai et al., 2003). However, even at this stage, klotho mice exhibit certain alterations in cognitive ability, renal function, calcium and phosphate metabolism, cardiovascular function, and bone health; notably, in WT mice, all of these processes involve the function of the klotho protein.

### There is an inseparable relationship between Japanese herbal medicine and the pre-symptomatic state of klotho mice

Chinese herbal medicine (UNESCO, 2011) has a long history in China. Japanese herbal medicine, a modification of Chinese herbal medicine, originated in ancient China and is widely used in Japan today to treat a variety of ailments. Unlike chemically synthesized Western medicines, whose effects are carefully evaluated at various stages of testing and whose mechanisms of action are generally rigorously studied, it is extremely difficult to identify the molecular substances responsible for the efficacy of herbal medicines and to analyze their mechanisms of action. This is because the thousands of chemicals contained in Japanese herbal medicines sometimes act synergistically, and their effects are often attributed to metabolites broken down in the intestine (Yamamoto, 2022). Side effects, if any, are said to be relatively minor (Zhou et al., 2016). In any case, the mechanism of action remains a complete mystery, and efficacy can only be evaluated collectively using OMICS methods (Okada et al. 2021).

JTT is one of the formulas of Japanese herbal medicine, consisting of 10 herbs. It is prescribed for patients with fatigue, anemia, night sweats, anorexia, or circulatory problems. JTT stimulates the immune response including enhancement of phagocytosis, cytokine induction, induction of mitogenic activity of spleen cells, and activation of macrophages and natural killer cells (Matsumoto et al., 2000). JTT can also prolong life when combined with the surgical removal of tumors and can protect against the deleterious effects of chemotherapeutic drugs (Wang et al., 2018b).

### Aberrant IR does not simply reflect errors in pre-mRNA splicing but rather plays a distinct role in biological processes

As mentioned above, we found that introns were aberrantly retained during the pre-symptomatic state of klotho mice (Okada et al. 2021). IR is a form of alternative splicing (Jacob and Smith, 2017). Although IR has traditionally been considered deleterious to an organism (Weischenfeldt et al., 2005), recent studies have shown that programmed IR is involved in granulocyte differentiation (Wong et al., 2013), male germ cell differentiation (Naro et al., 2017), and B cell development (Ullrich and Guigo, 2019). Thus, IR is a normal aspect of the routine processing of many pre-mRNAs and appears to be integrated into the regulatory network of RNA processing. Moreover, aberrant IR has even been proposed to be a post-transcriptional signature of aging. Analysis of transcriptomes from mouse, human brain, and Drosophila head has shown an overall increase in IR with aging, suggesting that this process may be evolutionarily conserved (Adusumalli et al., 2019). In addition to the fact that distinct IR patterns have been observed in various physiological and pathological contexts (Liu et al., 2017), we also found that IR is increased in metabolic genes in the liver by starvation stress and returned to a healthy state by treatment with Japanese herbal medicine, suggesting that an increased IR is a natural regulatory response (Okada et al. 2021). To date, however, no clear role for general IR events has been proposed.

### Why did we re-analyze our published data?

We wanted to re-analyze our published data to show that the aberrant IR that occurs in klotho mice during the pre-symptomatic/pseudo-starvation state and the restoration of IR to a healthy state observed with Japanese herbal medicine is not simply a phenomenon unique to klotho mutant mice but due to a more general and definite cause. Klotho is involved in the insulin/IGF-1 pathway (Kuro-o, 2009, 2010), and studies with klotho mice have suggested that glucose metabolism may be altered in this mouse model. Indeed, blood glucose levels (measured during an oral glucose tolerance test) was significantly lower in klotho mice than in WT mice, whereas insulin sensitivity (insulin tolerance test) was higher in klotho mice (Utsugi et al., 2000). The cause of the lower blood glucose levels in the mutant mice may be due to the suppression of nutrient uptake in the intestine despite an adequate dietary intake (Asuzu et al., 2011). Because the effects of hypoglycemia are systemic and not limited to the blood, we wondered if the pseudo-starvation state (i.e., hypoglycemia) of klotho mice might underlie the excessive IR observed in several organs of these mice (Okada et al., 2021, 2022).

### How can IR and a pseudo-starvation state be related?

Here we performed a detailed analysis of transcriptional changes in components of several pathways that are activated downstream of insulin/IGF1 signaling, including the molecular target of rapamycin (mTOR) (Laplante and Sabatini, 2009), AMP kinase (AMPK) (Steinberg and Kemp, 2009), and gluconeogenesis (Hatting et al., 2018), each of which can be affected by the pre-symptomatic/pseudo-starvation state of klotho mice. We found that IR can be affected by physiological changes as well as such transcriptional changes. These results allowed us to classify IR events into three types, which, together with the analysis of gene ontology (GO) enrichment data for each type, provides a basis for evaluating the effects of aging-related stresses based on transcriptome datasets. We conclude that each of all three types of increased IR events (referred to IncIRs) are a consequence of the pseudo-starvation stress experienced by klotho mice. Since treatment with JTT could restore the normal physiological state, we propose a new pathway linking these two aspects̶namely physiological state and IR̶in mice. This proposal is supported by two previous studies in budding yeast showing that intron is required for survival under starvation conditions. In these papers, the requirement for intron during starvation is described in two modes of existence of introns, one as a physical existence in ribosomal genes (Parenteau et al., 2011; Parenteau et al., 2019) and the other as an excised stable form of intron RNA (Morgan et al., 2019). Given the evolutionary context, it is difficult to imagine that this association between IR and starvation-like stress, as reported in budding yeast, will not be found in higher animals.

## 2. Methods

### 2.1. Sample preparation and RNA sequencing

We previously conducted experiments using male *α*-klotho-knockout (*Kl*^‒/‒^/Jcl) mice, referred to as klotho (KL) mice, and male WT C57BL/6JJc1 mice, all aged 3 weeks, obtained from CLEA Japan. All the procedures have been published (Okada et al., 2021). This procedure was repeated twice, and the three replicates were named #1, #2, and #3; we focused on the data from replicate #1, data for which was representative of all three replicates. As a control, the experiment was performed two times with WT mice. The experimental design is described in Fig. S1. All transcriptomes were analyzed by principal component analysis (Fig. S2), demonstrating that the datasets did not contain outliers. Preparation of JTT and RNA extraction followed established protocols (Okada et al., 2021). Quality assessment, filtering, and read mapping based on RNA sequencing data were performed as described, (Martin, 2011, Bolger et al., 2014, Langmead and Salzberg, 2012)

### 2.2. Analysis of alternative splicing using rMATS tool

Significantly altered splicing patterns (*p*-value < 0.05 and difference of IR ratios > 0.05 in rMATS) were identified in three comparisons: KL+/KL‒, KL+/WT, and KL‒/WT, denoting treatment with (+) or without (‒) JTT, using rMATS v.4.0.2 (Shen et al., 2014). Statistical significance was assessed based on skipping junction counts (SJCs) and inclusion junction counts (IJCs) computed by rMATS at the respective loci. In rMATS, the intron ratio at each locus was calculated approximately as IJC / (IJC + SJC × 2). Mapping validation and results were verified using the Integrative Genomics Viewer (http://software.broadinstitute.org/software/igv/) (Robinson et al., 2011).

### 2.3. Calculation of intron length and GC content

Using transcript data from the annotation of the mouse genome (GRCm38 release102) in Ensembl with transcript_name ’201’ (as representative transcript), a general feature format file consisting of intron loci information was created with the AGAT tool (script: agat_sp_add_introns.pl). The general feature format file was used to extract intron sequences from the mouse genome sequence (script: agat_sp_extract_sequences.pl). The AGAT tool package was downloaded from https://github.com/NBISweden/AGAT. Intron length and GC content were calculated from the obtained intron sequences and analyzed with violin plots.

### 2.4. Calculation of transcriptional drift or IR drift, and drift variance

For calculating transcriptional drift (Rangaraju et al., 2015), 15,589 genes were used for which the normalized (trimmed mean of M values method) expression had at least one count in all samples. First, the mean value for each gene of the three WT samples was calculated and used as the reference value (control, *t*_0_). Based on the obtained *t*_0_, from the expression levels of each gene in all samples (*t*), the transcription drift value (TD = log_10_(*t*) ‒ log_10_(*t*_0_)) was calculated. Then, the mean of the TD values for each sample (TD_0_) was then calculated, and from the square root of the difference, the drift variance (DV = (Σ(TD ‒ TD_0_)^2^) / (n ‒ 1)) was calculated (“n” indicates the number of genes used in the calculation). Likewise, the intron ratios at the IR loci of each gene calculated from the IRFinder (Middleton et al., 2017) were used to calculate the average value for each gene of the three WT samples and used as the reference value (control, *i*_0_). Based on the obtained *i*_0_, the intron drift value (ID = log_10_(*i* + 1) ‒ log_10_(*i*_0_ + 1)) was calculated from the intron ratio (*i*) for each gene in all samples. Then, the intron drift variance was calculated.

### 2.5. Enrichment analysis of GO processes and the KEGG pathway

Enrichment analysis was carried out on the DAVID website using significant gene names (Ensembl ID). Any term that was enriched in any of the GO biological processes or KEGG pathways was outputted.

### 2.6. Characterization of IR types

To characterize IR types, scatter plots were generated to depict the fold-change in RNA expression and the difference in intron ratios between WT and KL‒. Each horizontal axis represents the log_2_ fold-change in RNA expression, whereas each vertical axis quantifies the difference in intron ratios between WT and KL‒ (see Fig. 4E). RNA expression was categorized as “not significant” or as “upregulated” or “downregulated” based on a threshold *p*-value of <0.05 and fold-change of >1.2. Introns for which the intron ratio increased significantly in KL‒ were labeled as “IncIR” and those for which the ratio decreased significantly were labeled as “DecIR” (*p*-value < 0.05 in rMATS). Among introns represented in the IncIR category, those with upregulation in KL‒ were categorized as “Activated type,” those with downregulation were categorized as “Suppression type,” and those without significant differences were categorized as “Basic type” for further characterization.

## Results

### Next-generation sequencing and analyses of differentially expressed genes

We re-analyzed the next-generation sequencing data that we published in 2021 (Okada et al. 2021). This yielded 4,737 upregulated and 4,507 downregulated genes in KL‒ (klotho mice not treated with kampo) compared with WT, and 2,283 upregulated and 2,132 downregulated genes in KL+ (klotho mice treated with kampo) compared with KL‒ (Fig. 1AB). Analysis of overlapping genes between the 4,737 upregulated genes in KL‒ compared with WT and 2,132 downregulated genes in KL+ compared with KL‒ revealed 1,440 recovered genes with reverse V-shape (Fig. 1C_left_pair). Similarly, analysis of overlapping genes between the 4,507 downregulated genes in KL‒ compared with WT and 2,283 upregulated genes in KL+ compared with KL‒ revealed 1,574 recovered genes with a V-shape (Fig. 1C_right_pair).

**Fig. 1.**
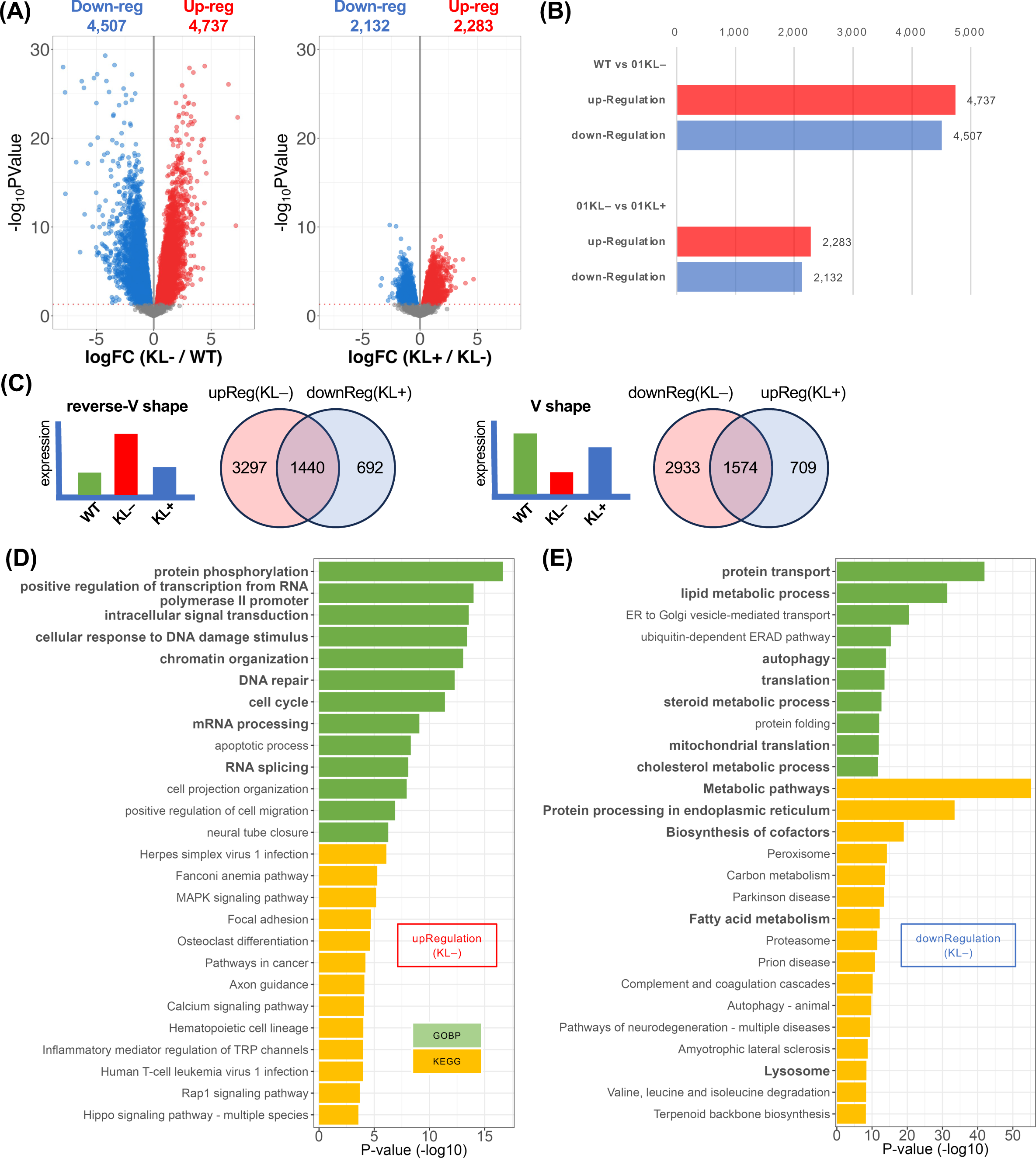
Analysis of differentially expressed genes in KL‒ mice (A) Volcano plots show the comparison of expression levels between WT and KL‒ (left) and between KL‒ mice and KL+ (right). A total of 15,589 genes were used, each of which had normalized (TMM, trimmed mean of M values) expression with more than one count in all samples. The two threshold values for significant differential expression were set at *p*-value < 0.05 (likelihood ratio test) and fold change > 1.2. (B) Bar graphs showing the number of genes for which expression differed significantly. The upper panel indicates genes upregulated in KL‒ mice compared with WT (red) and downregulated in KL‒ mice compared with WT (blue). The lower panel indicates genes upregulated in KL‒ mice compared with KL+ (red) and downregulated in KL+ mice compared with KL‒ (blue). (C) Venn diagrams show how to estimate the number of genes for which expression reverted to the level of WT after administration of JTT medicine. Red circles in the left diagram indicate genes upregulated in KL‒ mice compared with WT, and blue circles indicate genes downregulated in KL+ mice compared with KL‒. The overlapping area represents genes for which expression reverted to that of WT mice with a reverse V-shape after JTT administration (left pair). Likewise, the right diagram indicates the overlap of the number of genes downregulated in KL‒ mice compared with WT (red circles) and upregulated in KL+ mice compared with KL‒ (blue circles). The overlapping area represents the number of genes for which expression reverted to that of WT mice with a V shape after JTT administration (right pair). (D, E) Enrichment analysis of Gene Ontology Biological Processes (green) or KEGG pathways (yellow) using DAVID. The analysis utilized 4,737 genes that were upregulated in KL‒ mice compared with WT (D) and 4,507 genes downregulated in KL‒ mice compared with WT (E).

Enrichment analysis of the 4,737 upregulated genes in KL‒ revealed that certain GO pathways such as protein phosphorylation, transcription-related genes, DNA repair, and cell cycle were especially enriched in klotho mice (Fig. 1D, Table S1), whereas analysis of the 4,507 downregulated genes in KL‒ revealed that pathways such as protein transport, lipid metabolism, and translation were suppressed (Table S2). Because genes involved in translation as well as other anabolic pathways appeared to be repressed in KL‒ mice, these enrichment analyses suggested that the AMPK pathway was activated in KL‒ due to the pseudo-starvation condition of klotho mice.

### Klotho mice are in a state of ‘pseudo-starvation’

Utsugi et al. (Utsugi et al., 2000) reported results of tolerance tests for both glucose and insulin in KL‒ mice, showing lower blood glucose levels compared to WT mice but significantly higher insulin sensitivity. Accordingly, low insulin production and high insulin sensitivity are hallmarks of these mutant mice. Thus, this relatively complex dysregulation of the insulin pathway may underlie the abnormalities we observed in several cellular energy sensing pathways in KL‒ mice.

We have previously reported that KL‒ mice are in a state that mimicks starvation, based on the fact that these mice have elevated levels of 3-hydroxybutylic acid, one of the ketone bodies (Okada et al., 2021, 2022). Therefore, we hypothesized that gluconeogenesis is activated in klotho mice (Hatting et al., 2018; see below). This is consistent with other metabolomics data showing that several glycolytic intermediates such as glucose 1-phosphate, fructose 1,6-diphosphate, and dihydroxyacetone phosphate are lower in KL‒ mice than in WT mice (Okada et al., 2022), suggesting that glycolysis is generally repressed in the liver of KL‒ mice. Consistent with these observations, the abundance of free amino acids such as Arg and Gly were found to be lower in KL‒ mice compared with WT mice (Okada et al., 2022), suggesting that the AMPK pathway is activated and that the mTOR pathway is repressed (see below).

### AMPK activity is increased in response to the pseudo-starvation state of klotho mice

The AMPK pathway senses cellular energy abundance and is activated when cellular energy levels are low, such as during exercise or starvation (Jeon, 2016; Steinberg and Kemp, 2009; Fig. 2A). When activated, AMPK phosphorylates and inhibits mTORC1 in the mTOR pathway as well as HMG-CoA reductase and acetyl-CoA carboxylase, resulting in the suppression of the protein, cholesterol, and fatty acid synthesis, respectively, and ultimately the inhibition of cell growth and proliferation, thereby conserving cellular energy. In addition, catabolic pathways such as fatty acid catabolism are activated upon AMPK activation (Fig. 2A).

**Fig. 2.**
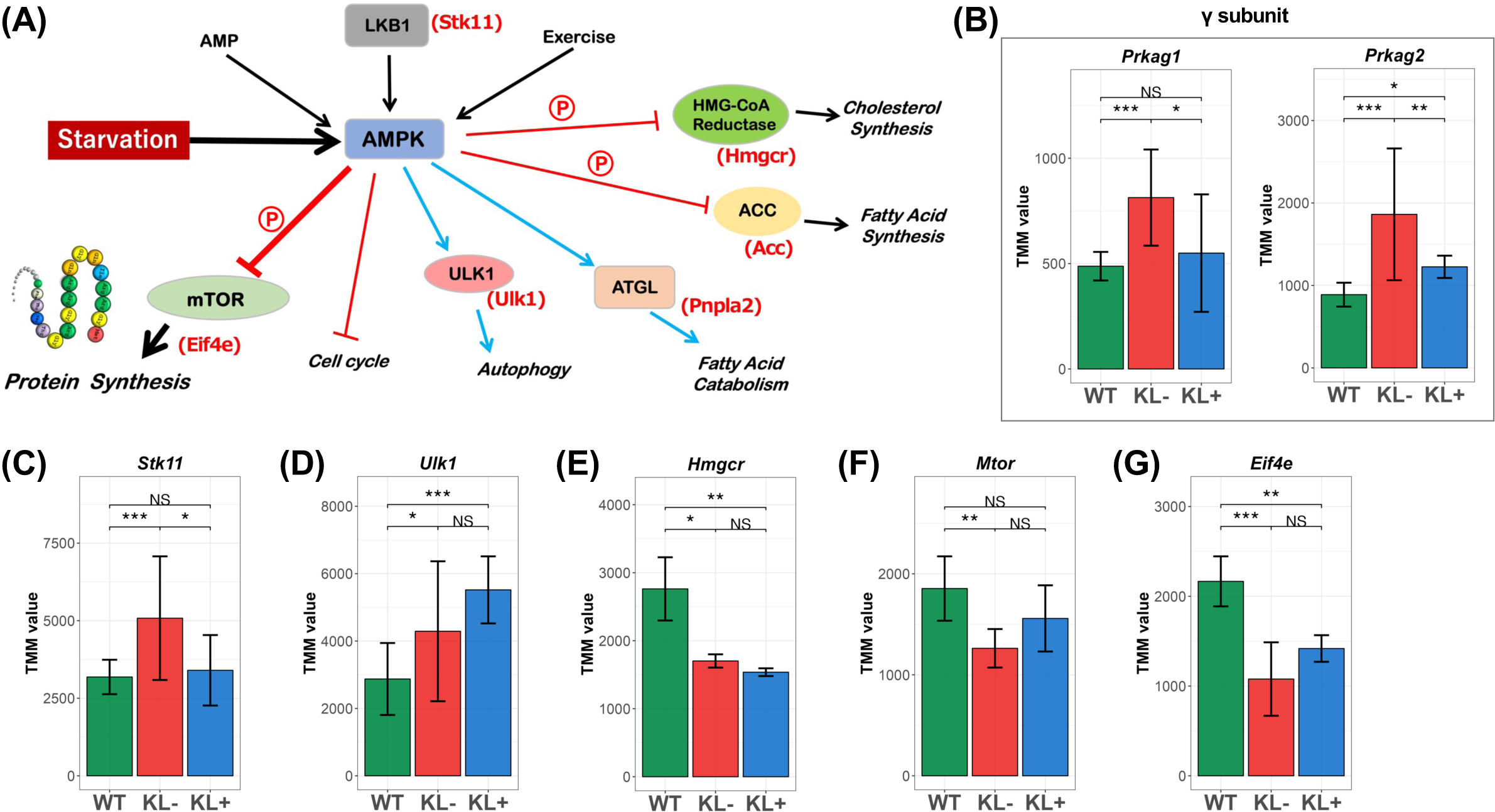
Effects of starvation on the AMPK signaling pathway (A) Schematic representation of several stimuli including starvation stress (black arrows) and responses mediated through the AMPK signaling pathway in mice. Blue arrows indicate stimulation, and red bars indicate suppression. Genes involved are shown in parenthesis in red. The circled red ‘P’ indicates phosphorylation. (B‒G) Bar graph showing the RNA expression level of a gene in WT, KL‒ and KL+. Asterisks indicate the levels of statistically significant differences: \**p* < 0.05, \*\**p* < 0.01, \*\*\**p* < 0.001; NS, not significant (likelihood ratio test). Error bars represent standard deviation. The following genes were analyzed: AMPK gamma subunit (left: *Prkag1*; right: *Prkag2*) (B), *Stk11* (C), *Ulk1* (D), *Hmgcr* (E), *Mtor* (F) and *Eif4e* (G).

To confirm whether the AMPK pathway was activated in our klotho mice due to their pseudo-starvation state (Fig. 2A), we first examined the cellular level of AMPK mRNA in WT, KL‒ and KL+ mice (Fig. 2B). AMPK is a heterotrimeric enzyme conposed of three subunits, namely α, β, and γ, each of which has a specific function (Jeon, 2016; Steinberg and Kemp, 2009). Namely, the α subunit is responsible for directly phosphorylating mTORC1 and thereby inhibiting the mTOR signaling pathway. The β subunit acts as a scaffolding protein that facilitates the assembly and stabilization of the entire AMPK enzyme. The γ subunit is primarily responsible for sensing cellular energy status and plays a critical role in regulating mTOR activity. Fig. 2B shows that the levels of transcripts for the two γ subunits (Prkag1 and Prkag2) were both significantly elevated in KL‒ mice, consistent with the hypothesis that AMPK activation results from pseudo-starvation. Among the γ subunits, those in replicate #1 were upregulated the most in three independent experiments (Fig. S3A). Each of the two γ subunits of AMPK contains multiple binding sites for AMP and ADP, and the binding of either metabolite results in allosteric activation of AMPK, making it more sensitive to phosphorylation by upstream kinases, such as liver kinase B1 (encoded by *Stk11*), which ultimately leads to AMPK activation. Our data that *Stk11* was activated in KL‒ mice (Fig. 2C) confirmed the notion that AMPK was activated in these mice.

### Transcriptional control of catabolic (energy-releasing) and anabolic (energy-consuming) changes induced by the pseudo-starvation of klotho mice

Upon activation of AMPK, general catabolism is activated. As expected, a gene involved in autophagy, namely *Ulk1*, was found to be activated in klotho mice (Fig. 2AD). ULK1 is a catabolic enzyme that facilitates the removal and degradation of unnecessary cellular components, resulting in the release of nutrients for energy production and recycling cellular materials (Shang and Wang, 2011). Another catabolic process, namely, fatty acid oxidation, was also activated in klotho mice. In the first step of fatty acid oxidation, stored triglycerides are broken down into fatty acids and glycerol by adipose triglyceride lipase (*Pnpla2*) (Wang et al., 2018a; Fig. 3A). Although this process occurs primarily in adipose tissue, transcription of the gene encoding this lipase was also very high in the liver in KL‒ compared to WT mice (Fig. 3B), again suggesting that klotho mice have a relatively high energy requirement (for all three experimental replicates, see Fig. S3F). Free fatty acids are transported into mitochondria, where β-oxidation occurs, and the enzyme encoded by *CPT1* is responsible for the first step of β-oxidation (Houten and Wanders, 2010). Our data showed that the transcription of *CPT1* was also upregulated as evidenced by the average value for the three experimental replicates (Fig. S3G), although there was some variation between the replicates (Fig. 3C).

**Fig. 3.**
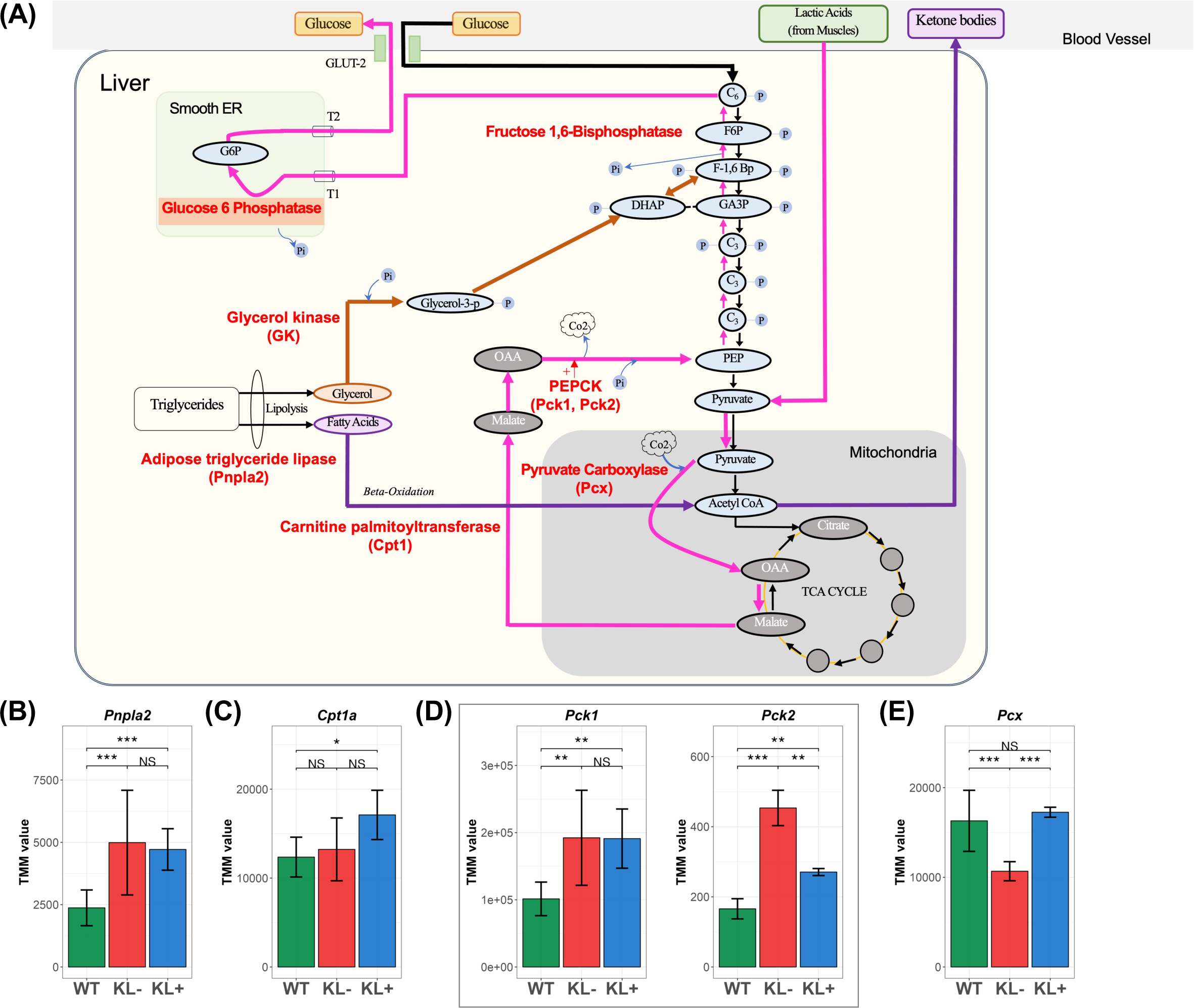
Effects of starvation on energy production (A) Schematic representation of pathways of gluconeogenesis, lipolysis, and ketogenesis under starvation conditions under normal nutrient conditions. (B‒E) Bar graph showing the RNA expression level of a gene in WT, KL‒ and KL+. Asterisks indicate the levels of statistically significant differences: \**p* < 0.05, \*\**p* < 0.01, \*\*\**p* < 0.001; NS, not significant (likelihood ratio test). Error bars represent standard deviation. The following genes were analyzed: *Pnpla2* (B), *Cpt1a* (C), *Pck1* (left) and *Pck2* (right) (D), and *Pcx* (E).

With respect to anabolic (energy consuming) pathways, we first determined whether cholesterol biosynthesis was repressed in klotho mice. HMG-CoA reductase participates in cholesterol biosynthesis (Soto-Acosta et al., 2017), and this enzyme is directly phosphorylated (and consequently inhibited) by the α subunit of AMPK. Our data indicate that transcription of the gene encoding HMG-CoA reductase (*Hmgcr*) was suppressed in KL‒ mice (Fig. 2E), likely due to inhibition of *Hmgcr* expression by AMPK.

The mTOR pathway promotes cell growth and proliferation by upregulating protein synthesis, ribosome biogenesis, and cell cycle progression. The cellular functions of the mTOR pathway have been elucidated mainly by analyzing protein-protein interactions involved in the phosphorylation of pathway components (Fig. 2A, S4), and the transcriptional control of the mTOR pathway remains mostly unknown. Therefore, to determine the outcome of transcriptional changes of the entire mTOR pathway based on RNA-sequencing data (apparent suppression of mTOR, see Fig. 2F), it is necessary to examine the end-products of the pathway is required, i.e., because the transcriptional network is too complex to easily infer the outcome of the pathway (Nijland et al., 2007). Among the 10 end products of the mTOR pathway (Fig. S5), three are involved in the modulation of protein synthesis (Fig. S5), namely eIF4E, eIF4B and S6. eIF4E is involved in cap-dependent translation initiation, it’s cap-binding activity is essential for the recruitment of the ribosome and other initiation factors to the mRNA, and eIF4E is a component of the eIF4E complex that helps recruit the small 40S ribosomal subunit to the 5’ cap structure of the mRNA. After recruitment, the ribosome scans the 5’ untranslated region of the mRNA until it recognizes the start codon. Accordingly, the amount of eIF4E is proportional to the translational activity. As shown in Fig. 2G and Fig. S5, the abundance of mRNAs encoding both eIF4E and eIF4B was significantly suppressed, supporting the notion that translation is significantly suppressed in KL‒ mice.

### Upregulation of gluconeogenesis-related genes in klotho mice

During starvation, pyruvate, lactate, glycerol and certain amino acids such as lysine, derived from protein degradation in skeletal muscle or produced from muscle, serve as substrates for gluconeogenesis starting from non-carbohydrate precursors (Hatting et al., 2018; Fig. 3A). To investigate whether gluconeogenesis is activated in KL‒ individuals, we first examined the expression of the gene encoding phosphoenolpyruvate carboxykinase, which is involved in the conversion of oxaloacetate to phosphoenolpyruvate. This carboxykinase is key for gluconeogenesis, which occurs primarily in the liver and is essential for maintaining blood glucose levels during fasting or prolonged exercise when glucose is not readily available from dietary sources. Fig. 3D and Fig. S3D show the expression levels for the two genes (*Pck1* and *Pck2*) encoding isozymes of phosphoenolpyruvate carboxykinase. These genes were significantly upregulated in each of our experimental replicates, suggesting that gluconeogenesis does indeed occur in KL‒ mice. We next examined the expression of *Pcx* (encoding pyruvate carboxylase), another enzyme involved in gluconeogenesis. Although the transcription of *Pcx* was upregulated on average, transcription varied between the three replicates (Fig. 3E and Fig. S3E).

Taken together, the results of the five sections written thus far indicate that the pseudo-starvation state of klotho mice enhances the AMPK pathway, which activates autophagy and fatty acid degradation and suppresses the biosynthesis of proteins, lipids, and cholesterol. In addition, glucose deprivation in klotho mice activates gluconeogenesis.

### Classification of IncIR and DecIR based on a scatterplot

We quantified the IR genes that were either increased or decreased in KL‒ mice compared to WT (Fig. 4A). Consistent with our previous data (Okada et al., 2021), the number of IncIR was much larger than that of DecIR (Fig. 4A). However, the number itself increased significantly compared to our previous data (Okada et al., 2021). This was likely due to an increase in accuracy resulting from the addition of five individuals in the WT control group (Fig. S2).

When the amount of transcription of a gene did not differ between WT and KL‒, an increase in IR for that gene tended to decrease the translatability of the mRNA, resulting in decreased abundance of the protein product. Thus, both the downregulation of a gene’s expression and its increased IR had a very similar effect, namely decreased abundance of the protein product, although these two aspects are based on distinct cellular mechanisms. Downregulation of gene expression occurs through transcriptional control and is generally triggered by extrinsic changes including stress. In klotho mice, pseudo-starvation activates the AMPK pathway, leading to the downregulation of numerous genes involved in anabolic processes including protein synthesis (Figs. 2 & 3), so that energy expenditure should be suppressed. In contrast to such control of the transcriptional level, increased IR represents the temporal control of pre-mRNA processing to adjust protein production by increasing pre-mRNA abundance in the nucleus and decreasing the pool of translatable mRNA in the cytoplasm, which also favors energy conservation in a pseudo-starvation state by adjusting the amount of translatable mRNA to an appropriate level in the cytoplasm.

Based on the above discussion, we compared the enrichment of the downregulated genes in KL‒ (Fig. 1E) with that of the IncIR genes in KL‒ (Fig. 4C), revealing enrichment of many KEGG and GO terms relevant to the AMPK/mTOR pathway). Genes related to several metabolic processes, cofactor biosynthesis, and the lysosome (which degrades proteins to produce amino acids for the mTOR signaling pathway) (Fig. 1E) were commonly enriched for the downregulated genes in KL‒ and the KL‒ IncIR genes. Thus, at least some of the enriched terms in both groups were derived from the same cellular stress, namely starvation.

We then examined the IR genes that were increased or decreased in KL+ compared with KL‒ mice (Fig. 4A). Venn diagram analysis revealed 776 recovered loci with a reverse V-shape (Fig. 4B_upper) and 39 recovered loci with a V-shape (Fig. 4B_lower). Enrichment analysis of the 681 genes corresponding to the 776 loci revealed enrichment of metabolic pathways, lysosomes, and spliceosome biogenesis in KEGG pathways with a range of 6.9E-6 to 3.1E-3 (Fig. 4G_right, Fig. S6). In Fig. 4G, we compared enrichment terms between the 1,574 genes for which expression was recovered with a V-shape (Fig. 1C_right, Table S4) and 776 IncIR genes (Fig. 4B_upper) for which IR was recovered with a reverse V-shape (Table S3). It is clear that although not all the genes for which IR was restored to the WT state in response to JTT corresponded to those that were downregulated and recovered in klotho mice, many genes associated with catabolism or lysosomal function were enriched in these two systems.

To further clarify the relationships between gene expression and IR, we classified the IR genes whose expression changed in response to JTT using a scatterplot (Fig. 4D & E). As shown in the schematic representation in Fig. 4D, such genes could be divided into four main groups, namely upregulation and IncIR (Fig. 4D_right_top), downregulation and IncIR (Fig. 4D_left_top), upregulation and DecIR (Fig. 4D_right_bottom), and downregulation and DecIR (Fig. 4D_left_bottom), in addition to two “not significant” groups. Because the number of DecIR genes was small (Fig. 4D, E), we focused only on the IncIR genes, which could be grouped into three types as noted in the next section.

### Categorization of three types of IncIR, most transcriptional changes of which were a consequence of the pseudo-starvation state of klotho mice

Among three types of IncIR, a total of 253 IncIR genes that were downregulated in KL‒ (Fig. 4F, downregulation/IncIR) were designated as the suppression type (S-type). Enrichment analysis of these genes revealed downregulation of anabolic pathways such as those involved in metabolic processes, translation-related pathways, and cofactor biosynthesis (Fig. 5B_left, Fig. S7). The downregulation of these pathways was caused by AMPK activation or mTOR repression induced by pseudo-starvation, which we described in the previous section (Fig. 2). Patterns of examples of three loci drawn by the Integrative Genomics Viewer, namely Dpp3, Dist and Decr2 are shown in Fig. 5A_upper, where examples of IncIR with full recovery of IR were selected. As evidenced by the downregulation data for KL‒ compared to WT (Fig. 5A_upper), we termed this the <u>s</u>uppression type (S-type).

**Fig. 4.**
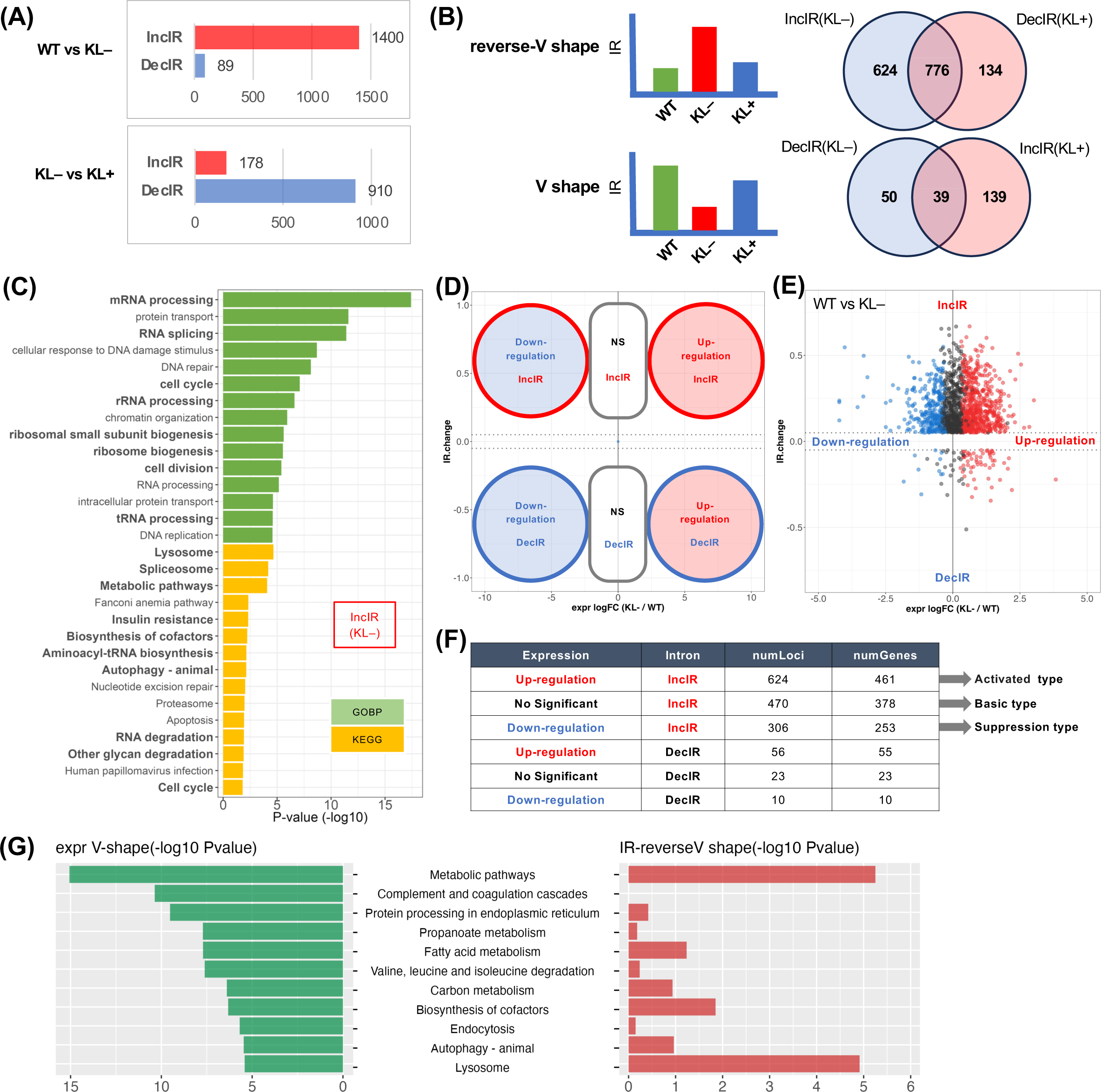
IncIR characterized into three types based on relative gene expression (A) Bar graphs show the number of loci with significantly increased IR (IncIR) or decreased IR (DecIR) based on two different comparisons (*p* < 0.05 and difference of intron ratio > 0.05 in rMATS). The left panel indicates the comparison between WT and KL‒ mice, for which the red bar indicates the number of loci with IncIR in KL‒ and the blue bar indicates DecIR in KL‒. The right panel indicates the comparison between KL‒ mice and KL+, for which the red bar indicates the number of loci with IncIR in KL+ and the blue bar indicates DecIR in KL+. (B) Venn diagrams show how to estimate the number of loci for which IRs were recovered after administration of JTT medicine. The blue circle in the top panel indicates the number of loci with IncIR in KL‒ mice compared with WT, and the red circle indicates DecIR in KL+ mice compared with KL‒. The overlapping area represents the number of loci for which IR levels were recovered with a reverse V-shape after JTT administration (top). Likewise, the bottom panel shows the overlap of the number of loci between DecIR in KL‒ mice compared with WT in blue and IncIR in KL+ mice compared with KL‒ in red. The overlapping area represents the number of loci for which IR levels were recovered with a V-shape after JTT administration (bottom). (C) Enrichment analysis using 1,092 genes with a locus with significantly IncIR (1,400 IncIR loci) between WT and KL‒. Green indicates enriched terms for Gene Ontology Biological Processes, and yellow indicates enriched terms for KEGG pathways. (D) Schematic representation of a scatter plot of changes in RNA expression and IR, for which the horizontal axis represents the log_2_FC value of RNA expression between KL‒ mice and WT, and the vertical axis represents the difference of the ratio of IR in KL‒ mice compared with WT (IncIR or DecIR). (E) Scatter plot of changes in RNA expression and IR in KL‒ mice compared with WT. Each plot shows loci (or genes with that locus) with significantly altered IR. Red dots indicate significantly upregulated genes, blue dots indicate significantly downregulated genes, and gray dots indicate genes for which expression did not change. (F) The number of loci and genes in each group classified in panel E. The classification of three types is shown. (G) Comparison between two different enrichment analyses of genes for which RNA expression (1,574 genes) were recovered with a V-shape (Fig. 1C) shown in the left panel and IR was recovered with a reverse V-shape (687 IR genes, 776 loci; Fig. 4B) shown in the right panel.

**Fig. 5.**
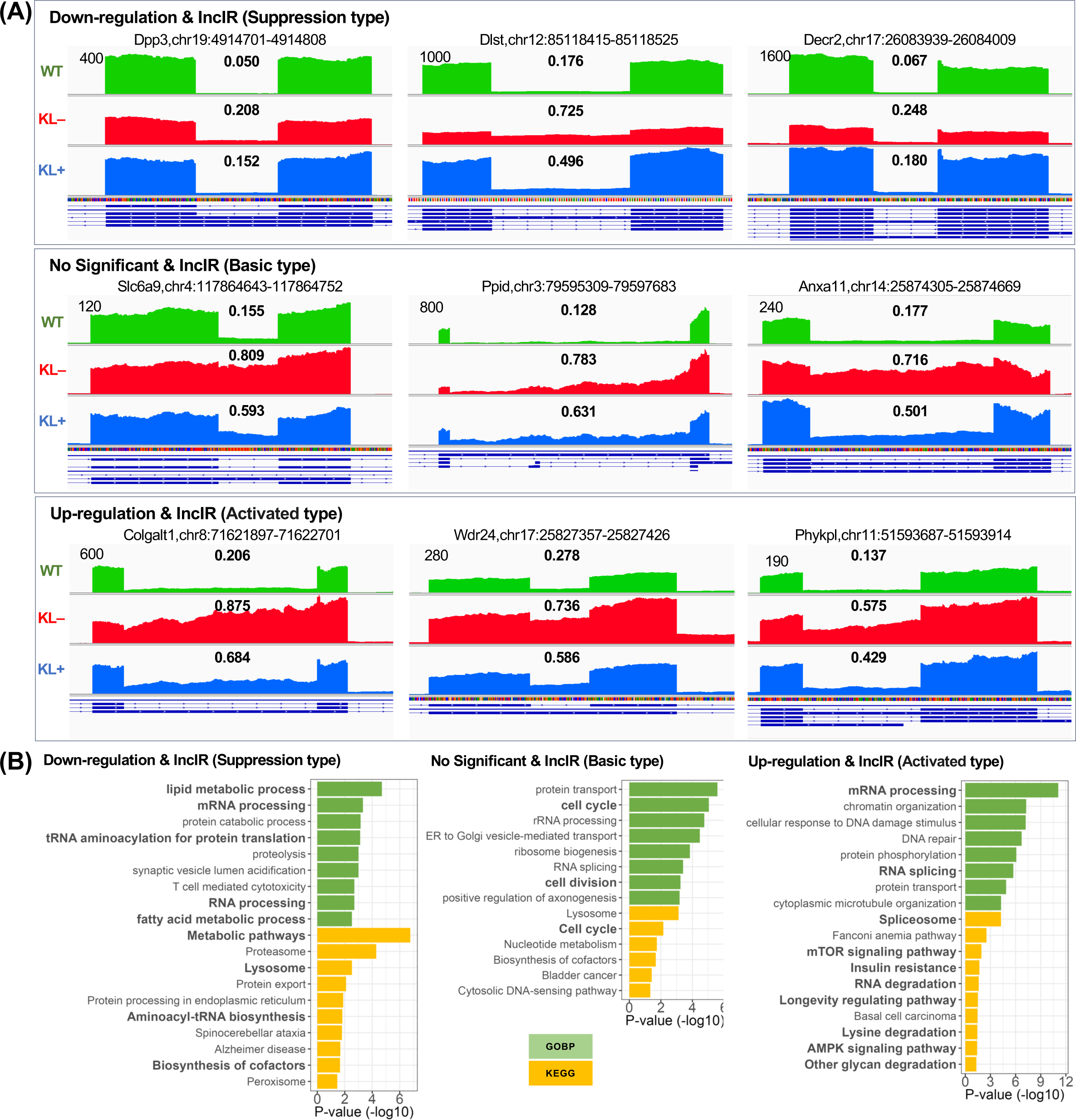
Characterization of three types of IncIRs (A) Mapping results of intron loci selected based on a significant difference between KL mice and WT using Integrative Genomics Viewer (IGV). The upper row shows increased RNA expression and increased IR ratio (activated type), the middle row shows no significant increase or decrease in RNA expression and increased IR ratio (basic type), and the lower row shows decreased RNA expression and increased IR ratio (suppression type). Mapping results are shown as wild type (WT, green), KL‒ (red), and KL+ (blue). Bold numbers in the center indicate intron ratios. (B) Results of a gene enrichment analysis for each type. Green indicates Gene Ontology Processes, and yellow indicates KEGG pathways.

The 378 genes with IncIR for which the transcriptional output did not differ between WT and KL‒ (Fig. 4F; Fig. 5_middle) were designated as the <u>b</u>asic type (B-type; Fig. 4F). Although the B-type category was created due to technicalities of the scatterplot analyses, the interpretation of the biological significance of the B-type was essentially the same as that of the S-type in terms of suppression of anabolic processes for the KL‒ mice. In fact, when the enrichment analysis was performed for the 631 IR genes combined between the S-type and B-type, the top-ranked terms were all related to translation and RNA processing (Fig. S12), indicating that they were the result of starvation-induced inhibition of anabolic processes. For the 378 IR genes (Fig. 5B_middle), terms related to cell division and cell cycle were particularly enriched. Notably, the same terms were similarly enriched in the other two experimental replicates (Fig. S11), suggesting that suppression of cell division and the cell cycle by controlling the level of processing for the pre-mRNA of the B-type genes is a common feature of mouse cells.

The 461 genes belonging to “upregulation/IncIR” in Fig. 4F were designated as A-type (<u>a</u>ctivated type; Fig.5A_lower). Interestingly, several KEGG terms related to energy control such as mTOR signaling, insulin signaling, and AMPK signaling and the energy production system such as lysine degradation and other glycan degradation were enriched in these genes (Fig. 5B_right, Fig. S7). Enriched genes involved in mTOR signaling, lysine degradation and AMPK signaling are listed in Fig. 6A. Fig. 6B shows the expression patterns of seven enriched genes in AMPK signaling, showing clear elevated transcript patterns of genes in KL‒, several of which were significantly restored in KL+. Genes encoding Pck2, Prkag1, and Stk11 were previously discussed in the context of increased gluconeogenesis (Fig. 3A), AMPK subunit activation (Fig. 2B), and AMPK activation by liver kinase B1 (Fig. 2A), respectively, all of which were induced by the pseudo-starvation state of klotho mice. In addition, lysine degradation is a process of gluconeogenesis induced by starvation because lysine catabolism leads to the production of acetyl-CoA, the starting material for the TCA cycle (Fig. 3A). Lysine catabolism is also involved in ketogenesis in starved animals because acetyl-CoA is also a starting material for the biosynthesis of ketone bodies (Fig. 3A). Accordingly, although not all of the enriched A-type genes were not well explained due to a starvation-like state (e.g., spliceosome-related genes, Fig. 5B_right, S10), it is reasonable to conclude that the activation of A-type genes involved in energy balance and production could be due to the pseudo-starvation state of klotho mice.

**Fig. 6.**
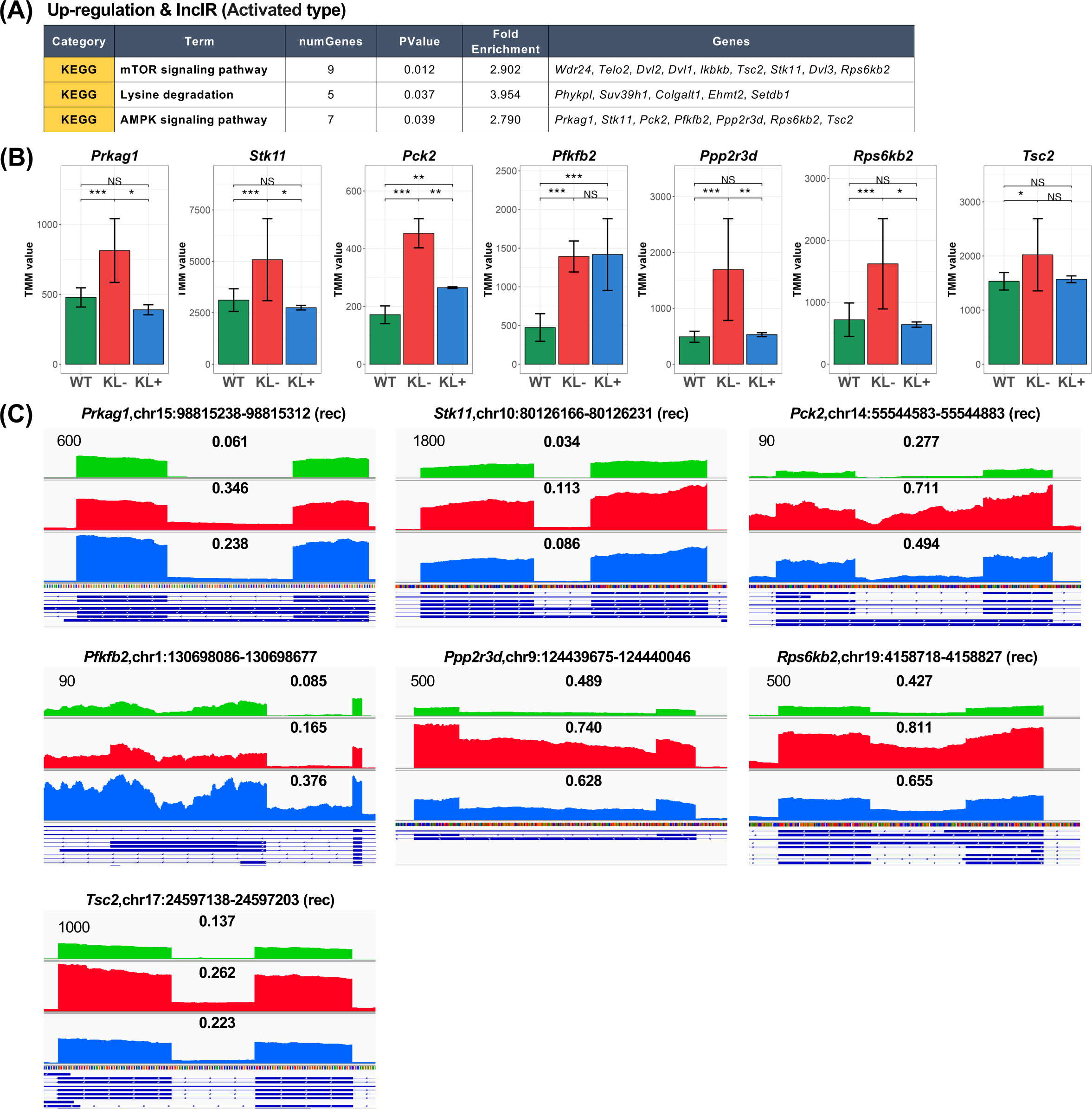
Expression analyses of genes in three signaling pathways belonging to the A-type (A) Three enriched signaling pathways and these associated genes. (B) All the patterns of expressions of seven genes involved. (C) Mapping results of seven intron loci using Integrative Genomics Viewer (IGV).

Taken together, all of the transcriptional changes measured for these three types of IncIR, i.e., A-, B- and S-types, could be explained primarily as a physiological consequence of pseudo-starvation in klotho mice. Thus, the accompanying IR changes are also a consequence of pseudo-starvation (see later section).

### Tuning intron hypothesis: a new mechanism to modulate gene-product abundance under stress

In the case of IncIR in the A-type (Fig. 5A_lower), a situation arose in which transcription was activated and yet IR increased. The existence of such opposite movements in the context of protein production suggests that increased IR is a mechanism for fine-tuning the supply of mRNAs for protein synthesis. It is well known that, in terms of cellular physiology, the entire process of gene expression is costly in terms of both time and energy, e.g., due to the need for poly-A addition, 5’ cap formation, pre-mRNA splicing, and mRNA export and targeting. In other words, it is almost impossible that transcriptional control alone could ensure that a cell produces the exact amount of a particular mRNA needed for normal cellular function. A much more reliable regulatory mechanism would be to initially produce an excess of pre-mRNAs and then use turnover to modulate the amount of each mRNA that is needed in a particular cellular context. Could it be that IR provides a mechanism for such fine-tuning, whereby immature mRNAs are stored in the nucleus and then exported to the cytoplasm as needed?

A tuning intron model is shown in Fig. S14, in which includes both DecIR and IncIR genes. As explained above, IncIR occurs when adjustments by reducing the cellular abundance of any particular protein(s) are needed, whereas DecIR occurs when more of a particular protein(s) is needed. Given the known functions of the nuclear pore, it is likely that such adaptations (i.e., IncIR and/or DecIR) occur via modulation of nuclear pore activity (Fig. S14_left). Kwiatek et al. (2023) reported a critical role for human PABPN1 (polyadenylate-binding nuclear protein 1) in preventing the nuclear export of unspliced pre-mRNAs i.e., via IR, as a means of controlling protein abundance, suggesting the existence of a mechanism for fine-tuning gene expression via IR. If IR events are indeed responsible for fine-tuning post-transcriptional dynamics to accurately reflect the physiological conditions in a cell, then IR events may be one of the key mechanisms for maintaining cellular homeostasis during stress.

### Cellular homeostasis can be maintained via fine-tuning IR of short introns to aid the physiological recovery of klotho mice after treatment with JTT

We have previously reported that administration of JTT could restore the pre-symptomatic IR profile of klotho mice to that observed in WT mice (Okada et al., 2021) and that such introns are characterized by being shorter in length and richer in GC content than other introns (Okada et al., 2021). The data shown in Fig. 7A & B confirmed this observation. To avoid bias due to differences in GC content in a chromosome, known as chromosome bands (i.e., GC-rich R bands and AT-rich G bands), we examined differences in intron length and GC content by comparing loci that undergo IR with other loci in the same gene that do not undergo IR. The results confirmed the relevance of two features, namely short intron length (Fig. 7E_left) and relatively high GC content (Fig. 7E_right) in our retained introns. In addition, we showed that the locus with recovered intron in a gene is not confined in a particular location (Fig. 7D).

**Fig. 7.**
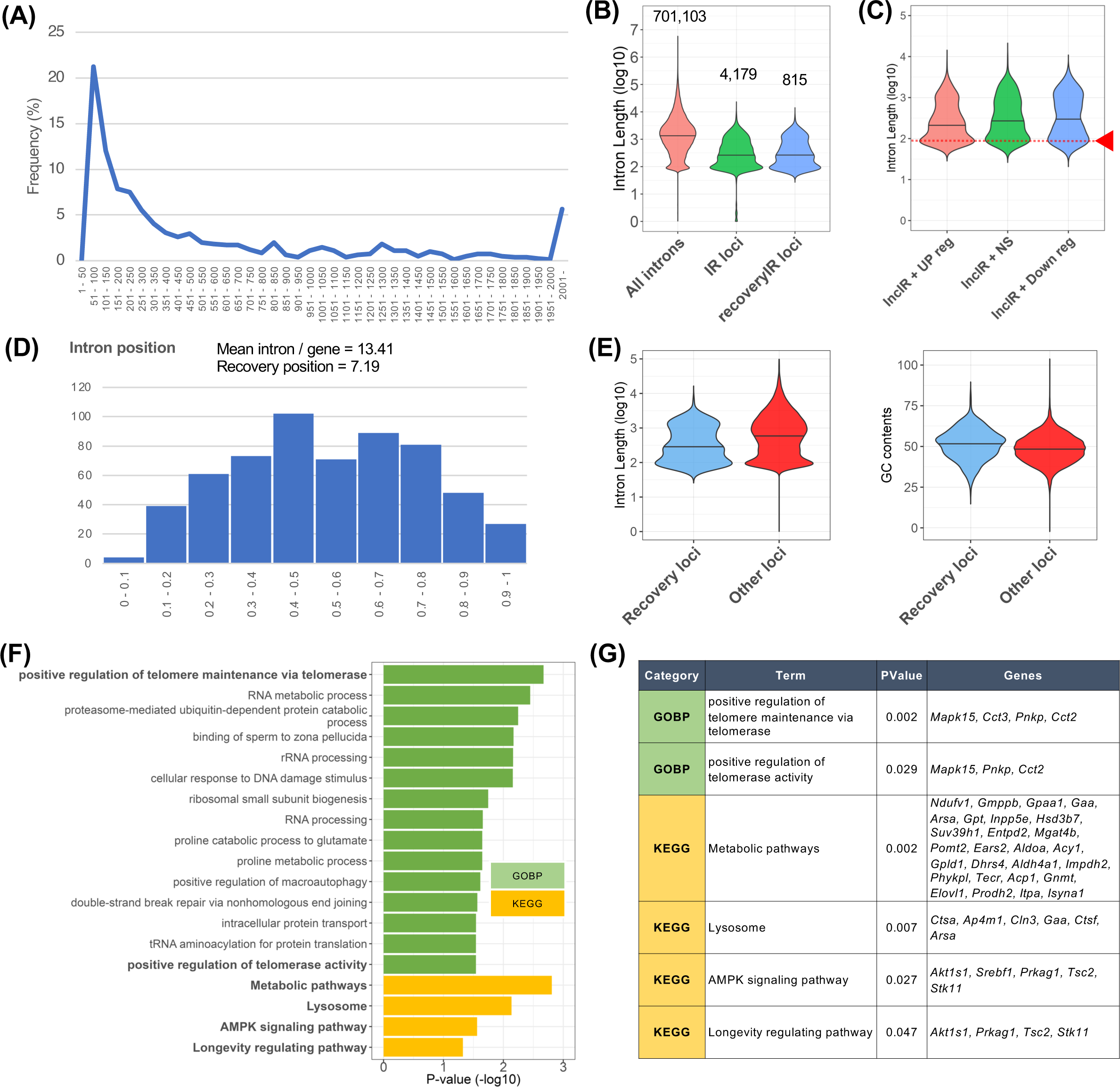
Characterization of short introns for which intron status returned to normal in response to physiological recovery to a healthy state after administration of JTT (A) Characterization of intron length distribution of 815 loci (776 reverse-V type + 39 V type loci) that were recovered after administration of JTT. The horizontal axis shows the range of intron lengths, and the vertical axis shows their frequency. (B) Violin plot of intron length showing that short introns were enriched. The red plot indicates all intron loci predicted from the annotation file (701,103 loci), the green plot indicates intron-retained loci predicted from the annotation file by the intron detection tool rMATS (4,105 loci), and blue plot indicates loci with recovered IR (815 loci). (C) Violin plot of intron length showing the slight difference among the three types of IncIR. The red plot (461 loci; upregulation/IncIR) indicates that the smaller introns were enriched in this A-type more than others, and the green plot (378 loci; no significant/IncIR) and blue plot (253 loci; downregulation/IncIR) indicate the patterns for B-type and S-type, respectively, with no difference in their distribution. (D) Bar graph showing the distribution of positions for 815 recovered intron loci. For example, when a particular gene has 15 intron loci and the 9th intron is recovered, the position is set to 0.6 (= 9/15). (E) After extracting 687 genes including 701 of the 776 loci of the reverse-V shape type, the sequences of all introns in 8,405 loci in the 687 genes were extracted. “Recovery loci” indicates the recovered 701 loci, and “Other loci” indicates the other 7,704 loci. Violin plots showing intron length (left) and GC content (right). (F) Enrichment analysis (Gene Ontology Biological Processes and KEGG pathways) was performed using DAVID for 177 genes with upregulated/IncIR (A-type) that had fewer than 100 bases in KL‒ mice compared with WT. Green indicates enriched Gene Ontology Biological Process terms, and yellow indicates enriched KEGG pathways. (G) List of IR genes in several highlighted terms of GO in panel G, which are involved in longevity.

Furthermore, among the three types of IncIR (i.e., A-, B-, S-type), the shorter introns were more enriched in the A-type than in the other two types (highlighted by an arrow in Fig. 7C). Interestingly, GO analysis of 177 genes containing such short introns (<100 nucleotides) in the A-type showed that genes responsible for longevity (e.g., *Akt1s1*, *Prkag1*, *Tsc2*, and *Stk11*) including those involved in telomere regulation (e.g., *Mapk15*, *Cct3*, *Pnkp*, and *Cct2*) were highly enriched in this group of genes with short introns (Fig. 7FG, Fig. S13), suggesting a link between longevity and retention of short introns. In the previous section, we noted that IncIR in the A-type includes pathways involved in energy production, which is indispensable for survival in a pseudo-starvation state. This response is similar to that of HeLa cells subjected to heat shock, in which protein synthesis is largely halted by inhibition of splicing; however, translation continues for mRNAs encoding molecular chaperones that have protein folding functions and thus are thus necessary for survival (Shalgi et al., 2014). Our current data suggest that genes responsible for longevity are activated under stress, and are mediated by splicing control of the short introns in mature mRNAs.

As noted above, the spliceosome complex is too large to efficiently recognize the splice sites of short introns (Fukumura et al., 2021; Wahl et al., 2009). The A complex of the spliceosome is an asymmetric globular particle (∼26 × 20 × 19.5 nm; Behzadnia et al., 2007) that occupies 79‒125 nucleotides of the pre-mRNA (see Zhang et al., 2019, for a review of the architecture of the A complex). Interestingly, human ultrashort introns (43‒ 65 nucleotides) are removed relatively efficiently (Sasaki-Haraguchi et al., 2012). In this regard, Mayeda’s group recently reported that the removal of a distinct subset of short introns requires the newly defined splicing factor SPF45/RBM17 but not U2AF, which is a general splicing factor (Fukumura et al., 2021). Thus, it would be interesting to determine how many SPF45-requiring short introns are aberrantly retained in response to a stress but are then appropriately excised after JTT administration. We speculate that short introns and special splicing factors such as SPF45 may have evolved for specific roles, such as prolonging lifespan even under starvation.

### Suppression of transcriptional drift and IR drift in response to JTT medicine

In previous discussions, we have seen that klotho mice are subjected to pseudo-starvation-induced aging stress. We have also seen that administration of JTT ameliorates some of these stresses. But using the RNA-seq data, is there a way to show that the body as a whole actually improves? This is a very important question because JTT can have side effects as well as positive effects.

If aging is considered in terms of gene expression, then aging can be defined as a process in which the coordinated expression of various genes in youth is gradually disrupted, with certain genes being upregulated and others downregulated (Gupta et al., 2021). Based on this definition, Rangaraju et al. (2015) devised a method to quantify this drift from such coordinated expression based on a defined index using *Caenorhabditis elegans* as a model. This method is called “transcriptional drift”, and the concept of “transcriptional drift variance” (Tdv) was devised to measure the extent to which aging affects biological processes and the efficacy of anti-aging drugs such as mianserin. These authors showed that mianserin could extend the lifespan of *C. elegans* as evidenced by the suppression of Tdv. Using the same strategy described above, we evaluated the pre-symptomatic state in klotho mice as well as the extent of recovery to the healthy state in response to JTT treatment (Fig. 8A).

**Fig. 8.**
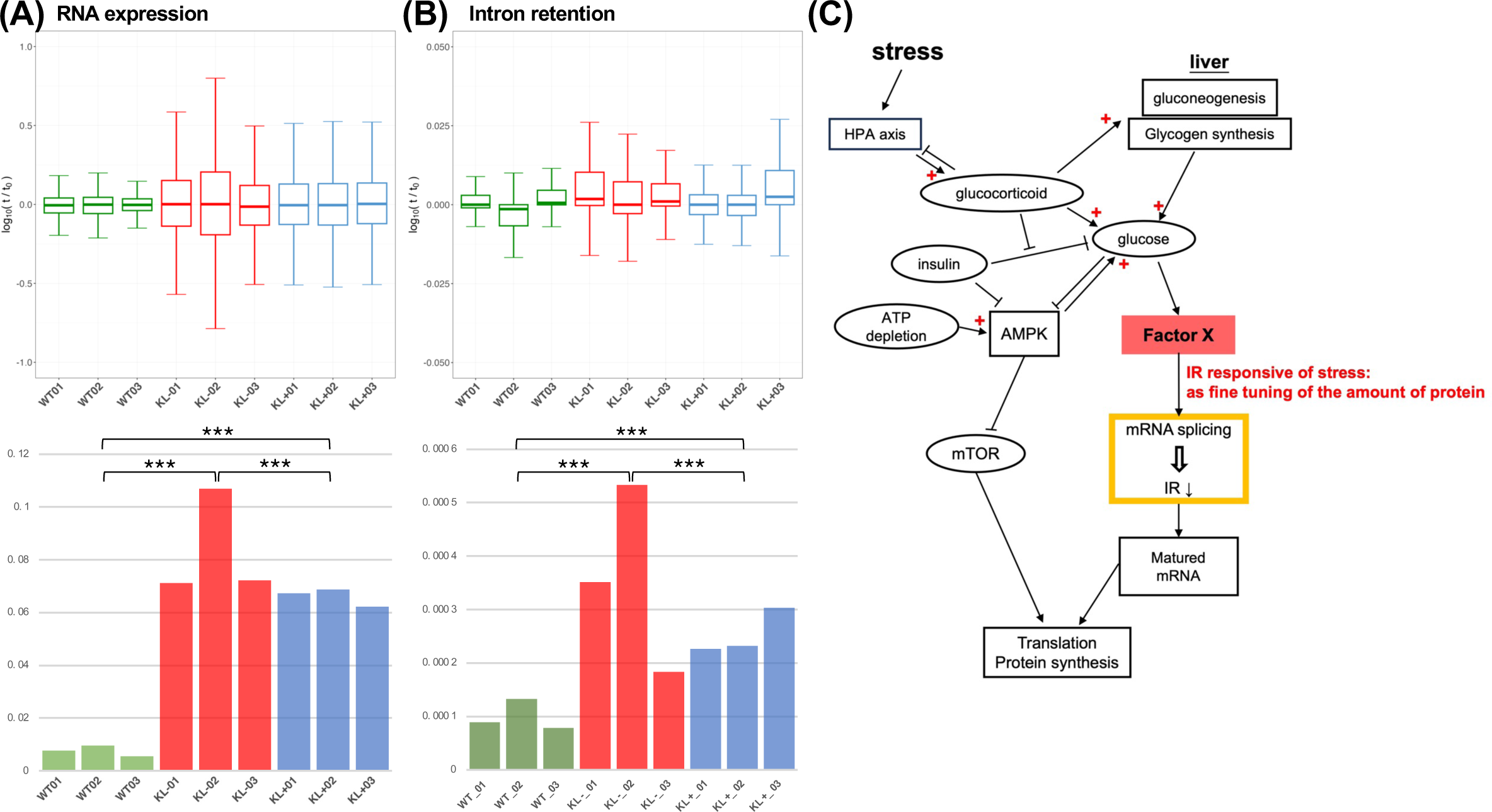
Suppression of transcriptional and IR drifts by JTT medicine (A) (upper) Box plot showing transcriptome drift calculated using 15,589 genes with RNA expression greater than the minimum TMM value of 1. (Lower) Bar graph showing the drift variance calculated by integrating the deviations. Asterisks indicate the level of the statistically significant difference: \*\*\**p* < 0.001, Mann‒Whitney *U* test. (B) (upper) Box plot showing IR drift calculated from IR ratios calculated using the IRFinder tool. (Lower) Bar graph showing the drift variance of IR. (C) Schematic representation of a link between cellular energy status and IR. See text for details.

The Tdv values of the pre-symptomatic state of klotho mice were much higher than those measured in WT mice (red bars in Fig. 8A_lower), whereas the Tdv values calculated after JTT administration were significantly decreased, indicating that JTT counteracts the disruption of coordinated transcription caused by aging (blue bars in Fig. 8A_lower).

As mentioned above, the pre-symptomatic state of klotho mice has its own characteristic transcriptional pattern, but this pattern can be altered by the JTT administration. Although our global analysis of differentially expressed genes allowed us to characterize some reversion of the transcriptome in klotho mice to that of the WT state in response to JTT, this analysis did not necessarily indicate that liver cells as a whole experienced such a reversion. Therefore, an analysis using Tdv as the determinant was an excellent method to assess the recovery status in the liver.

Based on the knowledge that IR was increased in pre-symptomatic klotho mice and that certain mice experienced a reversion of IR to that of WT mice upon administration of JTT, we reasoned that the method applied to the transcriptome data could be applied to the IR data. Therefore, we calculated IR drift and IR drift variance values using IRFinder data derived from the same dataset used in the present study. The results showed that IR drift variance was significantly increased in KL‒ and decreased in KL+ mice (red and blue bars, respectively, in Fig. 8B_lower).

Taken together, these data indicated that IR drift variance as well as Tdv are useful for visualizing the pre-symptomatic state of klotho mice as well as the physiological recovery by the administration of JTT as a whole.

## Discussion

### How much did the administration of JTT restore the physiological state of the klotho mice?

There are several lines of evidence that JTT administration restored the physiological state of klotho mice, although perhaps not completely. First, as described in the previous section, JTT administration reduced the values of the transcriptome and IR drift variances in the liver of klotho mice. Considering the definition of Tdv (Rangaraju et al.; 2015), this inferred Tdv data is the best indicator of the effect of JTT. To investigate the restorative effect of JTT on another organ of klotho mice, we also applied this method to the transcriptome of the kidney of klotho mice. The data show that JTT indeed has the restorative effect on the kidney, as the value of Tdv is significantly decreased in KL‒ (Fig. S15). Second, here are the data of metabolomic analysis. Initially, it was shown that the mice were under pseudo-starvation conditions because their 3-hydroxybuthylic acid (ketone bodies) was elevated and it was observed that JTT administration slightly, but not significantly, reduced these ketones (Okada et al. 2021, 2022). This seems to indicate a slight improvement in the energy status of the liver cells. Indeed, for Pck2, one of the major enzymes involved in gluconeogenesis, we observed that JTT treatment restored its expression to wild-type levels (Fig. 3D). Third, we have previously reported that JTT administration promotes heme synthesis and further mitochondrial recovery (Okada et al., 2022). Heme is synthesized in eight steps using glycine and succinyl-CoA as starting materials, and four of the eight enzymes were found to be 2- to 5-fold more transcribed by JTT (Okada et al., 2022). Since heme is known to be a cofactor for many mitochondrial enzymes (Kim et al., 2012), this may also lead to mitochondrial activation (Okada et al. 2022). Indeed, several enzymes of the mitochondrial complex V (e.g., Atp5a1, Atp5b, Atp5d, etc.) whose activity was reduced by KL‒ were found to be restored by JTT (Fig. S16). Fourth, as discussed in the previous section, the γ subunits of AMPK (*Prkag1* and *Prkag2*) were activated in KL‒, indicative of the pseudo-starvation state of klotho mice (Fig.2B). Notably, these two subunits were restored to the WT levels by JTT administration (Fig.2B). Considering the importance of this enzyme for the overall energy status of an organ, this may be excellent evidence for the physiological recovery of klotho mice. In addition, among seven genes of AMPK signaling enriched in the A-type (Fig. 6A), all the patterns of transcripts and IR except those for Pfkfb2 show the recovery pattern of these transcription and IR after JTT administration (Fig. 6BC). Fifth, as detailed in the paper (Okada et al. 2021), gene expression corresponding to metabolism-related enzymes specifically expressed in the liver was greatly improved by administration of JTT. Since the regulation of metabolism-related enzymes such as sugars, cholesterol, lipids, and amino acids in the liver is known to occur at the transcriptional level (Desvergne et al., 2006), this suggests that these metabolites are indeed restored by JTT administration compared to KL‒.

Taken together, these data suggest an actual restorative effect of JTT on klotho mice, but whether or not physiological status is actually restored would need to be confirmed by behavioral testing.

### A linkage of the cellular energy status to IR

When the body is in a state of starvation, the hypothalamus detects the low levels of both cellular energy and nutrients. This leads to the production of corticotropin-releasing hormone. Corticotropin-releasing hormone signals the anterior pituitary gland to produce adrenocorticotropic hormone and release it into the bloodstream where it travels to the adrenal glands, causing the release of glucocorticoids. This stress-induced pathway is called the HPA axis (Kudielka and Kirschbaum, 2005). Glucocorticoids then stimulate the breakdown of protein and fat to produce glucose through gluconeogenesis, which provides critical energy in a starvation-like state (Hatting et al., 2018; Fig. 3A & 8C). Starvation has systemic effects as described above, and the AMPK pathway is activated in every cell of an organ (Fig. 2A & 3A). Indeed, the AMPK pathway was activated in the lung, brain and liver of klotho mice (unpublished data). When the cellular physiological conditions induced by a starvation-like state were restored to some extent in response to JTT medicine (Fig. 8A & B), this information was apparently transmitted back to induce AMPK inhibition by an unknown factor(s) (Factor X in Fig. 8C), which promoted the reversion of IR to that of a WT state. This suggests that a link between cellular energy homeostasis (e.g., glucose concentration) and the regulation of IR during pre-mRNA splicing. This scenario is reminiscent of a very recent publication by Miao et al. (2023), who demonstrated that glucose binds the RNA helicase DDX21 to modulate its role in splicing and that this process promotes epidermal differentiation; this is the first report of a link between glucose concentration and IR. However, we speculate that there must be a similar but distinct pathway that further links glucose and IR to the cellular energy state as represented by the activity of the AMPK pathway, rather than to the state of cellular differentiation, as we visualized it by the drift variation analysis as shown in Fig. 8B & C.

Taken together, the present study suggests the existence of a mechanism that measures the appropriate amount of a protein in the cytoplasm and relays this information to the nucleus, where the amount of the pre-mRNA is adjusted by controlling splicing activity. In other words, there must be a mechanism by which a splicing complex with an intron-containing pre-mRNA is targeted for intron removal due to the activity of a certain factor that is sensitive to the physiological state of a cell.

### IR is linked to the AMPK/mTOR signaling pathway through starvation during evolution

Using the budding yeast *Saccharomyces cerevisiae* as a model, Parenteau et al. (2019) extended their previous work on introns in ribosomal protein genes (Parenteau et al. 2011) to show that the physical presence of introns in the genome promotes cell survival under starvation conditions regardless of the function or expression of the intron’s host gene. Transcriptomic and genetic analyses revealed that introns promote starvation resistance by increasing the expression of ribosomal protein genes downstream of the nutrient-sensing TORC1. On the other hand, by using the same model species, Morgan et al. (2019) reported another interesting study linking introns and TOR. They discovered that excised linear introns from 34 genomic loci in budding yeast stably accumulate under stress conditions where TOR signaling is repressed such as by prolonged rapamycin treatment. These introns differ from other introns in that the distance between their lariat branch point and the 3’ splice site is short. The authors showed that this type of intron structure is necessary and sufficient for intron stabilization and function. They demonstrated that the excised introns have a functional significance for cell growth in the stationary nutrient-depletion condition. These two yeast papers clearly showed that introns in the genome are indispensable for survival under stress condition including starvation.

From a cell physiological point of view, the strategy against starvation and other stresses has evolved to be controlled by the AMPK/TOR signaling pathway in the form of suppression of anabolism and promotion of catabolism. The above work in yeast suggests that the presence of introns in genes has evolved as a survival strategy against starvation, thus linking starvation to IR. Our results suggest that, in mammals, there is a potential link between starvation activated by AMPK signaling and IR. The coincidence between the involvement of short introns (although short introns are not necessarily those with a short distance between their lariat branch point and the 3’ splice site) in survival under stress in budding yeast (Morgan et al.,2019) and several short introns’ involvement in longevity (Fig. 7F) in klotho mice suggests that some functional short introns have been conserved during evolution.

## Conclusion

Experiments with klotho mice revealed that most IRs are caused by starvation stress and may be involved in fine-tuning the stress-induced activation or repression of certain genes. The introns involved in such fine-tuning were characterized as relatively short and were enriched in GC content. Since some of the IR events returned to a healthy state when nutritional status was restored by administration of JTT, we propose that cells have a potential mechanism that senses such nutritional status and links it to an IR.

## CRediT authorship contribution statement

Norihiro Okada: conceptualization, project administration, supervision, and writing of the manuscript. Kenshiro Oshima: data curation, formal analysis, and writing of the Methods text. Akiko Maruko: data curation and formal analysis. Trieu-Duc Vu: formal analysis, data validation and discussions. Ming-Tzu Chiu: formal analysis, data validation and discussions. Mitsue Nishiyama, Masahiro Yamamoto, and Akinori Nishi: data validation and discussions. Yoshinori Kobayashi: data validation, discussions and supervision.

## Acknowledgement

We thank professors Akila Mayeda, Kazuhiro Fukumura and Naoyuki Kataoka for useful discussions. We also thank Ms. Kyoko Yamada for drawing illustrations.

