## Supplementary material for "A potential pathway that links intron retention with the physiological recovery by a Japanese herbal medicine": Table S1 - S4

Table S1. Enrichment analysis of 4,737 up-regulated genes between WT and KL– in the 1st rep replicate

| **Category** | **Term** | **Count** | **PValue** | **Genes** |
| --- | --- | --- | --- | --- |
| GOBP | protein phosphorylation | 207 | 2.26E-17 | *Stk26, Stk19, Map3k12, Map3k6, Speg, Ephb4, Grk3, Trim24, Brsk1, Irak3, Map4k2, Grk6, Ntrk1, Map4k1, Blk, Nek3, Pim3, Ccl3, Ksr2, Limk2, Cdk18, Mapk7, Cdk9, Rps6kl1, Map3k5, Phkg2, Fes, Clk3, Gas6, Epha7, Ntrk3, Brd4, Pkn3, Ikbke, Map4k3, Erbb2, Araf, Stk38, Cdk5, Obscn, Ulk3, Alpk3, Csf2rb2, Cdk10, Camk2b, Rorc, Birc5, Rnasel, Nuak2, Mastl, Mapk12, Map3k8, Fastk, Strada, Trp53rka, Prkcb, Haspin, Fgfr1, Csf2ra, Cdc25b, Cit, Clk4, Ulk1, Prkab1, Pdgfrb, Cdk8, Cdkl3, Map3k14, Nek8, Plk1, Mink1, Mast3, Cdk1, Kit, Taf4b, Chek1, Btk, Mast2, Stkld1, Csnk1e, Hsf1, Prkd2, Grk2, Prrt1, Tex14, Bmx, Mst1r, Sik1, Aatk, Epha1, Ccnt2, Csnk1g2, Syk, Trpm6, Mark2, Bub1b, Cdk11b, Pak4, Mark4, Prkx, Cdc7, Camk1, Dclk2, Fgr, Taf8, Igf1r, Rps6kb2, Map2k7, Lck, Mertk, Camk2d, Jak3, Mapk11, Limk1, Nek5, Stat4, Matk, Incenp, Clk2, Eif2ak4, Abl1, Npr1, Map2k6, Pak6, Ilk, Slk, Trpm7, Cdk16, Clk1, Prkag1, Gucy2c, Stk25, Prkcd, Smg1, Pak1, Txk, Nrbp2, Plk3, Phka2, Bmp2, Ripk3, Trim28, Prkar1b, Cdc42bpg, Trib2, Dmpk, Flt3l, Morc3, Map3k21, Fyn, Srpk2, Prkag3, Pan3, Stk10, Flt1, Cdk20, Itk, Hcst, Trp53rkb, Lrrk1, Sox9, Klhl3, Map3k11, Pdgfra, Alpk1, Prkag2, Camkk1, Mknk1, Ngf, Ksr1, Atr, Stk32c, Mok, Map4k4, Grk4, Ddr1, Mapk13, Pick1, P2rx7, Aurkb, Mylk, Ephb3, Sik3, Cdkl4, Map3k15, Spn, Stk39, Mapkapk5, Taok2, Trio, Mark1, Stk11, Phkg1, Rara, Ikbkb, Wnk2, Ilf3, Mknk2, Ripk4, Tyk2, Pdgfb, Tssk1, Acvr2b, Pkmyt1, Tbp, Prkcg, Slc12a3* |
| GOBP | positive regulation of transcription from RNA polymerase II promoter | 337 | 1.01E-14 | *Gps2, Ppp1r12a, Efcab7, Mzf1, Dhx36, Tead1, Lpin3, Rhoq, Rrp1b, Spi1, Ptprn, Crebbp, F2rl1, Mybl1, Hes1, Mcrs1, Zfp263, Thap3, Bloc1s2, Ttc5, Mamstr, Arid3b, Mtf2, Hamp, Erg, Slc9a1, Ldb1, Tcerg1, Ccl3, Hdac4, Tead3, Arhgef2, Nampt, Nfkb2, Mapk7, Patz1, Cdk9, Crtc1, E4f1, Sox17, Fli1, Trerf1, Rnf40, Zfp410, Tstd1, Arid5a, Nr2f1, Elf4, Fosl2, Ppp3ca, Ppargc1b, Epc1, Agrn, Senp1, Brd4, Supt20, Dot1l, Ntf3, Hey1, Vegfa, Six5, Smarca2, Brca1, Kdm6b, Adrb2, Nr1h5, Scx, Kat5, Zfp53, Rgma, Hjv, Tead4, Mcf2l, Supt5, Rnasel, Camta2, Paf1, Zbtb49, Csrnp2, Hand2, Paxbp1, Tal1, Kat6b, Prkn, Zfp212, Pelp1, Nfyc, Fgfr1, Tbx2, Usf2, Pogz, Pou2f2, Ccdc62, Irf8, Tcf7l2, Nfatc4, Med10, Ssbp4, Rad54l2, Bcl9l, Mlxip, Pik3r2, Zfp384, Zscan21, Kansl3, Ctbp2, Nlrc5, Sertad1, Lpin2, Dpf3, Nr1i3, Zfp677, Preb, Foxm1, Hsf1, Prkd2, Cd40, Hoxb4, Med25, Zfp715, Dvl1, Hinfp, Prdm9, Myo1c, Glis3, Rbmx, Dcn, Tfeb, Sp100, Tsc2, Nfya, Ccnt2, Ncoa3, Sirt7, Klf6, Rarg, Junb, Mybl2, Med17, Wwp2, Kdm1a, Zfp395, Trp53bp1, Ppargc1a, Gtf2a2, Klf13, Camk1, Skap1, Dcaf6, Foxf1, Nfatc2, Kmt2a, Lpin1, Dvl3, Peg3, Hoxa1, Zfp639, Zmiz2, Ikzf3, Esrra, Ahr, Maml2, Jak3, Ctcfl, Top2a, Mafg, Ski, Fgf2, Kat2a, Maf, Tcf7l1, Mllt6, Auts2, Tfec, Gmeb2, Stat4, Atad2, Klf1, Mmp12, Gmeb1, Crlf3, Crtc2, Nufip1, Mybbp1a, Stox2, Caprin2, Sbno2, Nr1i2, Ikzf1, Irf1, Twist1, Ahi1, Med12, Zfp219, Bclaf1, Sf3b1, Thrap3, Zfp646, Meis3, Ier5, Zfp451, Sox12, Bcl9, Zbed4, Hsf4, Ier2, Txk, Meis1, Spic, Bmp2, Fos, Tcea2, Tcf3, Myc, Maff, Tnf, Rapgef3, Spag8, Flt3l, Ncoa7, Irx3, Arid4b, Prdm15, Arid2, Foxj2, Was, Klf4, Jmjd6, Prdm4, Sox4, Chd6, Cd28, Zfp143, Relb, Rbck1, Zfp142, Pparg, Carf, Eng, Sox9, Akap8l, Hnrnpd, Baz1b, Foxa2, Tbx20, Mrtfa, Ablim2, Lmo4, Kmt2d, Safb, Sfpq, Tet2, Prdm5, Usf1, Zfp523, Dab2ip, Grhl2, Ubp1, Chd7, Hmgb2, Zfp467, Nkx2-6, Pbx1, Hivep2, Churc1, Akna, Cxcr3, Fhod1, Hey2, Nfkbiz, Dll1, Zfp335, Arrb1, Deaf1, Dab2, Hdac3, Brd8, Fbln5, Pprc1, Per1, Nr1h2, Zc3h12a, Runx1, Tfap4, Gata6, Eaf2, Chd8, Sox18, Csrnp1, Tbx6, Iqce, Flcn, Glis2, Sall1, Tnip2, Irf3, Ifi207, Klf2, Jag1, Carm1, Asxl1, Zfp658, Ercc6, Zbtb17, Tef, Mllt10, Atf3, Rfx2, Dvl2, Setd4, Bptf, Rara, Bcl2l12, Ikbkb, Esr1, Cebpd, Gata3, Tfcp2, Egr2, Pygo1, Jun, Slc11a1, Pou2af1, Zfp827, Zfp746, Lhx2, Ap3d1, Gata2, Ddit3, Daxx, Hltf, Bclaf3, Maz, Mef2a, Atoh8, Pkd1, Tead2* |
| GOBP | phosphorylation | 193 | 1.33E-14 | *Stk26, Stk19, Map3k12, Map3k6, Speg, Ephb4, Grk3, Dgka, Adpgk, Brsk1, Map4k2, Grk6, Ntrk1, Map4k1, Blk, Nek3, Ckb, Tk2, Pim3, Ksr2, Limk2, Cdk18, Mapk7, Cdk9, Rps6kl1, Map3k5, Riok1, Phkg2, Fes, Clk3, Uckl1, Epha7, Ntrk3, Pkn3, Ikbke, Map4k3, Erbb2, Araf, Tk1, Stk38, Pck1, Dgke, Cdk5, Obscn, Ulk3, Alpk3, Cdk10, Camk2b, Nuak2, Mastl, Mapk12, Map3k8, Fastk, Chkb, Prkcb, Haspin, Fgfr1, Pfkfb1, Cit, Clk4, Ulk1, Pdgfrb, Cdk8, Cdkl3, Map3k14, Nek8, Plk1, Mink1, Mast3, Cdk1, Kit, Chek1, Btk, Ip6k1, Mast2, Csnk1e, Prkd2, Grk2, Ip6k2, Bmx, Mst1r, Sik1, Aatk, Epha1, Csnk1g2, Syk, Trpm6, Mark2, Bub1b, Cdk11b, Pak4, Mark4, Prkx, Cdc7, Camk1, Dclk2, Fgr, Igf1r, Pi4ka, Rps6kb2, Lck, Mertk, Camk2d, Jak3, Mapk11, Limk1, Nek5, Pip5k1c, Matk, Clk2, Hk3, Eif2ak4, Abl1, Map2k6, Dgkq, Pak6, Nmrk1, Ilk, Slk, Trpm7, Cdk16, Clk1, Stk25, Sgms1, Prkcd, Smg1, Pak1, Txk, Plk3, Ripk3, Cdc42bpg, Dmpk, Cerk, Map3k21, Fyn, Sphk1, Srpk2, Xylb, Stk10, Flt1, Cdk20, Itk, Lrrk1, Baz1b, Map3k11, Pdgfra, Itpka, Alpk1, Coq8b, Camkk1, Mknk1, Etnk2, Ksr1, Atr, Pi4kb, Pik3c2g, Taf1, Stk32c, Mok, Map4k4, Grk4, Ddr1, Mapk13, Shpk, Aurkb, Mylk, Ephb3, Sik3, Cdkl4, Map3k15, Stk39, Mapkapk5, Pik3cd, Hkdc1, Taok2, Trio, Mark1, Stk11, Phkg1, Galk1, Ikbkb, Wnk2, Mknk2, Ripk4, Tyk2, Dgkz, Tssk1, Gk5, Pip4k2a, Pfkfb4, Acvr2b, Pkmyt1, Prkcg* |
| GOBP | intracellular signal transduction | 152 | 2.75E-14 | *Tns1, Dnmbp, Inpp5d, Dgka, Impact, Npr1, Zfp36, Brsk1, Irak3, Plcb2, Map4k2, Dgkq, Neurl2, Gmip, Map4k1, Arhgef7, Pak6, Dcdc2b, Nrg4, Nrgn, Tiam1, Gucy2c, Arhgef2, Asb1, Ksr2, Mapk7, Adcy6, Sh2b3, Myo9a, Cblb, Prkcd, Plcd1, Prex1, Pak1, Ccn2, Rasa3, Nrbp2, Rab40c, Pkn3, Map4k3, Erbb2, Araf, Adcy4, Stk38, Mdk, Dgke, Cdc42bpg, Rapgef3, Stac3, Dmpk, Gucy1a1, Fyn, Mcf2l, Unc13a, Srpk2, Sh2b2, Nlrc3, Rgs14, Itk, Nuak2, Mastl, Mapk12, Lrrk1, Myzap, Prkcb, Haspin, Rasa2, Wsb1, Dcdc2a, Ccdc68, Rgs11, Cit, Racgap1, Prkag2, Pdgfrb, Adcy5, Camkk1, Mknk1, Socs2, Ksr1, Rapgef5, Plek2, Mink1, Mast3, Smpd2, Stk32c, Lat, Mok, Kit, Map4k4, Chek1, Nlrc5, Btk, Socs3, Mapk13, Prex2, Mast2, Pick1, Prkd2, Adcy3, Tns2, Plcd3, Nod1, Dvl1, Sh2b1, Sik3, Rassf1, Bmx, Stk39, Mapkapk5, Arhgef6, Sik1, Taok2, Myo9b, Nfam1, Plcb3, Rasa4, Mark1, Dok1, Syk, Dvl2, Nrg1, Mark2, Adcy7, Nlrx1, Plcg1, Pak4, Arhgap45, Mark4, Wnk2, Mknk2, Tyk2, Smad7, Dclk2, Dgkz, Dvl3, Hmox1, Lat2, Tssk1, Bax, Rasal1, Rgs9, Asb6, Dab1, Lck, Jak3, Mapk11, Prkcg, Abr, Lax1, Cish, Vav1* |
| GOBP | cellular response to DNA damage stimulus | 168 | 3.86E-14 | *Sirt6, Tdp2, Dclre1a, Mdc1, Ercc5, Xab2, Palb2, Abl1, Rad9a, Gnl1, Brsk1, Mapt, Uimc1, Smc5, Zranb3, Mcrs1, Suv39h1, Timeless, Xrcc1, Eme2, Blk, Poli, Rad18, Ttc5, Apc, Chd2, Bclaf1, Slx1b, Brca2, Cdk9, Alkbh3, Pidd1, Fanci, Prkcd, Rbbp6, Ap5z1, Helq, Smg1, Sprtn, Topbp1, Pak1, Usp28, Polq, Plk3, Slx4, Mcm8, Brd4, Ikbke, Cfap410, Mutyh, Lig1, Ier3, Susd6, Xpa, Mpg, Pmaip1, Myc, Ccnk, Brca1, Otub1, Nfrkb, Kat5, Zbtb7a, Pold1, Stub1, Acer2, Ercc2, Phf1, Mastl, Paxip1, Pwwp3a, Rhno1, Atad5, Kin, Baz1b, Ddb2, Fanca, Faap20, Pole, Tdg, Emsy, Cep63, Fancm, Rev1, Pogz, Rad50, Parp10, Rbbp8, Ercc6l2, Hmces, Sfpq, Mms19, Slf1, Btg2, Nupr1l, Hdgfl2, Uhrf1, Atr, Chaf1a, Fancd2, Abraxas1, Primpol, Rad51ap1, Fbxo31, Morc2a, Chek1, Uba7, Hdac10, Faap100, Bard1, Rnf169, Spindoc, Adprs, Fancc, Brat1, Pold4, Dclre1b, Alkbh2, Wrap53, Apex2, Zc3h12a, Flywch1, Foxm1, Hsf1, Ascc2, Ctc1, Eya4, Actr5, Rassf1, Rad52, Traip, Cgas, Chd1l, Taok2, Setd1a, Ercc6, Stk11, Rad51b, Tfpt, Sirt7, Ercc4, Chaf1b, Pold3, Uvssa, Parp2, Rtel1, Ccar2, Mbd4, Bcl2, Trp53bp1, Msh5, Rad54b, Nfatc2, Ppp4r3a, Xrcc3, Rnf8, Zbtb40, Atrip, Polm, Cep164, Bax, Fzr1, Fbxo5, Pms2, Gen1, Top2a, Aim2, Ddx11* |
| GOBP | chromatin organization | 127 | 8.83E-14 | *Sirt6, Arid1a, Atad2, Crebbp, Tada2a, Uimc1, Mcrs1, Tox, Suv39h1, Alkbh4, Pcgf2, Hmg20a, Ikzf1, Phf21a, Samd1, Mtf2, Chd2, Lrwd1, Hdac4, Hdac7, Kansl2, Baz2a, Zzz3, Rnf40, Rccd1, Kdm3b, Cbx4, Epc1, Epop, Brd4, Trim28, Cbx2, Cbx8, Msl1, Smarca2, Kdm6b, Tspyl2, Prmt7, Kat5, Zbtb7a, Arid4b, Brd9, Arid2, Phf1, Jmjd6, Kmt2e, Chd6, Hdac6, Kmt2b, Hmgn2, Pwwp3a, 0610010K14Rik, Kat6b, Usp36, Baz1b, Foxa2, Phf20, Tdg, Prkcb, Emsy, Smarcd2, Haspin, Kdm2b, Kmt2d, Parp10, Tet2, Prdm5, Chd7, Banp, Smarcd1, Uhrf1, Abraxas1, Nfkbiz, Kansl3, Cdan1, Hdac10, Kmt5c, Phf19, Ep400, Hdac3, Brpf3, Brd8, Brpf1, Dpf3, Mrgbp, Snai2, Fendrr, Bap1, Satb1, Eya4, Chd8, Prdm9, Ftx, Usp21, Carm1, Asxl1, Kdm4b, Setdb2, Bahd1, Macroh2a2, Setd1a, Ing2, Ring1, Cbx6, Brd1, Sirt7, Hira, Prmt1, Usp49, Ehmt2, Kdm1a, Atxn7l3, Setmar, Bag6, Kmt2a, Rnf8, Hmg20b, Tdrd3, Daxx, Hltf, Ctcfl, 2810410L24Rik, Trrap, Cbx7, Kat2a, Setdb1, Mllt6* |
| GOBP | DNA repair | 131 | 5.19E-13 | *Sirt6, Tdp2, Dclre1a, Mdc1, Ercc5, Xab2, Palb2, Abl1, Rad9a, Uimc1, Smc5, Zfp668, Zranb3, Mcrs1, Wdhd1, Timeless, Xrcc1, Eme2, Poli, Rad18, Ttc5, Recql4, Slx1b, Brca2, Cdk9, Alkbh3, Fanci, Ap5z1, Helq, Smg1, Sprtn, Topbp1, Usp28, Alkbh1, Polq, Slx4, Mcm8, Mutyh, Lig1, Trim28, Dot1l, Xpa, Mpg, Brca1, Otub1, Endov, Nfrkb, Kat5, Zbtb7a, Pold1, Stub1, Ercc2, Paxip1, Pwwp3a, Kin, Ddb2, Fanca, Faap20, Pole, Tdg, Emsy, Fancm, Rev1, Pogz, Rad50, Parp10, Rbbp8, Ercc6l2, Sfpq, Chrna4, Mms19, Slf1, Hdgfl2, Uhrf1, Atr, Chaf1a, Fancd2, Abraxas1, Primpol, Rad51ap1, Chek1, Hdac10, Kmt5c, Faap100, Bard1, Rnf169, Ep400, Adprs, Fancc, Pold4, Dclre1b, Alkbh2, Wrap53, Apex2, Foxm1, Hsf1, Ascc2, Eya4, Actr5, Hinfp, Rad52, Traip, Cgas, Chd1l, Ercc6, Rad51b, Tfpt, Sirt7, Ercc4, Chaf1b, Pold3, Uvssa, Parp2, Rtel1, Mbd4, Rad9b, Trp53bp1, Msh5, Upf1, Xrcc3, Rnf8, Atrip, Polm, Cep164, Fzr1, Hltf, Pms2, Gen1, Trrap, Rpain, Ddx11* |
| GOBP | cell cycle | 191 | 3.96E-12 | *Cdc20, Smc4, Urgcp, Ambra1, Dclre1a, Mdc1, Mdm2, Gmnn, Rassf4, Bin3, Incenp, Crocc, Eif2ak4, Epb41, Mtbp, Brsk1, Mis18bp1, Hjurp, Smc5, Arl2, Klhdc8b, Suv39h1, Timeless, Ikzf1, Nek3, Prc1, Bora, Ccne2, Cntrl, Pim3, Arhgef2, Brca2, Mapk7, Cdca2, E4f1, Bex2, Smpd3, Ncapd2, Fanci, Prkcd, Ncapd3, Dctn1, Eml3, Ccsap, Plk3, Gadd45gip1, Cntrob, Mcm8, Haus3, Lig1, Dsn1, Ccnk, Ube2i, Brca1, Cdk5, Spag8, Ube2c, Syce2, Cep250, Tspyl2, Anapc15, Mau2, Ccnd3, Tacc3, Kmt2e, Birc5, Usp2, Rgs14, Stk10, Cdk20, Ppp2r2d, Ing1, Mastl, Cenpe, Zbtb49, Map3k8, Mapk12, Rhno1, Tpx2, Sgo1, Chtf18, Spdl1, Ist1, Mis18a, Cables2, Strada, Epb41l2, Cdkn1c, Haspin, Cdc14a, Cep63, Reep4, Zwilch, Pogz, Rcc2, Knstrn, Anapc5, Haus5, Cdc25b, Rad50, Rbbp8, Cit, Tet2, Dab2ip, Appl2, Racgap1, Banp, Rassf2, Mad2l1, Son, Nupr1l, Rab11fip3, Uhrf1, Entr1, Chaf1a, Plk1, Fancd2, Siah1a, Taf1, Fbxo31, Cdk1, Chek1, Mapk13, Trp53bp2, Ska2, Rcc1, Mcm6, Smc1b, Apex2, Foxm1, Aurkb, Cep131, Lzts2, Cenpt, Ttc28, Katnb1, Tex14, Rassf1, Gspt2, Lzts1, Sik1, Lrrcc1, Haus7, Anapc4, Setdb2, Clasp2, Nsl1, Kif2a, Stk11, Ccnt2, Cd2ap, Septin4, Ncapg2, Sgsm3, Ccna2, Chaf1b, Bub1b, Pak4, Cdk11b, Ccar2, Septin6, Aspm, Mark4, Bcl2l11, Cdc7, Camk1, Septin1, Kif20b, Ccar1, Hells, Inca1, Ccno, Hmcn1, Rnf8, Nusap1, Haus1, Hmg20b, Cep164, Fzr1, Ddit3, Cdca3, Fbxo5, Ahr, Ska3, Spice1, Terf2, Pkmyt1, Ccnb2, Snx33, Mki67, Anln* |
| GOBP | regulation of transcription from RNA polymerase II promoter | 403 | 6.61E-12 | *Gps2, Ambra1, Spi1, Zfp213, Zfp433, 9130019O22Rik, Klf16, Zfp865, Zfp628, Mamstr, Arid3b, Erf, Ldb1, Erg, Ppp1r13l, Tead3, Hdgfl3, Nfkb2, E4f1, Zfp184, Zfp90, Sp110, Kdm3b, Zfp653, Fbxl19, Arid5a, Nr2f1, Epc1, Zfp945, Supt20, Six5, Scx, Lyl1, Zbtb7a, Zfp472, Zkscan4, Gm14443, Mnt, A630001G21Rik, Zfp579, Supt5, Zbtb48, Camta2, Tal1, Zfp212, Prkcb, Tbx2, Usf2, Irf8, Zfp276, Gm5141, Nupr1l, Mlxip, Zbtb24, Zfp384, Zscan21, Ctbp2, Wiz, Zfp28, Zfp683, Brpf3, Zfp677, Mrgbp, Rbak, Phf14, Hoxb4, Rfx1, Zfp57, Zfp113, Glis3, Atp2b4, Sp100, Ccnt2, Zfp229, Snai1, Sirt7, Klf6, Junb, Zfp777, Med17, Platr25, Zfp558, Zfp934, Zfp608, Zfp808, Hivep3, Foxf1, Nfatc2, Peg3, Hoxa1, Zfp639, Tsc22d4, Ctcfl, Zscan2, Mafg, Zfp950, Kat2a, Mllt6, Gmeb2, Stat4, Mamld1, Klf1, Mdm2, Zbtb45, Gmeb1, Zfp9, Zfp668, Phf20l1, Zfp553, Pias4, Dgkq, Twist1, Med12, Zfp651, Zfp97, 9130023H24Rik, Zfp61, Mbip, Zfp956, Zfp623, Zfp62, Zfp787, Hsf4, Smad6, Meis1, Fos, Myc, Tcf3, Tnf, Ccnk, Zfp40, Zfp423, Zscan26, 5430403G16Rik, Med4, Arid2, Prdm4, Relb, 2610008E11Rik, Zfp672, Zfp202, Pparg, Zfp316, Havcr2, Zfp740, 2610044O15Rik8, Sox9, Zbtb3, Mrtfa, Hdx, Zfp458, Prdm5, Zfp523, Usf1, Grhl2, Smarcd1, Zfp467, Nkx2-6, Hivep2, Cc2d1a, Med20, Zbtb25, Med15, Arrb1, Zfp420, Deaf1, Zfp317, Zfp366, Batf3, Pias2, Nr1h2, 2610021A01Rik, Zfp691, Tfap4, Gata6, Glis2, Tnip2, Klf2, Zfp658, Brd1, Tef, Mllt10, Atf3, Rfx2, Zscan25, Cebpd, Ehmt2, Gata3, Egr2, Atxn7l3, Smad7, Lhx2, Zfp746, Zbtb40, Ddit3, Med18, Ccnl1, Trrap, Atoh8, Zfp157, Zfp606, Zfp248, Zfp811, Tead1, Zfp629, Zfp764, Zfp36, Hes1, Tox, Zfp263, Sox7, Zfp408, Zfp791, Chd2, AU041133, Gm45871, Zfp952, Zfp689, Patz1, Sox17, Fli1, Trerf1, Zzz3, Zfp410, Elf4, Fosl2, Zfp457, Epop, E430018J23Rik, Hey1, Zfp236, Cux2, Vegfa, Smarca2, Brca1, Zfp641, Kat5, Zfp53, Brd9, Tead4, Mdm4, Rorc, Zfp747, Zbtb49, Csrnp2, Zfp444, Zfp281, Hand2, Nfyc, Zfp94, Pou2f2, Zfp788, Nfatc4, Tcf7l2, Jade2, Zfp935, Med10, Hdgfl2, Zfp566, Ccnl2, Taf4b, Zfp429, 2810021J22Rik, Zfp54, Brpf1, Dpf3, Zfp282, Foxm1, Hsf1, Zfp715, Hinfp, Zfp524, Tfeb, Foxp4, Nfya, Zfp354a, Msc, Rcor2, Tshz3, Zfp383, Zfp995, Kdm1a, Zfp395, Zscan18, Zfp275, Sltm, Klf13, Bicra, Zkscan17, Lpin1, Zfp69, Zmiz2, Zfp612, Zkscan3, Ikzf3, Ahr, Esrra, Zbtb16, Zfp59, Ptafr, Zfp7, Maf, Tcf7l1, Cc2d1b, Tfec, Arid1a, Pias3, Bcl7a, Zfp160, Ecm1, Zfp26, Tada2a, Med23, Stox2, Elf2, Zfp13, Ikzf1, Irf1, Zfp3, Meis3, Brwd1, Zfp869, Zfp786, Pcgf6, Cic, Zfp354c, Spic, Safb2, Trak2, Maff, Batf2, Ncoa7, Zfp268, Irx3, Zfp970, Arid4b, Zfp334, Prdm15, Foxj2, Klf4, Zbtb46, Zkscan5, Zfp143, Zfp362, Hmgn2, Carf, Zfp27, Zfp821, Foxa2, Phf20, Tbx20, Ddx5, Zfp580, Zfp146, Smarcd2, Kdm2b, Lmo4, Safb, Zfp37, Ncor2, Zfp58, Ubp1, Zfp607a, Zfp287, Zfp846, Prox2, Hmgb2, Pbx1, Zfp14, Hey2, Zfp446, Zfp12, Hic2, Zfp991, Zfp775, Runx1, Satb1, Zfp647, Csrnp1, Tbx6, Sall1, Zfp661, Irf3, Zfp52, Rai1, Zbtb17, Zfp772, Yeats2, Bptf, Zfp182, Esr1, Tfcp2, Jun, Pou6f1, Zfp46, Zfp948, Gata2, Myrf, Zfp1, Maz, Mef2a, Fus, Tead2* |
| GOBP | negative regulation of transcription from RNA polymerase II promoter | 266 | 6.26E-10 | *Zfp248, Gps2, Sirt6, Sap130, Mzf1, Thap7, Spi1, Crebbp, Zfp36, Hes1, Zfp263, Klf16, Hmg20a, Timeless, Mtf2, Erf, Hamp, AU041133, Ldb1, Tcerg1, Hdac4, Ppp1r13l, Gm45871, Zfp952, Patz1, E4f1, Zfp90, Taf9b, Sinhcaf, Cbx4, Arid5a, Pkig, Nr2f1, Epc1, Pcbp3, Hey1, Sla2, Cbx2, Cbx8, Cux2, Vegfa, Noc2l, Zfp641, Nr1h5, Phf12, Kat5, Zbtb7a, Ccnd3, Traf7, Hjv, Mdm4, Mnt, Rorc, Supt5, Paf1, Ing1, Per3, Zbtb49, Zfp281, Tal1, Prkn, Mier2, Cdkn1c, Dnajc17, Fgfr1, Tbx2, Ybx3, Kcnip3, Tro, Slfn1, Irf8, Nfx1, Tcf7l2, Nfatc4, Nupr1l, Hexim2, Zbtb24, Notch4, Zfp566, Plk1, Zfp384, 2810021J22Rik, Zfp282, Nr1i3, Snai2, Rbak, Flywch1, Foxm1, Hsf1, Phf14, Hoxb4, Med25, Zfp715, Hinfp, Zfp57, Sfn, Glis3, Sik1, Dll4, Sp100, Ing2, Foxp4, Zfp354a, Snai1, Sirt7, Msc, Rarg, Zfp777, Zfp383, Zfp558, Ift172, Wwp2, Kdm1a, Zfp608, Sorbs3, Eef1aknmt, Foxf1, Nfatc2, Mepce, Lpin1, Ighmbp2, Peg3, Tsc22d4, Zfp69, Cry2, Esrra, Ahr, Eid2, Zbtb16, Ski, Maf, Setdb1, Cc2d1b, Tfec, Arid1a, Pias3, Mdm2, Zbtb45, Mmp12, Impact, Zfp9, Zfp668, Pias4, Zfp13, Suv39h1, Pcgf2, Nr1i2, Ikzf1, Tcp10b, Twist1, Phf21a, Atn1, Samd1, Gbp4, Zfp219, Rpl10-ps3, Rbm10, Mta1, Hdac7, Zfp3, Ralgapa1, Sox12, Otud7b, Mdfi, Hsf4, Pcgf6, Cic, Plk3, Bmp2, Nrip2, Id1, Trim28, Tcf3, Myc, Tnf, Ube2i, Trpv4, Irx3, Morc3, Arid4b, Per2, Id3, Klf4, Sox4, Usp2, Hdac6, Zbtb46, Hmgn2, 2610008E11Rik, Zfp202, Pparg, Ddx20, 2610044O15Rik8, Eng, Sox9, Dnajb5, Tbx20, Ddx5, Tdg, Zfp146, Kdm2b, Lmo4, Sfpq, Prdm5, Dab2ip, Zfp37, Ncor2, Btg2, Hmgb2, Uhrf1, Cc2d1a, Zbtb25, Hey2, Deaf1, Hdac10, Cbfa2t3, Dab2, Phf19, Hdac3, Zfp366, Fbln5, Batf3, Tcf25, Trpv1, Per1, Nr1h2, Runx1, Aurkb, Gata6, Satb1, Chd8, Sox18, Tbx6, Flcn, Glis2, Sall1, Apbb3, Ifi207, Klf2, Macroh2a2, Zfp658, Cbx6, Zmym5, Yeats2, Atf3, Ripply3, Bptf, Rara, Ehmt2, Esr1, Gata3, Zc3h8, Jun, Smad7, Pou6f1, Zfp46, Zfp746, Gata2, Ddit3, Pcgf1, Acvr2b, Daxx, Maz, Mef2a, Cbx7, Zfp157* |
| GOBP | mRNA processing | 116 | 8.23E-10 | *Sfswap, Xab2, Zc3h13, Rrp1b, Celf4, Scaf8, Sf1, Yju2, Srsf7, Srrm4, Casc3, Prpf38b, Cpeb1, Sf3b1, Zcchc8, Tcerg1, Thrap3, Sf3a2, Tut1, Hnrnpa1, Rbm26, Sf3a1, Rbbp6, Srsf6, Nova2, Pus7l, Pan2, Gpatch1, Acin1, Rbfox3, Csdc2, Srsf5, Cd2bp2, Scaf4, Tra2a, Rnps1, Akap17b, Scaf1, Lgals3, Srpk2, Jmjd6, Pan3, Pdcd11, Mtpap, Rnasel, Rbm39, Slbp, Arl6ip4, Ddx20, Akap8l, Tra2b, Kin, Ddx5, Gcfc2, Snrpa1, Sugp1, Esrp2, Pus7, Sfpq, Adarb1, Dus3l, Tsen54, Son, Tia1, Trmt2a, Zrsr1, Ess2, Lsm5, Cpsf1, Nol3, Prpf39, Sart3, Cpsf7, Hnrnph1, Zpr1, Bud13, Ptbp2, Wdr83, Cpsf4, Smndc1, Ddx46, Prpf40b, Hsf1, Ddx39b, Snrpf, Srrt, Phrf1, Wbp11, Sart1, Scnm1, Rbmx, Cactin, Ddx39a, Zrsr2, Rbm5, Thoc1, Clasrp, Sympk, Rbm22, Hnrnpl, Rbm17, Srrm1, Srsf2, Usp49, Ythdc1, Ccar2, Lsm7, Srek1, Trub1, Rbm25, Srsf4, Lsm3, Tdrd3, Lsm8, Zfp326, U2af2* |
| GOBP | apoptotic process | 182 | 4.89E-09 | *Stk26, Dnase1, Shf, Tns4, Fbxo10, Zfp385b, Mdm2, Inpp5d, Ngb, Unc5a, Rrp1b, Palb2, Abl1, Mapt, Map2k6, Irak3, Rnf216, Card14, Ntn1, Plscr3, Cul7, Bik, Ift57, Irf1, Rhob, Apc, Slk, Bclaf1, Pim3, Siva1, Ppp1r13l, Pacs2, Dffb, Emp3, Dram1, Tchp, Map3k5, Pidd1, Sgms1, Bex2, Bex3, Sp110, Prkcd, Pglyrp1, Ncf1, Gadd45b, Pak1, Birc2, Plk3, Fap, Gas6, Tmem219, Dedd2, Epha7, Cdip1, Mfsd10, Id1, Acin1, Ripk3, Relt, Pmaip1, Vegfa, Noc2l, Elmo3, Cdk5, Kat5, Traf7, Kcnj8, Kitl, Ercc2, Bnip1, Birc5, Nuak2, Naip1, Csrnp2, Tpx2, Mul1, Hand2, Stil, Fastk, Tnfaip3, Sox9, Elmo2, Serpina3g, Fis1, Apaf1, Aldh1a3, Prkcb, Tbx2, Ybx3, Hip1, Gapdh, Kcnip3, S100a8, Dab2ip, Tia1, Elmo1, Nisch, Tnfsf12, Tnfaip8, Nkx2-6, Siah1a, Ecscr, Cdk1, Rassf7, Gramd4, Chek1, Btk, Bnip2, Dab2, Trp53bp2, Chil1, Brat1, Smndc1, Sav1, Htra2, Traf1, Zc3h12a, Tcirg1, Naip5, Ppif, Ltbr, Nod1, Prickle1, Csrnp1, Cd27, Tnip2, Bmx, Sema3a, Rbm5, Chac1, Aatk, Epb41l3, Thoc1, Stk11, Gadd45g, S100a9, Tfpt, Fhit, Rarg, G2e3, Marcks, Bub1b, Tnfrsf21, Tial1, Naip6, Bcl2l12, Pak4, Ccar2, Sema6a, Bcl2, Sltm, Hint2, Bcl2l11, Zc3h8, Jun, Mknk2, Bag6, Trim39, Ccar1, Map1s, Hells, Atg4d, Tnfrsf25, Rbm25, Casp12, Hmox1, Casp1, Peg3, Bax, Map2k7, Ddit3, Adrb1, Ifi27l2b, Daxx, Tnfrsf12a, Cd5l, Aim2, Mef2a, Prune2, Tradd, Adamtsl4* |
| GOBP | RNA splicing | 92 | 8.33E-09 | *Sugp1, Esrp2, Pus7, Sfswap, Sfpq, Xab2, Zc3h13, Rrp1b, Celf4, Son, Tia1, Zrsr1, Sf1, Yju2, Ess2, Lsm5, Prpf39, Sart3, Hnrnph1, Zpr1, Srsf7, Bud13, Casc3, Srrm4, Ptbp2, Prpf38b, Wdr83, Rbm10, Sf3b1, Zcchc8, Tcerg1, Smndc1, Thrap3, Sf3a2, Ddx46, Prpf40b, Hnrnpa1, Ddx39b, Snrpf, Sf3a1, Srsf6, Wbp11, Sart1, Rp9, Scnm1, Nova2, Rbmx, Cactin, Ddx39a, Ivns1abp, Zrsr2, Rbm5, Acin1, Rbfox3, Taf15, Thoc1, Clasrp, Srsf5, Cd2bp2, Ccdc130, Rbm22, Rbm17, Tra2a, Srrm1, Srsf2, Prmt1, Rnps1, Usp49, Akap17b, Ythdc1, Ccar2, Scaf1, Lsm7, Lgals3, Srek1, Srpk2, Jmjd6, Rbm39, Rbm25, Srsf4, Arl6ip4, Ddx20, Lsm3, Tra2b, Lsm8, Zfp326, Ddx5, Gcfc2, U2af2, Srsf11, Snrpa1, Fus* |
| GOBP | cell projection organization | 72 | 1.15E-08 | *Dcdc2a, Pcm1, Bbs4, Tmem138, Crocc, Ofd1, Dnajb13, Rsph4a, Entr1, Cep152, Wdr90, Bbs2, Ahi1, Prickle3, Ilk, Ttc12, Odf2l, Dync2li1, Spef1, Tchp, Cfap54, Ccdc66, Dzip1, Ccdc78, Cep43, Cep131, Iqcb1, Ccdc61, Myo7a, Ccdc88a, Rpgr, Cep89, Trappc14, Rab34, Cfap410, Ift43, Hap1, Tsc2, Macir, Tapt1, Ro60, Rfx2, Ptpdc1, Cep250, Ift88, Cep162, Rabep2, Mks1, Cluap1, Dync2h1, Bbs1, Mark4, Cdk10, Ocrl, Ccdc28b, Ccno, Tsc1, Poc1a, Cep164, Fhdc1, Camsap3, Tctn1, Tmem237, Cc2d2a, Ccp110, Ptpn23, Dync2i1, Enkd1, Flna, Lrrc56, Wtip, Cplane1* |
| GOBP | chromatin remodeling | 72 | 6.81E-08 | *Kdm2b, Arid1a, Klf1, Bcl7a, Sfpq, Phc1, Chd7, Tada2a, Smarcd1, Ttf1, Mybbp1a, Hdgfl2, Mcrs1, Pcgf2, Nfkbiz, Morc2a, Chek1, Kmt5c, Sf3b1, Hdac4, Brpf1, Mta1, Dpf3, Hdgfl3, Per1, Baz2a, Satb1, Baz1a, Chd8, Actr5, Myo1c, Pak1, Pcgf6, Chd1l, Brd4, Kdm4b, Ercc6, Ring1, Tcf3, Myc, Yeats2, Tfpt, Padi2, Msl1, Hira, Smarca2, Kdm6b, Setd4, Bptf, Nfrkb, Zbtb7a, Ino80e, Rere, Brd9, Per2, Arid2, Gata3, Scmh1, Chd6, Bicra, Kmt2b, Zfp827, Paxip1, Hmg20b, Pwwp2a, Pcgf1, Kat6b, Sox9, Daxx, Baz1b, Kat2a, Smarcd2* |
| GOBP | regulation of cell cycle | 93 | 7.40E-08 | *Slfn1, Sfpq, Mdm2, Adarb1, Dab2ip, Commd5, Tcf7l2, Eif2ak4, Jade2, Son, Abl1, Tada2a, Fbxl12, Mtbp, Crlf3, Mybbp1a, Nupr1l, Mcrs1, Inha, Irf1, Apc, Ccnl2, Ilk, Ccne2, Ep400, Brd8, Mrgbp, Cdk9, Foxm1, Mbip, Fbxl6, Zzz3, Evi2, Tfap4, Bap1, Gata6, Med25, Actr5, Sfn, Gadd45b, Rassf1, Fbxl8, Birc2, Fap, Epc1, Zfp703, Stk11, Tsc2, Zbtb17, Srsf5, Gadd45g, Myc, Yeats2, Tfpt, Sgsm3, Brca1, Junb, Sipa1, Smim22, Kat5, Nfrkb, Ccnd3, Ino80e, Tacc3, Cdk11b, Fignl1, Per2, Bcl2, Id3, Mdm4, Cdk10, Mnt, Kmt2e, Jun, Birc5, Setmar, Tsc1, Mastl, Zbtb49, Bax, Ddit3, Phactr4, Ccnl1, Cables2, Trim36, Trrap, Prdm11, Fgf2, Cdkn1c, Mif, Kat2a, Pkd1* |
| GOBP | positive regulation of cell migration | 85 | 1.30E-07 | *Sema4c, Pdgfra, Grn, Myo1f, Pdgfrb, Clec7a, F2rl1, Sema6b, Ccl24, Fam83h, Coro1a, Daam2, Pld2, Itgax, Carmil2, Twist1, Actg1, Kit, Apc, Aldoa, Map4k4, Ilk, Sema6d, Dab2, Cpeb1, Tiam1, Abcc1, Egf, Ccl3, Col18a1, Snai2, Spry2, Myadm, Plau, Mylk, Sema4b, Cep43, Atp8a1, Prex1, Plp1, Myo1c, Sema6c, Pak1, Pik3cd, Ppp3ca, Sema3a, Zfp703, Ntrk3, Hbegf, Bmp2, Epha1, Ntf3, Snai1, Mdk, Vegfa, Ube2i, Mmp14, Fermt3, Cd274, Itgb3, Gpnmb, Smim22, Csf1, Sema3b, Sema3f, Sema6a, Ripor1, Mcam, Vegfc, Sphk1, Foxf1, Ccar1, Flt1, Fgr, Pdgfb, Igf1r, Trip6, Ccr1, Arhgef39, Sema4g, Ager, Ccl5, Maz, Fam110c, Tradd* |
| GOBP | neural tube closure | 41 | 5.39E-07 | *Arid1a, Sema4c, Lmo4, Tulp3, Bbs4, Grhl2, Tsc2, Rarg, Abl1, Spint1, Scrib, Dvl2, Rara, Rgma, Mks1, Ift172, Shroom3, Cluap1, Ift57, St14, Twist1, Med12, Sec24b, Deaf1, Adm, Kif20b, Cdk20, Lhx2, Tsc1, Mthfd1l, Enah, Vangl2, Stil, Phactr4, Cc2d2a, Ski, Prickle1, Apaf1, Kat2a, Sall1, Tead2* |
| GOBP | protein autophosphorylation | 65 | 5.41E-07 | *Stk26, Map3k11, Map3k12, Pdgfra, Clk2, Clk4, Ephb4, Ulk1, Pdgfrb, Eif2ak4, Abl1, Trim24, Mknk1, Irak3, Atr, Mink1, Ntrk1, Map4k1, Kit, Btk, Slk, Trpm7, Clk1, Pim3, Ddr1, Stk25, Mapk7, Prkd2, Ephb3, Prkcd, Phka1, Fes, Clk3, Smg1, Pak1, Txk, Bmx, Stk39, Mapkapk5, Sik1, Brd4, Ripk3, Epha1, Erbb2, Stk11, Trim28, Csnk1g2, Syk, Cdk5, Obscn, Mark2, Ulk3, Fyn, Wnk2, Prkx, Camk2b, Mknk2, Stk10, Flt1, Fgr, Igf1r, Lck, Camk2d, Prkcg, Fgfr1* |
| KEGG | Herpes simplex virus 1 infection | 123 | 8.00E-07 | *Zfp248, Zfp811, Zfp160, Zfp764, H2-T10, Eif2ak4, Zfp26, Zfp9, Zfp433, Zfp13, 9130019O22Rik, Nxf1, Zfp791, Srsf7, AU041133, Gm45871, Zfp97, Zfp952, Zfp689, Zfp184, Zfp61, Zfp90, Zfp956, Zfp623, Zfp869, Zfp786, H2-Ab1, Srsf6, Birc2, Zfp457, Zfp354c, Ikbke, E430018J23Rik, Srsf5, Tnf, Zfp40, Itgb3, Zfp641, Zfp53, Zfp472, Zfp268, Zfp334, H2-Q4, Rnasel, Zfp747, Tsc1, 2610008E11Rik, Zfp316, Zfp27, Zfp212, H2-DMb1, Apaf1, Zfp94, Pou2f2, Zfp788, Tnfsf14, Zfp458, Zfp37, Gm32687, Zfp58, Gm5141, Zfp935, Zfp607a, Card9, Zfp566, Zfp14, Pik3r2, H2-DMa, Zfp420, Zfp12, Socs3, Zfp317, 2810021J22Rik, Zfp28, Zfp54, Zfp991, Zfp282, Zfp677, Rbak, Zfp715, Zfp647, Zfp57, Zfp113, Irf3, Cgas, Zfp661, Pik3cd, Zfp52, Zfp658, Sp100, Tsc2, Zfp772, Zfp229, Zfp354a, Syk, Zfp777, Zfp383, Srsf2, Zfp688, Zfp558, Zfp182, Ikbkb, Zfp808, Bcl2, Zfp275, Tyk2, Zfp46, Zfp746, H2-Eb1, Zfp948, Srsf4, Bax, Zfp69, Tab1, Zfp612, H2-Q6, Ccl5, Daxx, Zfp1, Zfp950, Zfp7, Tradd, Zfp157* |
| KEGG | Fanconi anemia pathway | 24 | 5.33E-06 | *Telo2, Slx4, Fancm, Rev1, Faap100, Fance, Slx1b, Fancc, Palb2, Atrip, Brca2, Ercc4, Top3b, Brca1, Hes1, Fanci, Top3a, Atr, Fanca, Pms2, Fancd2, Rmi1, Eme2, Poli* |
| KEGG | MAPK signaling pathway | 85 | 6.78E-06 | *Map3k11, Map3k12, Cdc25b, Pdgfra, Flnb, Map3k6, Pdgfrb, Mknk1, Ngf, Mapt, Map3k14, Map2k6, Rras, Map4k2, Ntrk1, Map4k1, Rasgrp4, Kit, Arrb1, Cacna1a, Map4k4, Il1r1, Hspa2, Mapk13, Egf, Dusp2, Vegfd, Mapk7, Nfkb2, Map3k5, Dusp7, Lamtor3, Gadd45b, Pak1, Mapkapk5, Ppp3ca, Mapk8ip3, Cacna1d, Taok2, Fos, Map4k3, Erbb2, Araf, Ntf3, Gadd45g, Cacnb1, Myc, Arrb2, Tnf, Vegfa, Vegfb, Gm5741, Flt3l, Rapgef2, Csf1, Rasgrp2, Ikbkb, Map3k21, Kitl, Dusp5, Vegfc, Cacnb3, Ppp3cc, Mknk2, Jun, Hspa1l, Flt1, Relb, Mapk8ip1, Pdgfb, Igf1r, Mapk12, Map3k8, Map2k7, Tab1, Ddit3, Daxx, Mapk11, Fgf2, Flna, Prkcg, Tradd, Prkcb, Rasa2, Fgfr1* |
| KEGG | Focal adhesion | 61 | 1.99E-05 | *Col9a3, Myl12b, Tln1, Ppp1r12a, Pip5k1c, Col4a5, Pdgfra, Flnb, Lamc3, Pdgfrb, Actn1, Pik3r2, Itga2b, Ppp1r12b, Pak6, Actg1, Zyx, Ilk, Egf, Vegfd, Itga7, Itga4, Spp1, Thbs2, Parvb, Mylk, Parvg, Col4a4, Col6a1, Tnxb, Pak1, Itgb7, Birc2, Pik3cd, Thbs3, Erbb2, Vegfa, Vegfb, Itgb3, Ccnd3, Lama5, Pak4, Lamb2, Bcl2, Fyn, Col4a3, Vegfc, Jun, Flt1, Pxn, Itgb8, Pdgfb, Vwf, Igf1r, Ppp1r12c, Itgb4, Flna, Prkcg, Prkcb, Rapgef1, Vav1* |
| KEGG | Osteoclast differentiation | 43 | 2.39E-05 | *Ncf2, Fosl2, Pik3cd, Ppp3ca, Fos, Lilrb4a, Trem2, Spi1, Tnf, Sirpb1b, Syk, Junb, Map3k14, Map2k6, Itgb3, Cyba, Pirb, Csf1, Pik3r2, Ikbkb, Ncf4, Fyn, Sirpa, Btk, Ppp3cc, Pira1, Il1r1, Jun, Socs3, Tyk2, Fcgr3, Nfatc2, Mapk13, Relb, Pparg, Mapk12, Nfkb2, Tab1, Map2k7, Lck, Mapk11, Sirpb1c, Ncf1* |
| KEGG | Pathways in cancer | 132 | 6.33E-05 | *Stat4, Gnb2, Mdm2, Lamc3, Gstt3, Spi1, Il3ra, Abl1, Crebbp, Mgst3, Nqo1, Hes1, Plcb2, Ntrk1, Lpar1, Apc, Gng7, Ccne2, Brca2, Nfkb2, Adcy6, Il15ra, Csf3r, Col4a4, Gadd45b, Gngt2, Ptch2, Gstp3, Birc2, Il7, Bmp2, Fos, Erbb2, Araf, Adcy4, Hey1, Pmaip1, Myc, Vegfa, Gm5741, Flt3l, Ccnd3, Rasgrp2, Lama5, Lamb2, Kitl, Csf2rb2, Camk2b, Birc5, Pparg, Ddb2, Apaf1, Prkcb, Fgfr1, Gm3776, Calml4, Csf2ra, Col4a5, Pdgfra, Jag2, Il7r, Tcf7l2, Epor, Pdgfrb, Adcy5, Arhgef1, Notch4, Il15, Pld2, Pik3r2, Hey2, Itga2b, Rasgrp4, Dll1, Plekhg5, Kit, Ctbp2, Egf, Vegfd, Gnb5, Traf1, Apc2, Runx1, Adcy3, Ptger4, Dvl1, Plcb4, Rassf1, Ptger2, Kng2, Gsta1, Pik3cd, Gstm1, Jag1, Slc2a1, Dll4, Plcb3, Zbtb17, Gadd45g, Gstm3, Ncoa3, Vegfb, Ccna2, Dvl2, Adcy7, Rara, Plcg1, Ikbkb, Gstt2, Esr1, Bcl2, Col4a3, Vegfc, Bcl2l11, Jun, Kif7, Pdgfb, Igf1r, Dvl3, Hmox1, Gnb4, Bax, Rps6kb2, Il6ra, Il2rb, Zbtb16, Camk2d, Jak3, Egln2, Fgf2, Prkcg, Tcf7l1* |
| KEGG | Axon guidance | 54 | 7.83E-05 | *Plxna3, Limk1, Myl12b, Sema4c, L1cam, Unc5a, Nfatc4, Ephb4, Dpysl2, Abl1, Ssh1, Sema6b, Rras, Ntn1, Pik3r2, Robo3, Ntng2, Pak6, Plxnb3, Ilk, Sema6d, Ssh3, Limk2, Enah, Sema4b, Trpc1, Ephb3, Fes, Sema6c, Pak1, Pik3cd, Ppp3ca, Epha7, Sema3a, Epha1, Srgap1, Srgap2, Trpc3, Cdk5, Rgma, Sema3b, Plcg1, Pak4, Sema3f, Sema6a, Fyn, Plxnb1, Ppp3cc, Camk2b, Nfatc2, Ngef, Sema4g, Camk2d, Ablim2* |
| KEGG | Calcium signaling pathway | 70 | 8.69E-05 | *Calml4, Casq1, Pdgfra, Itpka, Nfatc4, Tpcn2, Pde1b, Pdgfrb, Itpr3, Ngf, Atp2a3, Plcb2, Cd38, Ntrk1, Slc25a4, Cacna1a, Pde1c, Egf, Vegfd, P2rx7, Nos3, Adora2a, Adcy3, Mylk, Ppif, Phkg2, Plcd3, Plcd1, Phka1, Orai2, Plcb4, Tpcn1, Mcoln2, Mst1r, Ppp3ca, Phka2, Atp2b4, Cacna1d, Ntrk3, Tfeb, Cysltr2, Plcb3, Erbb2, Adcy4, Vegfa, Vegfb, Phkg1, Adrb2, Adcy7, Tbxa2r, Plcg1, Vegfc, Sphk1, Ppp3cc, Camk2b, Camk1, Smim6, Flt1, Nfatc2, Mcoln1, Pdgfb, Adrb1, Camk2d, Ptafr, Tnnc1, Fgf2, Prkcg, Prkcb, Fgfr1, Adrb3* |
| KEGG | Hematopoietic cell lineage | 33 | 9.88E-05 | *Cd44, Csf2ra, Il7, Il7r, Cd3g, Epor, Il3ra, Tnf, Cd5, Itgb3, Thpo, Flt3l, Csf1, Gp1ba, Cd38, Itgam, Itga2b, Kitl, Kit, H2-DMa, Il1r2, Il1r1, Cd3e, H2-Eb1, Itga4, Cd33, Il6ra, Csf3r, H2-Ab1, H2-DMb1, Cd22, Cd37, Cd8a* |
| KEGG | Inflammatory mediator regulation of TRP channels | 41 | 1.04E-04 | *Calml4, Ptger2, Kng2, Pik3cd, Plcb3, Adcy4, Asic1, Adcy5, Cyp2c37, Itpr3, Ngf, F2rl1, Map2k6, Cyp2c38, Plcb2, Trpv4, Adcy7, Plcg1, Pik3r2, Ntrk1, Cyp2j9, Camk2b, Il1r1, Cyp2c39, Mapk13, Cyp4a14, Trpv1, Cyp2j8, Mapk12, Trpv3, Adcy6, Adcy3, Asic3, Camk2d, Prkcd, Ptger4, Mapk11, Trpv2, Prkcg, Prkcb, Plcb4* |
| KEGG | Human T-cell leukemia virus 1 infection | 69 | 1.06E-04 | *Cdc20, Gps2, Tln1, Anapc5, Cd3g, Nfatc4, Spi1, H2-T10, Adcy5, Mad2l1, Crebbp, Zfp36, Map3k14, Crtc2, Atr, Il15, Pik3r2, Slc25a4, H2-DMa, Il1r2, Chek1, Il1r1, Ccne2, Nfkb2, Crtc1, Adcy6, Il15ra, Adcy3, Cd40, Ltbr, H2-Ab1, Icam1, Ranbp3, Pik3cd, Ppp3ca, Anapc4, Slc2a1, Fos, Adcy4, Tcf3, Myc, Tnf, Tbpl1, Ccna2, Anapc15, Bub1b, Kat5, Adcy7, Ccnd3, Ikbkb, Egr2, Ppp3cc, Jun, H2-Q4, Nfatc2, Relb, Cd3e, H2-Eb1, Rasl2-9, Bax, Lck, Il2rb, H2-Q6, Jak3, Trrap, H2-DMb1, Ccnb2, Tbp, Kat2a* |
| KEGG | NF-kappa B signaling pathway | 35 | 1.85E-04 | *Tnfsf13b, Eda, Birc2, Tnfsf14, Ticam2, Card10, Gadd45g, Tnf, Syk, Ube2i, Map3k14, Card14, Pias4, Plcg1, Ikbkb, Lat, Bcl2, Btk, Il1r1, Relb, Vcam1, Nfkb2, Traf1, Tab1, Pidd1, Cd40, Plau, Tnfaip3, Lck, Bcl2a1d, Ltbr, Tradd, Prkcb, Gadd45b, Icam1* |
| KEGG | Rap1 signaling pathway | 60 | 2.01E-04 | *Calml4, Tln1, Pdgfra, Pdgfrb, Adcy5, Rap1gap, Ngf, Map2k6, Rapgef5, Plcb2, Rras, Pik3r2, Itgam, Lat, Itga2b, Lpar1, Actg1, Kit, Tiam1, Mapk13, Evl, Egf, Vegfd, Arap3, Adora2a, Enah, Adcy6, Prkd2, Adcy3, Plcb4, Pik3cd, Apbb1ip, Id1, Plcb3, Adcy4, Vegfa, Vegfb, Rapgef3, Itgb3, Sipa1, Rapgef2, Csf1, Adcy7, Rasgrp2, Plcg1, Kitl, Vegfc, Skap1, Rgs14, Flt1, Pdgfb, Igf1r, Mapk12, Mapk11, Fgf2, Prkcg, Prkcb, Rapgef1, Fgfr1, Vav1* |
| KEGG | Hippo signaling pathway - multiple species | 14 | 2.73E-04 | *Tead4, Frmd6, Tead1, Rassf4, Tead3, Sav1, Rassf2, Csnk1e, Fat4, Dchs1, Rassf1, Wtip, Tead2, Pak1* |
| KEGG | Homologous recombination | 18 | 3.14E-04 | *Rad52, Bard1, Rad54b, Rad50, Rbbp8, Xrcc3, Palb2, Pold4, Brca2, Rad51b, Uimc1, Top3b, Brca1, Top3a, Pold3, Pold1, Abraxas1, Topbp1* |
| KEGG | ABC transporters | 21 | 3.20E-04 | *Abcg3, Abcg5, Abcb6, Abcg1, Abcd2, Abcc1, Abcd4, Abcb8, Abcb9, Abca4, Abcg8, Defb1, Abca14, Abcb1a, Abcc5, Abcc3, Abcc4, Abcc6, Abcc9, Abcc10, Abca7* |
| KEGG | Fluid shear stress and atherosclerosis | 44 | 3.67E-04 | *Calml4, Gstp3, Gsta1, Ncf2, Pik3cd, Klf2, Gstm1, Fos, Gstt3, Gstm3, Vegfa, Mgst3, Tnf, Nqo1, Map2k6, Itgb3, Pias4, Cyba, Trpv4, Pik3r2, Ikbkb, Gstt2, Itga2b, Bcl2, Actg1, Il1r2, Il1r1, Jun, Mapk13, Vcam1, Pdgfb, Arhgef2, Hmox1, Nos3, Mapk7, Mapk12, Map2k7, Map3k5, Acvr2b, Mapk11, Mef2a, Gm3776, Icam1, Ncf1* |
| KEGG | Th1 and Th2 cell differentiation | 30 | 3.86E-04 | *Ppp3ca, Stat4, Jag1, Fos, Dll4, Jag2, Cd3g, Cd247, Plcg1, Ikbkb, Lat, Dll1, H2-DMa, Gata3, Ppp3cc, Jun, Tyk2, Nfatc2, Mapk13, Cd3e, H2-Eb1, Mapk12, Lck, Il2rb, Maml2, Jak3, H2-Ab1, Mapk11, H2-DMb1, Maf* |
| KEGG | Intestinal immune network for IgA production | 18 | 6.11E-04 | *Tnfsf13b, Itgb7, H2-DMa, Cd28, Cd80, Tnfsf13, H2-Eb1, Itga4, Icos, Il15ra, Map3k14, Cd40, Icosl, Ltbr, Tnfrsf13b, H2-Ab1, Il15, H2-DMb1* |
| KEGG | Platelet activation | 38 | 7.17E-04 | *Pik3cd, Myl12b, Tln1, Ppp1r12a, Apbb1ip, Plcb3, Adcy4, Pik3r6, Adcy5, Arhgef1, Itpr3, Syk, Fermt3, Itgb3, Plcb2, Pik3r5, Adcy7, Rasgrp2, Gp1ba, Tbxa2r, Pik3r2, Ptgs1, Fcer1g, Itga2b, Gucy1a1, Fyn, Actg1, Btk, Fcgr3, Mapk13, Vwf, Nos3, Mapk12, Adcy6, Adcy3, Mylk, Mapk11, Plcb4* |

Table S2. Enrichment analysis of 4,507 down-regulated genes between WT and KL– in the 1st replicate

| **Category** | **Term** | **Count** | **PValue** | **Genes** |
| --- | --- | --- | --- | --- |
| GOBP | protein transport | 260 | 1.14E-42 | Cope, Mtx2, Lin7a, Snx16, Dnajc14, Selenbp2, Pex5, Snx18, Snap29, Vps53, Cog8, Ap2b1, Rab6a, Nsf, Atg4a, Vps36, Pgap1, Stxbp3, Pex14, Rab29, Copa, Snx9, Ndc1, Nbas, Ipo11, Actn4, Tmed2, Cadps2, Cetn2, Cmtm6, Snx14, Cog5, Vps35, Ap3s1, Sft2d2, Rabif, Yipf5, Hikeshi, Sec22b, Arl8a, Rab17, Ipo7, Ap1b1, Rab22a, Rbsn, Cetn3, Rab5b, Stx1b, Rab9, Mtm1, Stx18, Rd3, Arcn1, Tmem9, Cog6, Rab10, Afm, Sil1, Appbp2, Exoc3, Cog7, Rab5a, Atg4a-ps, Golph3, Sar1b, Vti1a, Rab1a, Rab18, Cse1l, Tnpo1, Ap2a2, Gosr2, Chmp1b, Chmp4b, Arl6ip5, Vps4b, Aagab, Seh1l, Gabarapl2, Vps35l, Timm17b, Sec13, Gpr89, Tmed3, Erc1, Rab21, Rufy1, Ppt1, Aftph, Chmp5, Nxt2, Rabep1, Praf2, Chmp3, Copg1, Arf3, Timm23, Chmp6, Bcap31, Atg3, Xbp1, Pex7, Tram1, Vps54, Gdi2, Hook3, Snx3, Heatr5b, Sec63, 2610002M06Rik, Kpna4, Mcfd2, Nmd3, Snx13, Rrbp1, Zmat3, Stx12, Tomm7, Ap3b1, Hspb11, Scamp1, Chmp7, Sec23b, Copb2, Ran, Snx19, Atg4b, Arfgap2, Arf1, Copz1, Vps33a, Derl1, Rabgap1l, Arf6, Plekhf2, Gopc, Sec23a, Dennd4c, Unc119b, Vta1, Snx4, Vhl, Cog4, Ccdc91, Dennd1b, Tmed9, Atg9a, Surf4, Hook1, Kpna6, Washc5, Rab36, Snx24, Washc3, Cav1, Ap3m1, Serp1, Stam, Nxt1, Exoc6, Jagn1, Vps26b, Sec61b, Tnpo3, Snx1, Sec61a1, Rab5c, Pex13, Xpo7, Ap3s2, Dtx3l, Timm21, Lman1, Timm22, Psen1, Washc4, Kpna1, Fras1, Chchd4, Ap1g1, Exoc4, Eps15, Sec61g, Atg4c, Ykt6, Dennd10, Uso1, Rab1b, Cnih4, Lman2, Yif1a, Vti1b, Rab8a, Ap5m1, Timm29, Tmed10, Scfd1, Uevld, Ap4s1, Arfgef1, Copg2, Rab2a, Nup62, Kpna3, Vps37a, Copb1, Gga2, Vps45, Snx30, Ipo8, Bbs9, Tom1l1, Vps41, Ap2m1, Arf4, Nup37, Dag1, Yif1b, Arl8b, Lin7c, Snap23, Vps29, Stam2, Snx27, Rab14, Tbc1d5, Timm17a, Pex2, Bet1, Aktip, Vps13b, Tmed4, Kdelr2, Rab7, Snx7, Sar1a, Kpnb1, Snx25, Chp1, Kdelr1, Gosr1, Ranbp6, Scfd2, Snx2, Atg10, Necap1, Timm8b, Plekhm1, Psen2, Dnajc15, Vps26a, Myo5b, Sec62, Tnks, Rab38, Eif5a, Tsg101, Sec22a, Gabarap, Atg7 |
| GOBP | lipid metabolic process | 259 | 4.37E-32 | Acsm1, Cyp7b1, Slc27a5, Acpp, Gstm4, Asah1, Fabp5, Hexa, Hmgcr, Cyp2e1, Fads6, Abhd3, Mtmr2, Adhfe1, Srd5a2, Hsd11b1, Lipc, Acaa1b, Cyp1a2, Retsat, Gstp1, Aadac, Acss3, Echs1, Fitm2, Hsd17b7, Eci1, Chpt1, Oxsm, Lypla1, Faah, Acsm3, Ephx1, Nsdhl, Aldh1a1, Fads1, Cyp8b1, Plcxd2, Sult2a8, Acot11, Crkl, Mvk, Hsd17b12, Abhd5, Bdh2, Hmgcs2, Samd8, Scp2, Arsa, Acnat1, Cpt2, Scly, Insig1, Prkaa1, Hacd2, Adh4, Mtm1, Lipg, Dhrs9, Aacs, Lima1, Pcyt1a, Hadh, Cyp2u1, Mttp, Nr1h3, Hacd3, Elovl1, Pik3c3, Agmo, Elovl3, Degs1, Tmem86b, Cyp4v3, Acsl4, St3gal1, Pgap6, Hadhb, Fasn, Atp5b, Acsl5, Nceh1, Mvd, Abhd6, Srd5a1, Agpat2, Hsd17b4, B3galt1, Dgat2, Zadh2, Acadsb, Gla, Pafah1b2, Plcb1, Pi4k2b, Pemt, Gpld1, Gpam, Sptlc1, Hmgcs1, Hacd1, Tecr, Oma1, Crk, Prxl2b, C3, Akr1c6, Xbp1, Chka, Enpp2, Gsta3, Hdlbp, Slc16a1, Baat, Elovl5, Adipor2, Etnk1, Erlin2, Sacm1l, Abhd4, Adtrp, Cyp2c40, Psap, Nus1, Plpp3, Gpx1, Rdh10, Adipor1, Rdh11, Prkaa2, Akr1c21, Cyb5r3, Acacb, Pm20d1, Pnpla8, Acly, Lclat1, Selenoi, Acat1, Ugt2b5, Abhd12, Hsd17b11, Pik3c2a, Cyp2c23, Decr1, Erlin1, Fntb, Pip4p2, Insig2, Fdft1, Lipa, Pisd, Mcat, Fads2, Ptpn11, Ugt1a9, Rdh14, Pafah1b1, Ldlrap1, Acaa2, Rnf213, Bscl2, Cyp2c29, Akr1a1, Cers2, Sqle, Pcx, Acads, Hadha, Ugt1a1, Pcsk9, Serinc5, Smpd1, Serinc1, Mid1ip1, Dhcr7, Acox3, Erg28, Srebf2, Pi4k2a, Acad11, Prdx6, Slc27a2, Ptges, Pdss2, Pcca, Apof, Hpgd, Acot8, Apob, Slc27a4, Ebp, Angptl3, Fuca1, Hmgcl, Lactb, Npc2, Elovl2, St6galnac6, Fitm1, Comt, Acad8, Msmo1, Acsf2, Hsd17b2, Bco1, Sc5d, Daglb, Lss, Acsf3, Acsl3, Dhdds, Them4, Ephx2, Mgll, Apoa1, Cyp51, Acox2, Acadvl, Dolk, Ptdss1, Mboat7, Acot1, Ptgr2, Acadm, Adh5, Ldlr, Ttc39b, Elovl6, Acss2, Acox1, Atp5a1, Bdh1, Mtmr6, Dhrs3, Plbd1, Nfe2l1, Fdps, Cdipt, Ces2c, Bco2, Lrat, Dhcr24, Rab7, Pmvk, Pecr, Sgpp1, Crot, Lpgat1, Gm2a, Scd1, Acsl1, Pten, Sgms2, Cyp2r1, Tm7sf2, Ech1, Thrsp, Decr2 |
| GOBP | ER to Golgi vesicle-mediated transport | 67 | 3.14E-21 | Cope, Ero1b, Spast, Lmf1, Sar1b, Vti1a, Trappc13, Rab1a, Sec23a, Sec24d, Yipf6, Trappc12, Trappc11, Yif1b, Pgap1, Sec13, Tmed9, Vapa, Trappc4, Tmed3, Tex261, Vcp, Trappc1, Copa, 0610009B22Rik, Trappc9, Tfg, Tmed2, Bet1, Copg1, Hyou1, Ergic3, Tmed4, Vapb, Kdelr2, P4hb, Bcap31, Yipf5, Sar1a, Trappc3, Sec22b, Pdcd6, Lman1, Kdelr1, Gosr1, Trappc2, Yipf4, Cul3, Arcn1, Trappc8, Tmed7, Ergic1, Ykt6, Uso1, Mia2, Rab1b, Lman2, Cnih4, Yif1a, Steep1, Tmed10, Scfd1, Sec23b, Copg2, Sec22a, Copb1, Copb2 |
| GOBP | intracellular protein transport | 122 | 4.25E-20 | *Arf1, Srpr, Copz1, Vps33a, Rabgap1l, 5730455P16Rik, Arf6, Sec23a, Sec24d, Ap2b1, Rab6a, Stx8, Nsf, Exph5, Tbc1d9b, Tmed9, Stxbp3, Rab29, Copa, Snx9, Tbc1d8b, Cltc, Ipo11, Ap3m1, Exoc6, Tmed2, Exoc6b, Vps26b, Ric1, Sec61b, Vps35, Ap3s1, Snx1, Rab5c, Xpo7, Ap3s2, Rab17, Ap1b1, Ipo7, Rab22a, Rab5b, Stx1b, Rph3al, Ap1g1, Tbck, Stx18, Sec61g, Tmed7, Uso1, Rab1b, Rab10, Vti1b, Tmed10, Scfd1, Ap4s1, Appbp2, Copg2, Rab2a, Copb1, Cog7, Gga2, Rab5a, Vps45, Tlk1, Ipo8, Sar1b, Tom1l1, Vti1a, Vps41, Ap2m1, Rab1a, Rab18, Arf4, Cse1l, Tnpo1, Ap2a2, Tbc1d13, Evi5, Tmed3, Vps29, Tbc1d12, Snx27, Rab14, Rab21, M6pr, Tbc1d5, Arl4a, Tbc1d30, Tbc1d14, Arl5b, Copg1, Rab32, Arf3, Sytl4, Tmed4, Timm23, Rab7, Chm, Bcap31, Mpdz, Sar1a, Kpnb1, Pdcd6, Scfd2, Slu7, Snx2, Tbc1d22a, Arl1, Snx13, Psap, Clta, Stx12, Ngfr, Vps26a, Ap3b1, Cltb, Vps26c, Arl5a, Sec23b, Rab38, Copb2, Selenos* |
| GOBP | fatty acid metabolic process | 92 | 9.66E-20 | *Acsm1, Acsl3, Acsl4, Acat1, Hadhb, Decr1, Fasn, Slc27a5, Cyp2c23, Them4, Mgll, Asah1, Acsl5, Lipa, Fabp2, Amacr, Mcat, Acox2, Fads2, Cyp2e1, Fads6, Acadvl, Hsd17b4, Acaa2, Acot1, Acbd4, Lipc, Acadsb, Acaa1b, Cyp1a2, Cyp4a12b, Acadm, Echs1, Abcb11, Eci1, Acads, Hadha, Gpam, Oxsm, Elovl6, Lypla1, Acox1, Hacd1, Faah, Tecr, Acsm3, Cryl1, Fads1, Gnpat, Acot11, Abhd5, Prxl2b, Dbi, Acbd5, C3, Acox3, Pecr, Crot, Acad11, Slc27a2, Ptges, Acnat1, Scd1, Acsl1, Cpt2, Prkaa1, Hpgd, Hacd2, Baat, Acot8, Elovl5, Adipor2, Slc27a4, Angptl3, Aacs, Hadh, Elovl2, Alkbh7, Adipor1, Prkaa2, Acacb, Pm20d1, Ndufs6, Hacd3, Elovl1, Acsf2, Ech1, Elovl3, Degs1, Pnpla8, Decr2, Acsf3* |
| GOBP | vesicle-mediated transport | 113 | 2.59E-19 | *Arfgap2, Arf1, Cope, Copz1, Vps33a, Arf6, Sec23a, Ap2b1, Rab6a, Trappc12, Stx8, Nsf, Tvp23b, Stxbp3, Trappc1, Rab29, Copa, Rab36, Clint1, Syt11, Tbc1d8b, Cltc, Kif1b, Ap3m1, Tmed2, Jagn1, Kif3b, Ap3s1, Sft2d2, Yipf5, Ap3s2, Sec22b, Trappc3, Ston2, Mon1b, Ap1b1, Lman1, Stx1b, Rin2, Rab30, Ap1g1, Cul3, Stx18, Yipf1, Arcn1, Ykt6, Uso1, Cnih4, Rab10, Yif1a, Vti1b, Ap5m1, Rtn3, Tmed10, Scfd1, Ap4s1, Copg2, Rab2a, Hspa8, Copb1, Gga2, Vps45, Sar1b, Vti1a, Vps41, Ap2m1, Rab1a, Arf4, Ap2a2, Gosr2, Trappc11, Hspa14, Bloc1s5, Ston1, Sec13, Trappc4, Snap23, Cnih1, Arl4a, Arl5b, Rabep1, Bet1, Copg1, Syt1, Rab32, Vps13b, Arf3, Ergic3, Kdelr2, Chm, Bcap31, Yipf2, Sar1a, Gdi2, Kdelr1, Gosr1, Scfd2, Hspa1b, Necap1, Arl1, Mcfd2, Ergic1, Vamp4, Clta, Stx12, Myo5b, Ap3b1, Cltb, Arl5a, Sec23b, Rab38, Sec22a, Copb2* |
| GOBP | ubiquitin-dependent ERAD pathway | 48 | 4.47E-16 | *Amfr, Fbxo6, Faf1, Ccdc47, Erlin1, Faf2, Dnajb2, Derl1, Hspa5, Sgtb, Hsp90b1, Canx, Psmc6, Derl3, Os9, Sel1l, Rnf185, Man1a, Ube2g2, Syvn1, Dnajb12, Edem1, Trim13, Man1b1, Rnf121, Rnf103, Jkamp, Dnajb9, Dnajc18, Erlin2, Ube2j1, Ubxn4, Vcp, Ube4a, Erlec1, Ubqln1, Derl2, Marchf6, Calr, Ubxn8, Dnajb14, Eif2ak3, Stt3b, Nploc4, Selenos, Sec61b, Calr3, Rnf5* |
| GOBP | autophagy | 84 | 1.06E-14 | *Atg4b, Atg4a-ps, Tmem208, Trim12c, Tmbim6, Smcr8, Vps33a, Vps41, Vti1a, Snap29, Rab1a, Gopc, Chmp1b, Chmp4b, Atg4a, Vps4b, Uvrag, Retreg3, Arsb, 1600014C10Rik, Gabarapl2, Lgals8, Atg9a, Trappc4, Tex264, Epm2a, Atg13, Trp53inp1, Vcp, Sqstm1, Rmc1, Depp1, Tm9sf1, Irgm1, Syt11, Retreg2, Ubqln1, Tbc1d5, Tmem59, Stbd1, Chmp5, Cltc, Rb1cc1, Trp53inp2, Atg101, Eva1a, Rnf41, Acbd5, Rab7, Chmp6, Cisd2, Gabarapl1, Ulk2, Atg3, Xbp1, Wipi2, Arsa, Atg5, Rnf185, Atg12, Anxa7, Psen1, Prkaa1, Atg10, Plekhm1, Lamp2, Fnbp1l, Fundc1, Atg4c, Ei24, Rab1b, Creg1, Dram2, Rab8a, Pink1, Chmp7, Prkaa2, Dap, Becn1, Clec16a, Pik3c3, Epg5, Gabarap, Atg7* |
| GOBP | translation | 109 | 2.87E-14 | *Hars, Rpl5, Mrpl3, Mrps5, Tars, Mrpl37, Gfm1, Eif4a2, Eif2a, Mrpl39, Tufm, Eif2s3x, Mrpl21, Paip2, Iars, Rpl4, Eif2b4, Rpl39, Eif2b2, Srbd1, Fars2, Wars, Eif4h, Mrps23, Gfm2, Eef1g, Mrps18a, Eif4g1, Mrpl32, Eif3g, Etf1, Mrps10, Eif2d, Gars, Gatc, Mtrfr, Dars2, Eif3k, Nars2, Ears2, Mrpl1, Aars, Eif4e2, Eif4e, Eif3i, Gspt1, Eif3a, Eif4g2, Larp4, Mrps6, Eif3e, Eef1b2, Tarsl2, Eif2s2, Mrpl9, Eif5b, Mrpl30, Aimp1, Farsb, Mrrf, Nars, Eif2s1, Rpl9, Mrps18c, Rps3, Eefsec, Eif3l, Mrpl20, Dars, Mrps12, Yars2, Mrpl49, Yars, Eif3h, Eif3c, Kars, Eif2ak3, Rps29, Mapkap1, Mrpl15, Eif3m, Mrps11, Eif1ax, Mrpl19, Gatb, Eif4b, Gm15501, Dhx29, Mcts1, Mrpl10, Mtrf1, Mrpl16, Egfr, Efl1, Rps27l, Mrpl34, Mrps14, Mrpl13, Mars1, Mrps18b, Eif2b5, Eprs, Mrpl22, Eif5a, Mrpl36, Mrpl35, Rars, Eef1a1, Eef2* |
| GOBP | ubiquitin-dependent protein catabolic process | 103 | 8.40E-14 | *Amfr, Klhl8, Ubl7, Arih1, Rnf149, Herc4, Ubr3, Herc6, Ndfip1, Os9, Psma3, Ube2r2, Eloc, Nae1, Cacul1, Rnf213, Usp8, Vcp, Uchl3, Ube4a, Psma6, Psma2, Usp38, Trp53inp2, Arel1, Ube2z, Spop, Psma4, Ube2n, Ube2l6, Fbxo4, Rnf114, Itch, Usp47, Dtx3l, Usp14, Uba1, Mkrn2, Usp25, Rnf217, Rnf185, Syvn1, Usp46, Psmd11, Psma8, Rnf13, Uba6, Usp12, Cul3, Rnf19b, Rnf11, Cul2, Ube2g1, Smurf2, Trim32, Rnf125, Hectd1, Psma1, Rnf146, Rnf34, Ddb1, Rnf6, Rc3h1, Cdc34, Rnf144b, Usp18, Ube2h, Mib1, Atxn3, Rnf130, Ubqln1, Psmc2, Uchl5, Spsb4, Ppp1r11, Nploc4, Rnf5, Skp1, Usp9x, Wwp1, Xbp1, Ube2g2, Rnf168, Rnf14, Fem1c, Trpc4ap, Fbxw8, Fbxo8, Uchl4, Ube2l3, Cul1, Ndfip2, Psma5, Psmd13, Asb2, Kctd21, Usp24, Ankib1, Usp32, Ube3c, Cops3, Rnf7, Usp4* |
| GOBP | steroid metabolic process | 63 | 2.09E-13 | *Ugt2b37, Ugt2b5, Cyp7b1, Ugt2b1, Erlin1, Insig2, Apoa1, Mvd, Fdft1, Cyp51, Hmgcr, Srd5a1, Cyp2e1, Srd5a2, Ldlrap1, Hsd11b1, Akr1c20, Cyp1a2, Ldlr, Rdh7, Hmgcs1, Bdh1, Pcsk9, Nsdhl, Nfe2l1, Cyp3a16, Sult2a8, Mvk, Fdps, Dhcr7, Dhcr24, Cyp3a11, Ugt2b38, Hmgcs2, Sdr9c7, Pmvk, Erg28, Srebf2, Akr1c6, Akr1c12, Akr1c13, Apof, Akr1c19, Insig1, Hdlbp, Prkaa1, Akr1c14, Apob, G6pc, Ebp, Dhrs9, Erlin2, Lima1, Dhrs4, Mttp, Npc2, Tm7sf2, Prkaa2, Akr1c21, Cyb5r3, Msmo1, Hsd17b2, Sc5d* |
| GOBP | protein folding | 55 | 9.59E-13 | *Trap1, Cdc37l1, Erp44, Dnaja4, Hsp90aa1, Grpel1, Ahsa1, Tbcd, Hspa4l, Pdia5, Pdcl, Tbca, Hspe1, Cct2, Mesd, Dnajb11, Qsox1, Dnaja1, Fkbp9, Pdia3, Hspd1, Ppil3, Cct5, Dnaja3, Pdia4, Tbcel, Cdc37, Calr, Cct4, Dnaja2, Txndc9, Clpx, Dnajb1, Dnajb6, Ppib, P4hb, Mkks, Hspa4, Dnajc1, Cct8, Ahsa2, Hsph1, Hsp90b1, Canx, Pdilt, Hspa1b, Cct7, Dnajb4, Hspa9, Cct6a, Hsp90ab1, Txndc5, Cct3, Hspa8, Calr3* |
| GOBP | mitochondrial translation | 48 | 1.22E-12 | *Hars, Mrpl19, Gatb, Mrpl3, Mrpl9, Mrps5, Mrps10, Mrpl30, Chchd1, Mrpl37, Mrpl10, Mrpl16, Mrps27, Mrpl44, Gatc, Mrpl39, Mrpl48, Mrpl21, Mrpl40, Noa1, Mrpl34, Mrps18c, Mrps25, Mrps31, Mrpl1, Mrps14, Mrpl20, Mrpl13, Mrps12, Mrpl42, Mrpl49, Mrps18b, Fastkd2, Mrpl50, Mrps23, Mrps24, Mrps6, Mrpl38, Gfm2, Mrps18a, Mrpl32, Mrpl22, Mrpl36, Mrpl35, Mrpl46, Mrpl15, Mrps22, Mrps11* |
| GOBP | cholesterol metabolic process | 55 | 2.23E-12 | *Cyp7b1, Erlin1, Insig2, Apoa1, Mvd, Fdft1, Cyp51, Lipa, Hmgcr, Fech, Ldlrap1, Lipc, Aplp2, Cyp1a2, Sqle, Ldlr, Hmgcs1, Pcsk9, Nsdhl, Saa1, Smpd1, Nfe2l1, Sult2a8, Mvk, Ces1c, Fdps, Dhcr7, Dhcr24, Hmgcs2, Pmvk, Srebf2, Abca2, Apof, Insig1, Hdlbp, Prkaa1, Apob, Ebp, Cebpa, Angptl3, Erlin2, Ces1e, Cat, Pon1, Lima1, Mttp, Npc2, Tm7sf2, Pctp, Apon, Lrp5, Prkaa2, Cyb5r3, Msmo1, Ces1f* |
| GOBP | proteasome-mediated ubiquitin-dependent protein catabolic process | 80 | 9.97E-12 | *Rnf34, Fem1b, Gid8, Rnf4, Ddb1, Faf2, Dnajb2, Psmd2, Nsfl1c, Klhl42, Derl1, Edem3, Clock, Rad23b, Psmc6, Cop1, Cdc34, Ube2h, Psmc5, Trim13, Ppp2r5c, Gid4, Znrf2, Epm2a, Vcp, Plaa, Ppp2cb, Atxn3, Rnf187, Nhlrc3, Psmc2, Fbxl17, Spsb4, Psmd14, Kbtbd7, Rad23a, Klhdc2, Arel1, Psmd7, Marchf6, Tnfaip1, Spop, Pja2, Skp1, Crbn, Ifi27, Fbxo9, Itch, Ctnnb1, Trim2, Wwp1, Psmd1, Psmc1, Usp14, Ube2w, Psmd6, Psmc4, Fem1c, Mtm1, Cul3, Cul1, Ube2a, Psmd8, Fbxw11, Psma5, Tbl1x, Ube2k, Asb2, Ube2g1, Rmnd5a, Psmc3, Gsk3b, Ubxn2a, Appbp2, Smurf2, Clec16a, Ubxn7, Arntl, Fbxl22, Hectd1* |
| GOBP | retrograde vesicle-mediated transport, Golgi to ER | 29 | 1.58E-11 | *Ergic3, Kdelr2, Cope, Copz1, Sec22b, Golph3, Arf4, Kdelr1, Rab6a, Uvrag, Cog4, Pitpnb, Stx18, Rer1, Arcn1, Ergic1, Gbf1, Golph3l, Copa, Lman2, Plpp3, Nbas, Tmed10, Scfd1, Dnajc28, Copb2, Tmem115, Cog7, Arf3* |
| GOBP | fatty acid beta-oxidation | 34 | 3.83E-11 | *Acat1, Hadhb, Decr1, Bdh2, Acox3, Acad9, Abcd3, Crot, Acad11, Acox2, Pex5, Scp2, Pex7, Slc27a2, Acadvl, Acat2, Cpt2, Slc25a17, Echdc1, Hsd17b4, Acaa2, Acadsb, Acaa1b, Acadm, Echs1, Hadh, Eci1, Hadha, Pex2, Acat3, Acox1, Mtln, Acad8, Ech1* |
| GOBP | cholesterol biosynthetic process | 28 | 5.77E-11 | *Mvk, Ces1c, Idi1, Fdps, Dhcr7, Dhcr24, Hmgcs2, Insig2, Pmvk, Apoa1, Mvd, Fdft1, Cyp51, Lipa, Hmgcr, Insig1, Prkaa1, Ebp, Ces1e, Hsd17b7, Tm7sf2, Hmgcs1, Prkaa2, Cyb5r3, Ces1f, Msmo1, Nsdhl, Lss* |
| KEGG | Metabolic pathways | 583 | 6.50E-56 | *Sdhd, Ahcyl1, H6pd, Alg6, Cyp2c68, Gstm4, Hsd3b3, Pigf, Aldh9a1, Gmppb, Hexa, Impa1, Glo1, Cyp2e1, Suclg1, Mtmr2, Srd5a2, Pgap1, Bhmt, Pigt, Uap1, Csad, Echs1, Hsd17b7, Rpn2, Chpt1, Dlst, Idh3b, Uck2, Acsm3, Vnn1, Fig4, Cyp3a16, Cyp8b1, Mvk, Hmgcs2, Ndufa9, Atp6v0c, Bckdha, Pgd, Gaa, Cox7a2l, Eno1, Scp2, Scly, Camkmt, Uqcrh, Hacd2, Adh4, Lta4h, Car14, 9130409I23Rik, Lipg, Mocos, Pde5a, Acp2, Pcyt1a, Hadh, Atp6v1c1, Dlat, Dad1, Cyp2u1, Cmpk1, Ndufs2, Cyp2c50, Bpgm, Atp5g2, Aldh8a1, Elovl3, Degs1, Mocs2, Tmem86b, Cyp2j5, Odc1, Alas2, Gfpt1, Hadhb, Fasn, Atp5b, Rpe, Mvd, Mthfd2, Ndufa4, Aldob, Atp6v0d1, Ldhd, Mat2a, Dgat2, Acadsb, Pafah1b2, Glul, Fktn, Galk2, Gstk1, Acyp1, St3gal3, Ppt1, Btd, Kmo, Tecr, Mgat5, Alg1, Idi1, Rgn, Ndufb5, Car1, Tpk1, Coq5, Car3, Aco1, Baat, Urah, Sacm1l, Pik3ca, Cyp2c40, Nme7, Cbs, Plpp3, Gpx1, Hagh, Cox5a, Ddost, Ndufv3, Gch1, Pank3, Alg14, Ndufa1, Acly, Kyat1, Atp6v0e, Selenoi, Nat8f1, Acat1, Mtap, Aspdh, Pgm3, Ddc, Hprt, Fdft1, Cps1, Pisd, Gstz1, Smyd2, Mcat, Amacr, Fads2, Tpi1, Ugt1a9, Ganab, Acaa2, Gcnt4, Glb1, Ndufa3, Arsb, Aadat, Man1b1, Cyp2a12, Cers2, Pah, Sqle, Pcx, Sucla2, Ndufb9, Taldo1, Dbt, Hadha, Prps2, Cmbl, Suox, Gnmt, Ugt2a3, Gulo, Prune1, Cox6b1, B3gat3, Nme1, Alg5, Pdha1, Qdpr, Aldh1l1, Dhcr7, Pnp, Uxs1, Acox3, Cyp3a11, Otc, Haao, Pigx, Prdx6, Ptges, Atp5k, Cyp2j6, Crppa, Galnt4, Thtpa, Ndufb11, Ebp, Glt28d2, Glce, Ado, Gamt, Pycrl, Atp5pb, Gfus, Dhrs4, Pigh, Elovl2, Ugp2, St6galnac6, Ndufs1, Comt, Pigq, Sc5d, Lss, Ugt2b37, Pgm1, Mgll, Ephx2, Ggct, Cyp51, Fh1, Uck1, Fech, Nfs1, Car5a, Nudt12, Acp5, Dhdh, Gphn, Cyp2c54, Galm, Acot1, Ndufs8, Atp5c1, Acadm, Cyp4a12b, Extl2, Dglucy, Pcbd1, Gmpr2, Pygl, Acss2, Bdh1, Atp5a1, Dhrs3, Tkfc, Ugt1a6b, Cyp2c70, Fdps, Smyd1, Aasdhppt, C1galt1c1, Pmvk, Srr, Sgpp1, Uqcrc1, Kynu, Hnmt, Gusb, Man1a, Sqor, Atp6ap1, Pten, Sgms2, Adss, Pomt2, Alg9, Suclg2, Prps1l3, Tgds, Nadk2, Tm7sf2, Gmps, Slc33a1, Mthfd1, Mgam, Bpnt1, Eprs, Ugt1a7c, Mccc1, Prodh2, Chdh, Acsm1, Uox, Hccs, Ugt2b1, Slc27a5, Asah1, Hmgcr, Selenbp2, Glud1, St6gal1, Adk, Mgat1, Hsd11b1, Gldc, Lipc, Rxylt1, Acaa1b, Cyp1a2, Gsr, Gstp1, Acss3, Phospho2, Pigg, Nit2, Enpp4, Fkrp, Oxsm, Anpep, Car8, Gstm6, Nsdhl, Aldh1a1, Pgam1, Fads1, Stt3b, Prdm2, Ndst1, Ugt2b35, Hsd17b12, Pla2g12b, Rbks, Aldh5a1, Abat, Ext2, Ugt2b38, Aldoc, Mdh1, Cpox, Arsa, Mpst, Guk1, Acnat1, Acat2, Dmgdh, Aox3, B4galnt3, C1galt1, Ugt2b34, Mpi, Pdxk, Mtm1, Nat8f2, Nsd2, Nat8, Papss1, Dck, Ppat, Dhrs9, Aacs, Cat, Sirt5, Paics, Ndufs4, Mccc2, Pigk, Tymp, Atp6v1e1, Elovl1, Hacd3, Me1, Pik3c3, Lias, Acsl4, St3gal1, Ggcx, Dpagt1, Grhpr, Rpn1, Acsl5, Fbp1, Idh1, Srd5a1, Agpat2, Fahd1, Atp6v1d, Entpd5, Aldh1a7, mt-Co1, Hsd17b4, Csgalnact2, B3galt1, Sephs2, Gcsh, Hmbs, Gla, Nadk, Plcb1, Gclc, Pi4k2b, Galnt1, Pemt, mt-Nd5, Gpld1, Gpam, Sptlc1, Hmgcs1, Coasy, Hacd1, Mat2b, Blvra, mt-Nd4, Gm10053, Stt3a, Nme6, Gatb, Itpk1, Prxl2b, Atp6v1h, Ndufv1, Car5b, Cmas, Chka, Fpgs, Enpp2, Dpys, Sdha, Fbp2, Carnmt1, mt-Cytb, Gsta3, Ethe1, Ugt2b36, Elovl5, Pgk1, G6pc, Etnk1, Khk, Cox7a2, Atp5j, Rdh10, Pgam2, Rdh11, Rfk, Bpnt2, Acacb, Dpyd, Mogs, Lclat1, Ugt2b5, Pik3c2a, Cyp2c23, Vkorc1l1, Ldha, Pigu, Pafah1b1, L2hgdh, Tkt, Amt, Pnpo, Adi1, Enpp3, Atp6v1a, Akr1a1, Cyp2c29, Art2b, Nt5c3, Fggy, Ndufb3, Hibch, Ndufv2, Gsto1, Nans, Acads, Kyat3, Bbox1, Gmds, Pipox, Pde7b, Ugt1a1, Alg3, Cryl1, Dhfr, Smpd1, Alg10b, Cs, Ndufa10, Ahcyl2, Nudt2, Amy1, Urod, Pdhx, Apip, Txndc12, Pi4k2a, Hibadh, Gatc, Pcca, Alg11, Acot8, Uqcrc2, Ears2, Hmgcl, Ak2, Gale, mt-Nd2, Vkorc1, Dpm3, Sord, Acad8, Lap3, Msmo1, Hsd17b2, Bco1, Mtmr4, Acsf3, Pank1, Acsl3, Prps1, Gns, Sdhc, Pde9a, Eno1b, Ddo, Sdhb, Gart, Maob, Acox2, Pklr, Tst, Dld, Acadvl, Dolk, Esd, Ptdss1, Chac2, Colgalt2, Pccb, Dera, Spr, Adh1, Atic, Mgst1, Fpgt, Tdo2, Adh5, Cox15, Rdh7, Aspa, Mmut, Elovl6, Shmt1, Acox1, Mtmr6, Pdhb, Ido2, Hmox2, Itpa, Mmab, Aldh2, Cdipt, Gclm, Lrat, Ak6, Dhcr24, Gstm7, Mgat2, Galnt10, Hgd, Scd1, Acsl1, Echdc1, Pomt1, Aldh7a1, Gbe1, Man1a2, Hao1, Cyp2r1, Sardh, Ugdh, Ugt1a5, Pigyl, Mars1, Psph, Acat3, Umps, Cycs, B3galnt2, Ndufs6, Gstp2, Cmpk2, Urad, Qprt* |
| KEGG | Protein processing in endoplasmic reticulum | 111 | 3.54E-34 | *Amfr, Magt1, Ero1b, Hsp90aa1, Dnajb2, Capn1, Derl1, Ubxn1, Os9, Hspa4l, Sec23a, Sec24d, Ganab, Dnajb12, Man1b1, Vcp, Pdia3, Rpn2, Plaa, Pdia6, Ngly1, Ssr1, Ubxn8, Marchf6, Stt3b, Sec61b, Sec61a1, Atf6, Hspbp1, Hsph1, Hsp90b1, Lman1, Derl3, Sel1l, Rnf185, Syvn1, Sec61g, Ube2j1, Dad1, Lman2, Ube2g1, Bag2, Sil1, Erlec1, Svip, Hsp90ab1, Txndc5, Ostc, Ern1, Hspa8, Ubxn6, Fbxo6, Rpn1, Prkcsh, Nsfl1c, Ssr2, Sar1b, Edem3, Rad23b, Eif2s1, Atf6b, Edem1, Sec13, Dnajb11, Dnaja1, Ubxn4, Atxn3, Bag1, Ubqln1, Pdia4, Dnajc5, Rad23a, Calr, Eif2ak3, Dnaja2, Nploc4, Stt3a, Rnf5, Hyou1, Dnajb1, Skp1, P4hb, Bcap31, Dnajc1, Sar1a, Xbp1, Ssr4, Eif2ak1, Hspa5, Tram1, Canx, Uggt1, Man1a, Ube2g2, Capn2, Hspa1b, Sec63, Man1a2, Cul1, Rrbp1, Ddost, Dnajc3, Ssr3, Derl2, Sec62, Ubxn2a, Sec23b, Tmem258, Edem2, Selenos, Mogs* |
| KEGG | Biosynthesis of cofactors | 83 | 9.75E-20 | *Alas2, Pank1, Lias, Ggcx, Aspdh, Ugt2b37, Ugt2b5, Ugt2b1, Vkorc1l1, Gmppb, Mthfd2, Ak3, Fech, Ugt1a9, Dld, Nfs1, Gphn, Mat2a, Pnpo, Spr, Hmbs, Akr1a1, Nadk, Phospho2, Gclc, Tdo2, Cox15, Oxsm, Shmt1, Coasy, Ugt1a1, Kmo, Dhrs3, Mat2b, Ugt2a3, Ugt1a6b, Ido2, Gulo, Dhfr, Nme1, Ggh, Nme6, Ugt2b35, Mmab, Rgn, Aldh2, Gclm, Urod, Ak6, Ugt2b38, Coq5, Tpk1, Haao, Kynu, Fpgs, Cpox, Gusb, Ugt2b36, Mpi, Ugt2b34, Pdxk, Adss, Ears2, Ugdh, Ak2, Ugt1a5, Nme7, Vkorc1, Nadk2, Ugp2, Mthfd1, Cmpk1, Rdh11, Gch1, Rfk, Eprs, Ugt1a7c, Umps, Cmpk2, Pank3, Mocs2, Bco1, Qprt* |
| KEGG | Peroxisome | 52 | 6.02E-15 | *Acsl3, Acsl4, Pex11b, Ephx2, Acsl5, Idh1, Amacr, Ddo, Acox2, Pex5, Nudt12, Slc25a17, Hsd17b4, Pex11a, Acaa1b, Prdx1, Pex14, Paox, Gstk1, Pex2, Acox1, Pipox, Gnpat, Mvk, Pex3, Acox3, Pmvk, Phyh, Pex13, Abcd3, Pecr, Crot, Pex19, Scp2, Pex7, Slc27a2, Acnat1, Acsl1, Pxmp4, Sod2, Baat, Acot8, Hao1, Pex16, Cat, Hmgcl, Dhrs4, Prdx5, Pex11g, Ech1, Nudt7, Decr2* |
| KEGG | Carbon metabolism | 64 | 2.06E-14 | *Prps1, Sdhd, Acat1, H6pd, Sdhc, Fbp1, Idh1, Rpe, Cps1, Eno1b, Fh1, Aldob, Glud1, Sdhb, Pklr, Tpi1, Dld, Esd, Suclg1, Tkt, Amt, Gldc, Pccb, Gcsh, Echs1, Adh5, Hibch, Pcx, Sucla2, Dlst, Idh3b, Acads, Taldo1, Prps2, Mmut, Shmt1, Acox1, Acss2, Pdhb, Tkfc, Pgam1, Pdha1, Cs, Rgn, Acox3, Pgd, Aldoc, Aco1, Mdh1, Eno1, Sdha, Fbp2, Acat2, Pcca, Pgk1, Hao1, Suclg2, Cat, Prps1l3, Dlat, Psph, Pgam2, Acat3, Me1* |
| KEGG | Parkinson disease | 109 | 3.99E-14 | *Sdhd, Trap1, Tubb2a, Tuba4a, Prkaca, Psmc6, Psma3, Ubb, Ndufa3, Ndufb3, Ndufv2, Psma6, Ndufb9, Ubc, Psma2, Casp3, Slc39a4, Psmd7, Tubb4b, Cox6b1, Psma4, Ndufa10, Mcu, Ube2l6, Slc11a2, Ndufa9, Atf6, Cox7a2l, Slc39a11, Uba1, Psmd12, Uqcrh, Psmd11, Psma8, Ndufb11, Uqcrc2, Psmd8, Ube2j1, Atp5pb, mt-Nd2, Ube2g1, Psmc3, Ndufs4, Ndufs1, Psmb2, Ndufs2, Slc18a1, Atp5g2, Ern1, Psma1, Vdac3, Slc25a5, Atp5b, Sdhc, Psmd2, Ndufa4, Sdhb, Maob, Slc39a9, mt-Co1, Eif2s1, Psmc5, Tuba1c, Prkacb, Ndufs8, Calm2, Atp5c1, mt-Nd5, Psmb7, Psmc2, Psmd14, Atp5a1, mt-Nd4, Eif2ak3, Gm10053, Itpr2, Slc18a2, Slc39a1, Ndufb5, Ndufv1, Calm3, Uqcrc1, Txn1, Xbp1, Psmd1, Psmc1, Hspa5, Sdha, mt-Cytb, Vdac1, Ube2g2, Psmd6, Psmc4, Ube2l3, Cox7a2, Psmb1, Psma5, Atp5j, Psmd13, Psmb4, Calm1, Cox5a, Pink1, Gnai3, Ndufv3, Casp9, Cycs, Ndufs6, Slc39a8, Ndufa1* |
| KEGG | Fatty acid metabolism | 40 | 6.67E-13 | *Acsl3, Hsd17b12, Acsl4, Acat1, Hadhb, Fasn, Acox3, Acsl5, Mcat, Scp2, Fads2, Acadvl, Scd1, Acat2, Acsl1, Cpt2, Hsd17b4, Acaa2, Hacd2, Elovl5, Acadsb, Acaa1b, Acadm, Echs1, Hadh, Elovl2, Acads, Hadha, Ppt1, Oxsm, Elovl6, Acat3, Acox1, Hacd1, Tecr, Hacd3, Elovl1, Elovl3, Fads1, Acsf3* |
| KEGG | Proteasome | 33 | 3.31E-12 | *Psma4, Psmd2, Psmd1, Psmc1, Psme2, Psmd12, Psmc6, Psma3, Psmd6, Psmc4, Psmd11, Psmc5, Psma8, Psme2b, Psmd8, Psmb1, Psme3, Psma5, Psmb4, Psmd13, Pomp, Psmb7, Psme4, Psma6, Psmc2, Psmc3, Psme1, Psmd14, Psma2, Psmb2, Psmd7, Psmb9, Psma1* |
| KEGG | Alzheimer disease | 137 | 7.48E-12 | *Sdhd, Fadd, Casp7, Tubb2a, Tuba4a, Chuk, Capn1, Psmc6, Psma3, Map2k1, Ndufa3, Nae1, Atg13, Ndufb3, Mapk1, Ndufv2, Psma6, Ndufb9, Fas, Psma2, Rtn4, Rb1cc1, Casp3, Slc39a4, Psmd7, Irs2, Tubb4b, Cox6b1, Atg101, Psma4, Ndufa10, Mcu, Irs1, Slc11a2, Ndufa9, Atf6, Ctnnb1, Cox7a2l, Slc39a11, Psmd12, Uqcrh, Psen1, Psmd11, Psma8, Ndufb11, Uqcrc2, Psmd8, Atp5pb, mt-Nd2, Raf1, Psmc3, Ndufs4, Ndufs1, Rtn3, Lrp5, Psmb2, Ndufs2, Atp5g2, Becn1, Ern1, Pik3c3, Psma1, Akt2, Vdac3, Slc25a5, Bace1, Atp5b, Sdhc, Psmd2, Ndufa4, Sdhb, Slc39a9, mt-Co1, Eif2s1, Fzd7, Psmc5, Tuba1c, Psenen, Ndufs8, Calm2, Ikbkg, Plcb1, Atp5c1, mt-Nd5, Psmb7, Atp2a2, Psmc2, Psmd14, Frat1, Atp5a1, mt-Nd4, Eif2ak3, Gm10053, Itpr2, Nox4, Il1b, Slc39a1, Ndufb5, Ndufv1, Calm3, Hras, Ulk2, Uqcrc1, Xbp1, Wipi2, Psmd1, Psmc1, Sdha, mt-Cytb, Vdac1, Psmd6, Capn2, Psmc4, Lrp6, Ppp3r1, Ide, Cox7a2, Pik3r4, Psmb1, Mtor, Psma5, Atp5j, Psmd13, Psen2, Mme, Psmb4, Pik3ca, Calm1, Cox5a, Ndufv3, Fzd5, Gsk3b, Casp9, Cycs, Ndufs6, Slc39a8, Ndufa1, Csnk2a1* |
| KEGG | Prion disease | 104 | 1.52E-11 | *Sdhd, Tubb2a, Tuba4a, Prkaca, Creb3, Psmc6, Psma3, C8a, Ndufa3, Ndufb3, Mapk1, Ndufv2, Psma6, Ndufb9, Cav1, Psma2, Casp3, Psmd7, Tubb4b, Cox6b1, Psma4, Ndufa10, Mcu, C6, Prnp, Ndufa9, Cox7a2l, C8b, Psmd12, C8g, Uqcrh, Psmd11, Psma8, Ndufb11, Uqcrc2, Psmd8, Atp5pb, Atf2, mt-Nd2, Psmc3, Ndufs4, Ndufs1, Psmb2, Ndufs2, Atp5g2, Hspa8, Psma1, Vdac3, Slc25a5, Atp5b, Sdhc, Psmd2, Ndufa4, Sdhb, mt-Co1, Eif2s1, Atf6b, Psmc5, Tuba1c, Prkacb, Ndufs8, Atp5c1, Hc, mt-Nd5, Psmb7, Psmc2, Psmd14, Atp5a1, mt-Nd4, Eif2ak3, Gm10053, Itpr2, Il1b, Ndufb5, Ndufv1, Uqcrc1, Psmd1, Psmc1, Hspa5, Sdha, Rac1, C9, mt-Cytb, Vdac1, Stip1, Psmd6, Psmc4, Hspa1b, Ppp3r1, Cox7a2, Psmb1, Psma5, Atp5j, Psmd13, Psmb4, Pik3ca, Cox5a, Ndufv3, Gsk3b, Casp9, Cycs, Creb3l3, Ndufs6, Ndufa1, Csnk2a1* |
| KEGG | Complement and coagulation cascades | 49 | 6.53E-11 | *Cfi, F8, Serpina1c, F13b, Cfh, C4bp, Plg, Mbl2, Serpine2, Serpind1, Cpb2, Serpina1b, C8a, Serpina1d, Serpinf2, Cfhr2, Cfhr1, Hc, C1ra, Masp2, Clu, F2, Cd59b, Serpinc1, Cr1l, Cd59a, C3, C6, Fga, F2r, C1rb, C8b, Cfb, C9, Vtn, C8g, Klkb1, C1s1, Serpina1e, F9, Fgb, Fgg, Serping1, F5, C4a, Mbl1, Serpina1a, F11, F10* |
| KEGG | Autophagy - animal | 64 | 1.67E-10 | *Akt2, Atg4b, Camkk2, Smcr8, Prkaca, Rheb, Snap29, Eif2s1, Rab1a, Ppp2ca, Map2k1, Atg4a, Uvrag, Gabarapl2, Rras2, Prkacb, Atg9a, Igbp1, Atg13, Cflar, Sqstm1, Traf6, Mras, Mapk1, Ppp2cb, Rb1cc1, Hif1a, Tank, Trp53inp2, Irs2, Eif2ak3, Atg101, Zfyve1, Rab7, Gabarapl1, Irs1, Ulk2, Hras, Pdpk1, Atg3, Wipi2, Atg5, Atg12, Pten, Prkaa1, Atg10, Ctsb, Lamp2, Pik3r4, Mtor, Atg4c, Pik3ca, Rab8a, Raf1, Prkaa2, Becn1, Deptor, Ern1, Pik3c3, Gabarap, Atg7, Lamp1, Rraga, Mtmr4* |
| KEGG | Pathways of neurodegeneration - multiple diseases | 155 | 3.97E-10 | *Sdhd, Trap1, Fadd, Casp7, Tubb2a, Tuba4a, Smcr8, Dnai1, Capn1, Derl1, Psmc6, Psma3, Map2k1, Ubb, Ndufa3, Bdnf, Atg13, Vcp, Sqstm1, Ndufb3, Mapk1, Ndufv2, Psma6, Actr1a, Ndufb9, Ubc, Fas, Psma2, Tank, Rb1cc1, Casp3, Psmd7, Sigmar1, Tubb4b, Cox6b1, Atg101, Fig4, Psma4, Ndufa10, Mcu, Zfyve1, Ube2l6, Prnp, Ndufa9, Als2, Atf6, Ctnnb1, Cox7a2l, Uba1, Psmd12, Uqcrh, Psen1, Psmd11, Psma8, Ndufb11, Uqcrc2, Psmd8, Ube2j1, Cat, Atp5pb, mt-Nd2, Ube2g1, Rab8a, Raf1, Psmc3, Ndufs4, Ndufs1, Lrp5, Psmb2, Ndufs2, Atp5g2, Becn1, Ern1, Pik3c3, Psma1, Rab5a, Vdac3, Slc25a5, Atp5b, Sdhc, Psmd2, Actr10, Ndufa4, Sdhb, mt-Co1, Eif2s1, Rab1a, Fzd7, Psmc5, Tuba1c, Ndufs8, Calm2, Plcb1, Atp5c1, mt-Nd5, Psmb7, Atp2a2, Atxn3, Psmc2, Atxn1, Psmd14, Frat1, Atp5a1, mt-Nd4, Atxn1l, Eif2ak3, Gm10053, Itpr2, Nox4, Vapb, Il1b, Ndufb5, Ndufv1, Calm3, Hras, Ulk2, Uqcrc1, Xbp1, Wipi2, Psmd1, Psmc1, Hspa5, Sdha, Rac1, mt-Cytb, Vdac1, Dctn4, Ube2g2, Psmd6, Capn2, Psmc4, Lrp6, Ppp3r1, Ube2l3, Cox7a2, Pik3r4, Psmb1, Mtor, Psma5, Atp5j, Psmd13, Psen2, Psmb4, Calm1, Gpx1, Cox5a, Pink1, Ndufv3, Fzd5, Gsk3b, Casp9, Cycs, Ndufs6, Dnah8, Ndufa1, Csnk2a1* |
| KEGG | Huntington disease | 108 | 1.62E-09 | *Sdhd, Tubb2a, Tuba4a, Dnai1, Creb3, Polr2k, Psmc6, Psma3, Ap2b1, Ndufa3, Bdnf, Atg13, Ndufb3, Ndufv2, Psma6, Actr1a, Ndufb9, Psma2, Cltc, Rb1cc1, Casp3, Psmd7, Tubb4b, Cox6b1, Atg101, Psma4, Ndufa10, Polr2b, Ndufa9, Cox7a2l, Psmd12, Uqcrh, Sod2, Psmd11, Psma8, Ndufb11, Uqcrc2, Psmd8, Atp5pb, mt-Nd2, Psmc3, Ndufs4, Ndufs1, Psmb2, Ndufs2, Atp5g2, Becn1, Ern1, Pik3c3, Tgm2, Psma1, Vdac3, Slc25a5, Atp5b, Sdhc, Psmd2, Actr10, Ndufa4, Sdhb, Ap2m1, mt-Co1, Ap2a2, Rcor1, Psmc5, Tuba1c, Ndufs8, Plcb1, Atp5c1, mt-Nd5, Psmb7, Psmc2, Psmd14, Atp5a1, mt-Nd4, Gm10053, Slc1a2, Ndufb5, Ndufv1, Sp1, Ulk2, Uqcrc1, Wipi2, Psmd1, Psmc1, Sdha, mt-Cytb, Vdac1, Dctn4, Psmd6, Psmc4, Cox7a2, Pik3r4, Psmb1, Mtor, Psma5, Atp5j, Psmd13, Psmb4, Clta, Gpx1, Cox5a, Ndufv3, Cltb, Casp9, Cycs, Creb3l3, Ndufs6, Dnah8, Ndufa1* |
| KEGG | Amyotrophic lateral sclerosis | 126 | 1.72E-09 | *Sdhd, Tubb2a, Tuba4a, Smcr8, Dnai1, Derl1, Psmc6, Psma3, Ndufa3, Atg13, Vcp, Sqstm1, Ncbp1, Ndufb3, Ndufv2, Psma6, Actr1a, Ndufb9, Ndc1, Pfn2, Psma2, Tank, Rb1cc1, Casp3, Psmd7, Nxt1, Sigmar1, Tubb4b, Cox6b1, Atg101, Fig4, Psma4, Ndufa10, Mcu, Nup50, Ndufa9, Als2, Atf6, Cox7a2l, Psmd12, Uqcrh, Anxa7, Psmd11, Psma8, Ndufb11, Uqcrc2, Erbb4, Psmd8, Cat, Atp5pb, mt-Nd2, Rab8a, Psmc3, Ndufs4, Ndufs1, Psmb2, Ang, Ndufs2, Atp5g2, Becn1, Ern1, Nup62, Pik3c3, Psma1, Rab5a, Atp5b, Sdhc, Psmd2, Actr10, Ndufa4, Sdhb, mt-Co1, Eif2s1, Rab1a, Nup37, Seh1l, Psmc5, Tuba1c, Sec13, Ndufs8, Atp5c1, mt-Nd5, Psmb7, Ubqln1, Psmc2, Psmd14, Atp5a1, Nxt2, mt-Nd4, Eif2ak3, Gm10053, Slc1a2, Vapb, Ndufb5, Ndufv1, Ulk2, Uqcrc1, Xbp1, Wipi2, Psmd1, Psmc1, Hspa5, Sdha, Rac1, mt-Cytb, Vdac1, Dctn4, Psmd6, Psmc4, Ppp3r1, Cox7a2, Pik3r4, Psmb1, Mtor, Psma5, Atp5j, Psmd13, Psmb4, Gpx1, Cox5a, Pink1, Ndufv3, Casp9, Cycs, Ndufs6, Dnah8, Ndufa1* |
| KEGG | Lysosome | 59 | 3.94E-09 | *Sumf1, Dnase2b, Gns, Asah1, Hexa, Lipa, Laptm4a, Cln5, Atp6v0d1, Mfsd8, Acp5, Glb1, Arsb, Gla, M6pr, Scarb2, Ppt1, Cltc, Ap3m1, Dnase2a, Smpd1, Ctsh, Tpp1, Manba, Ctsc, Atp6v1h, Ap3s1, Slc11a2, Atp6v0c, Ap3s2, Abca2, Gaa, Slc17a5, Gm2a, Igf2r, Gusb, Arsa, Ap1b1, Wdr7, Arsg, Atp6ap1, Ctsb, Lamp2, Ap1g1, Fuca1, Cd164, Acp2, Psap, Clta, Npc2, Sort1, Laptm4b, Ap3b1, Cltb, Ap4s1, Gnptg, Lamp1, Aga, Gga2* |
| KEGG | Valine, leucine and isoleucine degradation | 33 | 4.79E-09 | *Acat1, Aldh2, Hadhb, Abat, Hmgcs2, Aldh9a1, Bckdha, Hibadh, Dld, Acat2, Pcca, Aox3, Acaa2, Aldh7a1, Pccb, Acadsb, Acaa1b, Aacs, Acadm, Hmgcl, Echs1, Hadh, Hibch, Acads, Dbt, Hadha, Mmut, Hmgcs1, Acat3, Mccc2, Acad8, Mccc1, Acsf3* |
| KEGG | Terpenoid backbone biosynthesis | 19 | 5.65E-09 | *Mvk, Acat1, Idi1, Fdps, Dhdds, Fntb, Hmgcs2, Pmvk, Zmpste24, Mvd, Nus1, Hmgcr, Pcyox1, Acat2, Pdss2, Hmgcs1, Acat3, Icmt, Fnta* |
| KEGG | Autophagy - other | 23 | 6.50E-09 | *Atg9a, Atg4b, Igbp1, Atg13, Pik3r4, Mtor, Gabarapl1, Atg4c, Ulk2, Atg3, Ppp2cb, Wipi2, Atg5, Ppp2ca, Atg12, Becn1, Atg10, Pik3c3, Atg4a, Gabarap, Atg7, Atg101, Gabarapl2* |
| KEGG | Protein export | 21 | 2.98E-08 | *Srprb, Srp14, Sec61g, Srpr, Sec61a1, Srp68, Hspa5, Sec11c, Sec62, Sec11a, Oxa1l, Srp54b, Immp1l, Srp54a, Srp72, Sec63, Spcs1, Spcs3, Srp54c, Sec61b, Spcs2* |

Table S3. Enrichment analysis of 1,440 reverse V-shaped recovered genes in the 1st replicate

| **Category** | **Term** | **Count** | **PValue** | **Genes** |
| --- | --- | --- | --- | --- |
| GOBP | DNA repair | 53 | 1.24E-08 | *Sirt6, Dclre1a, Parp10, Ercc6l2, Rad9a, Uimc1, Zfp668, Hdgfl2, Mcrs1, Wdhd1, Timeless, Abraxas1, Xrcc1, Eme2, Poli, Recql4, Hdac10, Faap100, Kmt5c, Slx1b, Fancc, Alkbh2, Pold4, Wrap53, Cdk9, Apex2, Alkbh3, Hsf1, Ascc2, Ap5z1, Actr5, Helq, Rad52, Traip, Chd1l, Trim28, Mpg, Tfpt, Sirt7, Otub1, Endov, Kat5, Pold1, Rtel1, Msh5, Polm, Pwwp3a, Cep164, Fzr1, Ddb2, Rpain, Emsy, Ddx11* |
| GOBP | protein phosphorylation | 73 | 6.39E-08 | *Map3k11, Stk19, Limk1, Csf2ra, Clk2, Speg, Prkag2, Npr1, Cdkl3, Ngf, Brsk1, Ksr1, Map4k2, Plk1, Nek8, Ntrk1, Map4k1, Mok, Nek3, Pak6, Kit, Pim3, Prkag1, Ccl3, Stk25, Mast2, Pick1, Mapk7, Cdk9, Rps6kl1, Hsf1, Grk2, Map3k5, Fes, Clk3, Map3k15, Tex14, Nrbp2, Plk3, Phka2, Pkn3, Erbb2, Stk11, Trim28, Prkar1b, Cdk5, Trib2, Flt3l, Trpm6, Rara, Morc3, Pak4, Cdk11b, Mark4, Cdk10, Camk2b, Rorc, Cdc7, Camk1, Birc5, Tyk2, Cdk20, Nuak2, Mapk12, Trp53rkb, Tssk1, Rps6kb2, Fastk, Lrrk1, Mertk, Strada, Mapk11, Haspin* |
| GOBP | cellular response to DNA damage stimulus | 62 | 1.35E-07 | *Sirt6, Dclre1a, Parp10, Ercc6l2, Hmces, Rad9a, Brsk1, Uimc1, Nupr1l, Hdgfl2, Mcrs1, Timeless, Abraxas1, Xrcc1, Morc2a, Eme2, Poli, Hdac10, Faap100, Slx1b, Fancc, Brat1, Alkbh2, Pold4, Wrap53, Cdk9, Apex2, Flywch1, Alkbh3, Hsf1, Pidd1, Ascc2, Ap5z1, Actr5, Helq, Rad52, Traip, Plk3, Chd1l, Stk11, Mpg, Myc, Tfpt, Sirt7, Otub1, Kat5, Pold1, Rtel1, Phf1, Msh5, Zbtb40, Polm, Pwwp3a, Cep164, Bax, Fzr1, Fbxo5, Ddb2, Baz1b, Emsy, Ddx11, Cep63* |
| GOBP | chromatin organization | 47 | 4.88E-07 | *Usp21, Sirt6, Parp10, Epop, Trim28, Ring1, Cbx6, Banp, Chd7, Cbx8, Sirt7, Tada2a, Smarca2, Uimc1, Hira, Mcrs1, Kat5, Prmt7, Alkbh4, Prmt1, Pcgf2, Abraxas1, Ehmt2, Brd9, Cdan1, Phf1, Samd1, Hdac10, Kmt5c, Hdac6, Hdac3, Lrwd1, Hdac7, Hmg20b, Pwwp3a, Mrgbp, Tdrd3, Rnf40, Bap1, Baz1b, Rccd1, Phf20, Kat2a, Prdm9, Emsy, Smarcd2, Haspin* |
| GOBP | cell projection organization | 31 | 1.88E-06 | *Myo7a, Rpgr, Trappc14, Rab34, Ift43, Tmem138, Crocc, Ofd1, Cep250, Ift88, Entr1, Rabep2, Cep152, Cluap1, Wdr90, Bbs2, Prickle3, Mark4, Cdk10, Ocrl, Ttc12, Ccdc28b, Poc1a, Cep164, Tchp, Cep43, Cep131, Enkd1, Ccdc61, Lrrc56, Wtip* |
| GOBP | phosphorylation | 66 | 2.41E-06 | *Map3k11, Stk19, Pfkfb1, Limk1, Clk2, Speg, Adpgk, Dgka, Cdkl3, Brsk1, Ksr1, Map4k2, Plk1, Nek8, Pi4kb, Ntrk1, Taf1, Map4k1, Mok, Nek3, Pak6, Nmrk1, Kit, Ip6k1, Pim3, Stk25, Mast2, Shpk, Mapk7, Cdk9, Rps6kl1, Grk2, Map3k5, Fes, Clk3, Map3k15, Uckl1, Plk3, Pkn3, Erbb2, Tk1, Stk11, Cdk5, Trpm6, Pak4, Cdk11b, Mark4, Sphk1, Cdk10, Xylb, Camk2b, Cdc7, Camk1, Tyk2, Dgkz, Cdk20, Nuak2, Mapk12, Tssk1, Rps6kb2, Fastk, Lrrk1, Mertk, Baz1b, Mapk11, Haspin* |
| GOBP | cell cycle | 64 | 3.42E-05 | *Ambra1, Dclre1a, Anapc5, Haus5, Appl2, Crocc, Banp, Brsk1, Hjurp, Nupr1l, Rab11fip3, Klhdc8b, Entr1, Plk1, Timeless, Taf1, Nek3, Pim3, Rcc1, Mapk7, Apex2, E4f1, Ncapd2, Cep131, Cenpt, Ncapd3, Katnb1, Dctn1, Tex14, Plk3, Cntrob, Haus7, Stk11, Septin4, Sgsm3, Cdk5, Spag8, Syce2, Cep250, Ccnd3, Tacc3, Pak4, Cdk11b, Mark4, Cdc7, Camk1, Birc5, Septin1, Cdk20, Ing1, Haus1, Hmg20b, Cep164, Mapk12, Fzr1, Ddit3, Chtf18, Cdca3, Fbxo5, Spice1, Cables2, Strada, Haspin, Cep63* |
| GOBP | regulation of cell cycle | 36 | 4.43E-05 | *Fbxl8, Birc2, Commd5, Stk11, Zbtb17, Gadd45g, Myc, Yeats2, Tfpt, Sgsm3, Tada2a, Fbxl12, Mybbp1a, Nupr1l, Mcrs1, Sipa1, Smim22, Kat5, Ccnd3, Tacc3, Cdk11b, Irf1, Ccnl2, Cdk10, Birc5, Mrgbp, Cdk9, Bax, Ddit3, Fbxl6, Bap1, Cables2, Actr5, Prdm11, Mif, Kat2a* |
| GOBP | regulation of transcription from RNA polymerase II promoter | 131 | 5.43E-05 | *Cc2d1b, Gps2, Ambra1, Mamld1, Pias3, Klf1, Zbtb45, Zfp629, Tada2a, Zfp9, Zfp668, Zfp213, Pias4, Zfp865, Irf1, Twist1, Ldb1, Ppp1r13l, Tead3, Nfkb2, Zfp184, E4f1, Sox17, Zfp61, Zfp956, Meis3, Zfp623, Zfp787, Zfp786, Zfp653, Fbxl19, Hsf4, Pcgf6, Zfp457, Epop, Safb2, Myc, Six5, Smarca2, Med4, Lyl1, Kat5, Batf2, Irx3, Zfp970, Brd9, Prdm15, Tead4, Rorc, Zfp579, Zkscan5, Zfp143, Zbtb48, Zfp672, Csrnp2, Zfp444, Hand2, Tal1, Zfp27, Zfp821, Phf20, Smarcd2, Nfyc, Tbx2, Zfp94, Zfp788, Safb, Zfp523, Grhl2, Zfp846, Nupr1l, Zfp467, Hdgfl2, Zfp566, Hivep2, Zfp14, Cc2d1a, Med20, Zfp384, Med15, Zfp446, Ccnl2, Deaf1, 2810021J22Rik, Zfp28, Zfp683, Batf3, Zfp54, Zfp775, Mrgbp, Rbak, Nr1h2, Hsf1, Zfp691, Hoxb4, Rfx1, Zfp647, Zfp57, Glis2, Zfp524, Irf3, Zfp661, Tfeb, Zfp658, Sp100, Zbtb17, Zfp772, Zfp354a, Yeats2, Sirt7, Mllt10, Zfp777, Zfp934, Ehmt2, Zfp808, Sltm, Zbtb40, Zfp69, Zmiz2, Ddit3, Esrra, Zkscan3, Zfp1, Zscan2, Mafg, Zfp950, Zfp7, Kat2a, Maf, Tcf7l1, Tead2* |
| GOBP | mRNA processing | 40 | 1.59E-04 | *Sugp1, Cactin, Pan2, Esrp2, Ddx39a, Rbm5, Sfswap, Rbfox3, Thoc1, Zc3h13, Tsen54, Clasrp, Cd2bp2, Trmt2a, Zrsr1, Ess2, Tra2a, Akap17b, Lsm7, Prpf38b, Wdr83, Pdcd11, Cpsf4, Zcchc8, Rbm25, Arl6ip4, Tut1, Hsf1, Prpf40b, Lsm3, Tdrd3, Ddx39b, Sf3a1, Akap8l, Srrt, Phrf1, Sart1, Scnm1, U2af2, Snrpa1* |
| KEGG | Herpes simplex virus 1 infection | 41 | 9.99E-04 | *Birc2, Zfp94, Irf3, Zfp661, Zfp788, Zfp457, Sp100, Zfp658, Zfp772, Zfp354a, H2-T10, Zfp9, Zfp777, Card9, Zfp566, Zfp14, Zfp688, Nxf1, Pik3r2, Zfp808, Tyk2, Zfp28, H2-Q4, 2810021J22Rik, Zfp54, Bax, Zfp184, Rbak, Tab1, Zfp69, Zfp61, Zfp956, Zfp27, Zfp623, Zfp1, Zfp786, Zfp647, Zfp57, Zfp950, Zfp7, Tradd* |
| KEGG | MAPK signaling pathway | 29 | 0.002321676 | *Map3k11, Mapk8ip3, Flnb, Erbb2, Gadd45g, Ntf3, Myc, Vegfb, Arrb2, Ngf, Gm5741, Flt3l, Map4k2, Csf1, Rasgrp2, Ntrk1, Map4k1, Rasgrp4, Kit, Cacna1a, Cacnb3, Mapk7, Mapk12, Nfkb2, Tab1, Ddit3, Map3k5, Mapk11, Tradd* |
| KEGG | Glucagon signaling pathway | 14 | 0.002971155 | *Calml4, Pfkfb1, Cpt1c, Camk2b, Phka2, Pck2, Prkag1, Gys1, Prkag2, Plcb2, Prmt1, Phka1, G6pc3, Plcb4* |
| KEGG | Apoptosis | 16 | 0.00497934 | *Birc2, Birc5, Dffb, Gadd45g, Htra2, Septin4, Bax, Il3ra, Map3k5, Ddit3, Pidd1, Ngf, Ctsw, Pik3r2, Ntrk1, Tradd* |
| KEGG | Inflammatory mediator regulation of TRP channels | 15 | 0.006579195 | *Calml4, Camk2b, Adcy4, Trpv1, Mapk12, Ngf, Plcb2, Asic3, Trpv4, Plcg1, Mapk11, Pik3r2, Ntrk1, Plcb4, Cyp2j9* |
| KEGG | Calcium signaling pathway | 24 | 0.006977495 | *Calml4, Cacna1a, Sphk1, Camk2b, Camk1, Phka2, Smim6, Tfeb, Erbb2, Adcy4, Tpcn2, Nos3, Pde1b, Vegfb, Ngf, Ppif, Plcb2, Plcd1, Phka1, Tnnc1, Tbxa2r, Plcg1, Ntrk1, Plcb4* |
| KEGG | NOD-like receptor signaling pathway | 21 | 0.009471673 | *Pstpip1, Irf3, Birc2, Tyk2, Rbck1, Sharpin, Casp1, Mapk12, Mfn1, Tab1, Naip5, Dnm1l, Plcb2, Card9, Rnf31, Nod1, Mapk11, Casp4, Plcb4, Gpsm3, Nlrp6* |
| KEGG | Lipid and atherosclerosis | 21 | 0.009471673 | *Calml4, Irf3, Camk2b, Ccl3, Casp1, Nos3, Mapk12, Bax, Arhgef1, Tab1, Map3k5, Ddit3, Ager, Plcb2, Mib2, Plcg1, Mapk11, Pik3r2, Ncf4, Plcb4, Cyp2j9* |
| KEGG | Phosphatidylinositol signaling system | 12 | 0.011493092 | *Calml4, Ocrl, Ip6k1, Plcb2, Dgkz, Inppl1, Plcd1, Plcg1, Pi4kb, Pik3r2, Dgka, Plcb4* |
| KEGG | PI3K-Akt signaling pathway | 30 | 0.013274574 | *Itgb7, Col9a3, Gnb2, Pkn3, Gys1, Erbb2, Stk11, Ntf3, Pik3r6, Myc, Epor, Il3ra, Vegfb, Ngf, Gm5741, Flt3l, Csf1, Ccnd3, Lama5, Pik3r2, Ntrk1, Ppp2r3d, Kit, Pck2, Vwf, Itga7, Nos3, Rps6kb2, G6pc3, Gngt2* |

Table S4. Enrichment analysis of 1,574 V-shaped recovered genes in the 1st replicate

| Category | Term | Count | PValue | Genes |
| --- | --- | --- | --- | --- |
| GOBP | protein transport | 107 | 1.48333E-18 | *Ran, Arfgap2, Arf1, Derl1, Rabgap1l, Arf6, Snx18, Plekhf2, Snap29, Sec23a, Rab6a, Nsf, Snx4, Pgap1, Ccdc91, Atg9a, Stxbp3, Hook1, Washc5, Snx24, Snx9, Nbas, Stam, Tmed2, Cadps2, Vps26b, Cmtm6, Cog5, Vps35, Sft2d2, Rab5c, Sec61a1, Ap3s2, Ipo7, Lman1, Rbsn, Rab5b, Washc4, Kpna1, Exoc4, Eps15, Arcn1, Dennd10, Rab1b, Rab10, Yif1a, Ap5m1, Timm29, Tmed10, Scfd1, Uevld, Appbp2, Copg2, Arfgef1, Rab2a, Vps37a, Copb1, Rab5a, Vps45, Bbs9, Ipo8, Golph3, Vps41, Rab18, Cse1l, Tnpo1, Dag1, Vps4b, Seh1l, Yif1b, Arl8b, Lin7c, Snap23, Vps29, Stam2, Snx27, Rab14, Rab21, Ppt1, Aftph, Tbc1d5, Chmp5, Nxt2, Chmp3, Vps13b, Arf3, Timm23, Rab7, Snx7, Snx25, Pex7, Vps54, Atg10, Sec63, Necap1, 2610002M06Rik, Kpna4, Snx13, Psen2, Zmat3, Stx12, Vps26a, Ap3b1, Scamp1, Tnks, Rab38, Copb2* |
| GOBP | intracellular protein transport | 65 | 7.60991E-17 | *Vps45, Arf1, Srpr, Ipo8, Vps41, 5730455P16Rik, Rabgap1l, Arf6, Rab18, Cse1l, Sec23a, Sec24d, Rab6a, Tnpo1, Nsf, Exph5, Tbc1d9b, Stxbp3, Vps29, Tbc1d12, Snx27, Rab14, Rab21, M6pr, Snx9, Tbc1d8b, Tbc1d5, Cltc, Tmed2, Tbc1d30, Arl5b, Exoc6b, Vps26b, Ric1, Arf3, Sytl4, Vps35, Timm23, Rab7, Chm, Rab5c, Ap3s2, Ipo7, Rab5b, Tbc1d22a, Tbck, Arl1, Snx13, Tmed7, Rab1b, Rab10, Stx12, Ngfr, Vps26a, Ap3b1, Arl5a, Tmed10, Scfd1, Appbp2, Copg2, Rab2a, Rab38, Copb1, Copb2, Rab5a* |
| GOBP | lipid metabolic process | 97 | 6.38274E-11 | *Acat1, Ugt2b5, Pik3c2a, Acpp, Erlin1, Asah1, Insig2, Lipa, Ptpn11, Ugt1a9, Mtmr2, Pafah1b1, Srd5a2, Cyp2c29, Acss3, Echs1, Hsd17b7, Pcx, Hadha, Lypla1, Ugt1a1, Serinc5, Acsm3, Ephx1, Serinc1, Fads1, Cyp8b1, Plcxd2, Crkl, Hsd17b12, Hmgcs2, Erg28, Samd8, Acad11, Scp2, Pcca, Apob, Dhrs9, Hmgcl, Lactb, Lima1, Hadh, Pcyt1a, Mttp, Elovl2, Agmo, Degs1, Daglb, Acsl4, Acsl3, Dhdds, Hadhb, Atp5b, Acsl5, Srd5a1, Acox2, Acadvl, Hsd17b4, Ptdss1, Mboat7, Zadh2, Ptgr2, Acadsb, Gla, Pafah1b2, Plcb1, Acadm, Gpld1, Ttc39b, Atp5a1, Lrat, C3, Dhcr24, Rab7, Crot, Akr1c6, Lpgat1, Enpp2, Scd1, Acsl1, Pten, Hdlbp, Slc16a1, Elovl5, Sgms2, Adipor2, Etnk1, Erlin2, Sacm1l, Cyp2c40, Rdh10, Rdh11, Prkaa2, Pnpla8, Acly, Decr2, Lclat1* |
| GOBP | vesicle-mediated transport | 49 | 9.15427E-10 | *Vps45, Arfgap2, Arf1, Vps41, Arf6, Sec23a, Rab6a, Nsf, Bloc1s5, Tvp23b, Ston1, Stxbp3, Snap23, Clint1, Tbc1d8b, Cltc, Kif1b, Tmed2, Arl5b, Syt1, Vps13b, Arf3, Kif3b, Sft2d2, Chm, Yipf2, Ap3s2, Ston2, Lman1, Necap1, Rab30, Cul3, Arl1, Arcn1, Ergic1, Yif1a, Rab10, Stx12, Ap3b1, Ap5m1, Arl5a, Rtn3, Tmed10, Scfd1, Copg2, Rab2a, Rab38, Copb1, Copb2* |
| GOBP | hemostasis | 18 | 3.44866E-09 | *Serpina10, Fgg, Tfpi2, F8, Fga, F2r, F13b, Serping1, Enpp4, Plg, Anxa5, F5, F2, Serpind1, Serpinc1, Cpb2, F9, Fgb* |
| GOBP | blood coagulation | 23 | 5.13742E-09 | *Bloc1s3, Serpina10, Fgg, Tfpi2, F8, C3, Fga, Habp2, F2r, F13b, Serping1, Enpp4, Plg, Anxa5, F5, F2, Ap3b1, Serpind1, C9, Serpinc1, Cpb2, F9, Fgb* |
| GOBP | protein polyubiquitination | 35 | 1.35453E-07 | *Amfr, Rnf41, Arih1, Ctnnb1, Trim2, Ube2e1, Rc3h1, Mkrn2, Rnf217, Ube2r2, Rnf152, Trim14, Ube2q2, Ube2h, Wsb2, Cul3, Ube2j1, Rnf19b, Traf6, Fbxw11, Ube2v2, Nhlrc3, Rmnd5a, Fbxl17, Ankib1, Ube2u, Marchf8, Tnks2, Peli2, Smurf2, Ube3c, Tnks, Trim32, Hectd2, Rnf125* |
| GOBP | fatty acid beta-oxidation | 18 | 1.62955E-07 | *Acat1, Hadhb, Acadm, Echs1, Acad9, Hadh, Abcd3, Crot, Acad11, Hadha, Acox2, Scp2, Pex7, Acadvl, Acat3, Echdc1, Hsd17b4, Acadsb* |
| GOBP | ubiquitin-dependent protein catabolic process | 43 | 7.80103E-07 | *Amfr, Usp9x, Arih1, Ddb1, Rnf149, Herc4, Ubr3, Rc3h1, Mkrn2, Uba1, Usp25, Rnf217, Ube2r2, Fem1c, Usp46, Ube2h, Psmd11, Fbxw8, Fbxo8, Psma8, Rnf13, Uba6, Usp12, Cul3, Rnf19b, Mib1, Ndfip2, Rnf11, Psmd13, Atxn3, Rnf130, Ubqln1, Ankib1, Smurf2, Ube3c, Cops3, Trim32, Nploc4, Hectd1, Rnf125, Psma1, Rnf146, Rnf5* |
| GOBP | ubiquitin-dependent ERAD pathway | 20 | 1.05258E-06 | *Dnajb9, Amfr, Erlin1, Ube2j1, Erlin2, Ccdc47, Derl1, Erlec1, Sgtb, Ubqln1, Hsp90b1, Canx, Man1a, Marchf6, Calr, Edem1, Stt3b, Nploc4, Rnf103, Rnf5* |
| KEGG | Metabolic pathways | 210 | 8.66945E-16 | *Uox, Ugt2b1, H6pd, Alg6, Asah1, Glud1, St6gal1, Adk, Suclg1, Mtmr2, Srd5a2, Rxylt1, Pgap1, Pigt, Gsr, Acss3, Pigg, Csad, Echs1, Hsd17b7, Rpn2, Enpp4, Anpep, Car8, Acsm3, Stt3b, Pgam1, Fads1, Cyp8b1, Prdm2, Ndst1, Hsd17b12, Aldh5a1, Abat, Ugt2b38, Hmgcs2, Atp6v0c, Bckdha, Cpox, Scp2, Camkmt, C1galt1, Car14, 9130409I23Rik, Pdxk, Mocos, Papss1, Ppat, Dhrs9, Pcyt1a, Hadh, Paics, Tymp, Degs1, Cyp2j5, Acsl4, Gfpt1, Hadhb, Atp5b, Acsl5, Fbp1, Rpe, Srd5a1, Hsd17b4, Mat2a, Csgalnact2, Acadsb, Gla, Pafah1b2, Nadk, Plcb1, Galnt1, mt-Nd5, Gpld1, Ppt1, Kmo, Mgat5, Stt3a, Idi1, Tpk1, Enpp2, Dpys, Sdha, Carnmt1, Elovl5, Pgk1, G6pc, Etnk1, Sacm1l, Cyp2c40, Nme7, Cbs, Rdh10, Pgam2, Rdh11, Rfk, Gch1, Bpnt2, Pank3, Alg14, Dpyd, Acly, Lclat1, Acat1, Ugt2b5, Pik3c2a, Ddc, Vkorc1l1, Cps1, Ldha, Ugt1a9, Pafah1b1, L2hgdh, Amt, Gcnt4, Adi1, Enpp3, Arsb, Atp6v1a, Cyp2c29, Pcx, Sucla2, Dbt, Hadha, Kyat3, Prps2, Suox, Bbox1, Pde7b, Ugt1a1, Prune1, Alg10b, Pdha1, Uxs1, Cyp3a11, Otc, Txndc12, Pcca, Crppa, Thtpa, Uqcrc2, Glt28d2, Glce, Hmgcl, Atp5pb, Elovl2, Lap3, Pank1, Prps1, Acsl3, Gns, Eno1b, Acox2, Gart, Maob, Fech, Pklr, Dld, Acadvl, Car5a, Gphn, Ptdss1, Pccb, Atic, Acadm, Tdo2, Extl2, Gmpr2, Pygl, Mmut, Atp5a1, Tkfc, Ugt1a6b, Ido2, Hmox2, Smyd1, Aldh2, Gclm, Lrat, Dhcr24, C1galt1c1, Srr, Kynu, Gstm7, Mgat2, Hnmt, Hgd, Scd1, Acsl1, Echdc1, Man1a, Sqor, Pten, Aldh7a1, Gbe1, Sgms2, Man1a2, Hao1, Suclg2, Tgds, Ugt1a5, Nadk2, Mthfd1, Mgam, Bpnt1, Acat3, Eprs, Mccc1, Urad, Chdh* |
| KEGG | Complement and coagulation cascades | 30 | 4.31408E-11 | *Cfi, F8, C3, Fga, F2r, F13b, Cfh, C4bp, Plg, C8b, Mbl2, Serpind1, Cfb, C9, Cpb2, Serpina1b, Serpina1d, Serpinf2, Serpina1e, F9, Fgb, Cfhr1, Fgg, Masp2, Serping1, F5, Clu, F2, Serpinc1, Cr1l* |
| KEGG | Protein processing in endoplasmic reticulum | 41 | 3.00512E-10 | *Amfr, Sec61a1, Atf6, Derl1, Rad23b, Eif2ak1, Hsp90b1, Lman1, Eif2s1, Canx, Hspa4l, Sec23a, Man1a, Sec24d, Sec63, Edem1, Man1a2, Ube2j1, Pdia3, Rpn2, Pdia6, Atxn3, Erlec1, Ubqln1, Bag2, Svip, Dnajc3, Ngly1, Ssr3, Pdia4, Hsp90ab1, Ubxn2a, Rad23a, Ostc, Marchf6, Calr, Stt3b, Nploc4, Ubxn6, Stt3a, Rnf5* |
| KEGG | Propanoate metabolism | 15 | 1.94054E-08 | *Acss3, Suclg2, Abat, Echs1, Bckdha, Sucla2, Dbt, Hadha, Ldha, Dld, Mmut, Suclg1, Pcca, Echdc1, Pccb* |
| KEGG | Fatty acid metabolism | 21 | 1.97288E-08 | *Acsl4, Acat1, Acsl3, Hsd17b12, Hadhb, Acadm, Acsl5, Echs1, Hadh, Elovl2, Hadha, Scp2, Ppt1, Acadvl, Scd1, Acsl1, Acat3, Hsd17b4, Elovl5, Fads1, Acadsb* |
| KEGG | Valine, leucine and isoleucine degradation | 20 | 2.4393E-08 | *Acat1, Hadhb, Aldh2, Abat, Acadm, Hmgcl, Hmgcs2, Echs1, Hadh, Bckdha, Dbt, Hadha, Dld, Mmut, Pcca, Acat3, Mccc1, Aldh7a1, Pccb, Acadsb* |
| KEGG | Carbon metabolism | 28 | 4.03416E-07 | *Pdha1, Prps1, Acat1, H6pd, Fbp1, Rpe, Cps1, Eno1b, Glud1, Pklr, Sdha, Dld, Suclg1, Pcca, Amt, Pccb, Pgk1, Hao1, Suclg2, Echs1, Pcx, Sucla2, Pgam2, Prps2, Mmut, Acat3, Tkfc, Pgam1* |
| KEGG | Biosynthesis of cofactors | 32 | 4.77679E-07 | *Pank1, Ugt2b5, Aldh2, Ugt2b1, Gclm, Ugt2b38, Vkorc1l1, Tpk1, Kynu, Cpox, Fech, Ugt1a9, Dld, Gphn, Mat2a, Pdxk, Nadk, Tdo2, Ugt1a5, Nme7, Nadk2, Mthfd1, Rdh11, Gch1, Rfk, Eprs, Ugt1a1, Kmo, Ugt1a6b, Pank3, Ido2, Ggh* |
| KEGG | Endocytosis | 45 | 2.06242E-06 | *Vps35, Arpc5, Actr3, Rnf41, Vps45, Arfgap2, Arf1, Rab7, Smad2, Rab5c, Cdc42, Igf2r, Arf6, Pard3, Rbsn, Rab5b, Washc4, Asap2, Vps4b, Snx4, Psd3, 2610002M06Rik, Eps15, Smap1, Traf6, Vps29, Rhoa, Washc5, Actr2, Stam2, Rab10, Vps26a, Arpc2, Chmp5, Cltc, Smurf2, Arfgef1, Stam, Capza2, Arpc4, Vps26b, Vps37a, Chmp3, Arf3, Rab5a* |
| KEGG | Autophagy - animal | 29 | 3.43307E-06 | *Rab7, Gabarapl1, Irs1, Smcr8, Prkaca, Pdpk1, Atg5, Snap29, Eif2s1, Atg12, Map2k1, Pten, Atg10, Ctsb, Lamp2, Prkacb, Atg9a, Sqstm1, Traf6, Mapk1, Raf1, Prkaa2, Rb1cc1, Hif1a, Tank, Deptor, Irs2, Lamp1, Rraga* |
