## Supplementary material for "A potential pathway that links intron retention with the physiological recovery by a Japanese herbal medicine": Fig. S1 - S16

### Slide 1
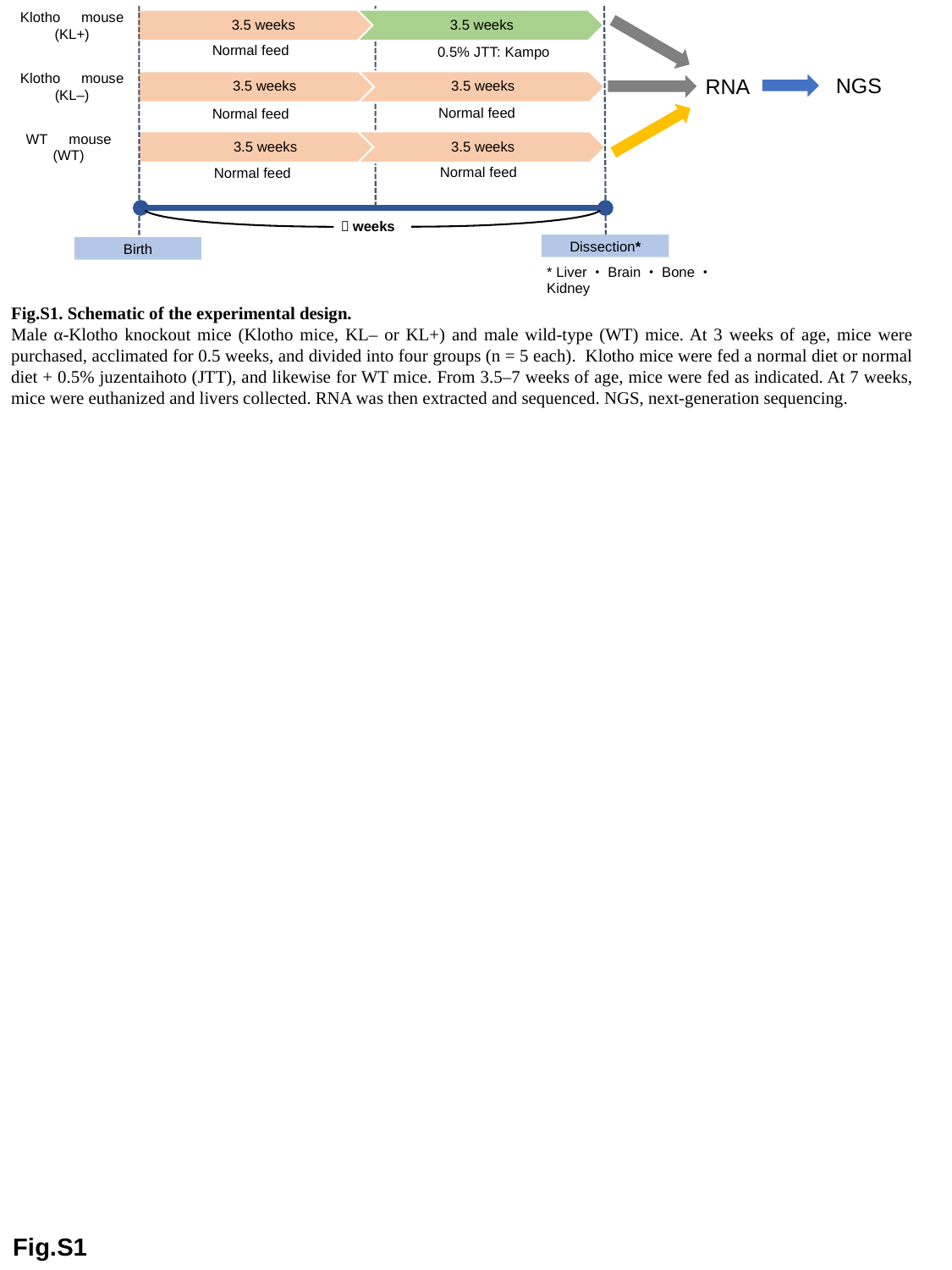

Klotho　mouse
(KL+)
Normal feed
0.5% JTT: Kampo
Klotho　mouse
(KL–)
NGS
RNA
Normal feed
Normal feed
WT　mouse
(WT)
Normal feed
Normal feed
７weeks
Dissection*
Birth
* Liver・Brain・Bone・Kidney
Fig.S1. Schematic of the experimental design.
Male α-Klotho knockout mice (Klotho mice, KL– or KL+) and male wild-type (WT) mice. At 3 weeks of age, mice were purchased, acclimated for 0.5 weeks, and divided into four groups (n = 5 each). Klotho mice were fed a normal diet or normal diet + 0.5% juzentaihoto (JTT), and likewise for WT mice. From 3.5–7 weeks of age, mice were fed as indicated. At 7 weeks, mice were euthanized and livers collected. RNA was then extracted and sequenced. NGS, next-generation sequencing.
Fig.S1

### Slide 2
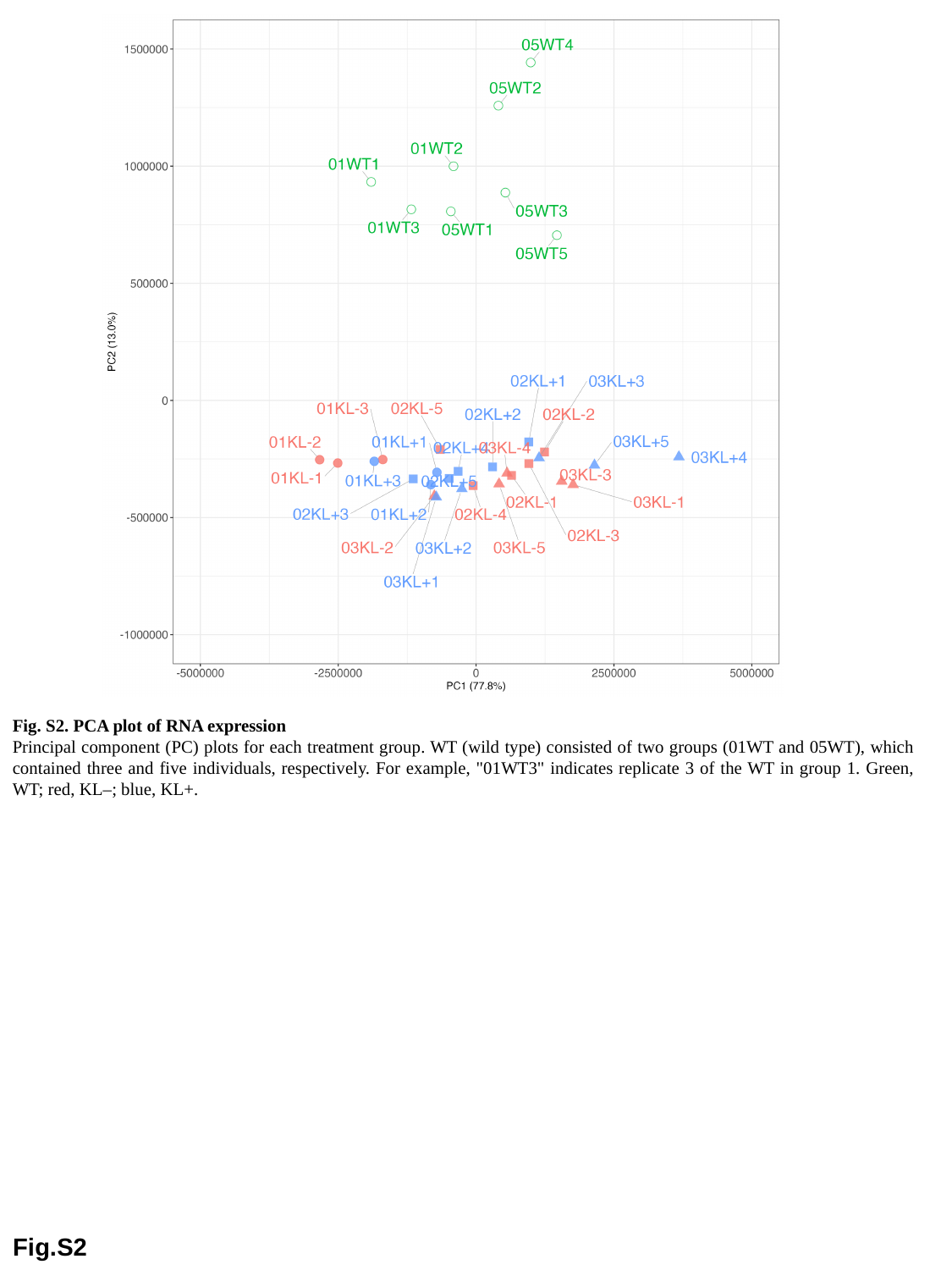

Fig. S2. PCA plot of RNA expression
Principal component (PC) plots for each treatment group. WT (wild type) consisted of two groups (01WT and 05WT), which contained three and five individuals, respectively. For example, "01WT3" indicates replicate 3 of the WT in group 1. Green, WT; red, KL–; blue, KL+.
Fig.S2

### Slide 3
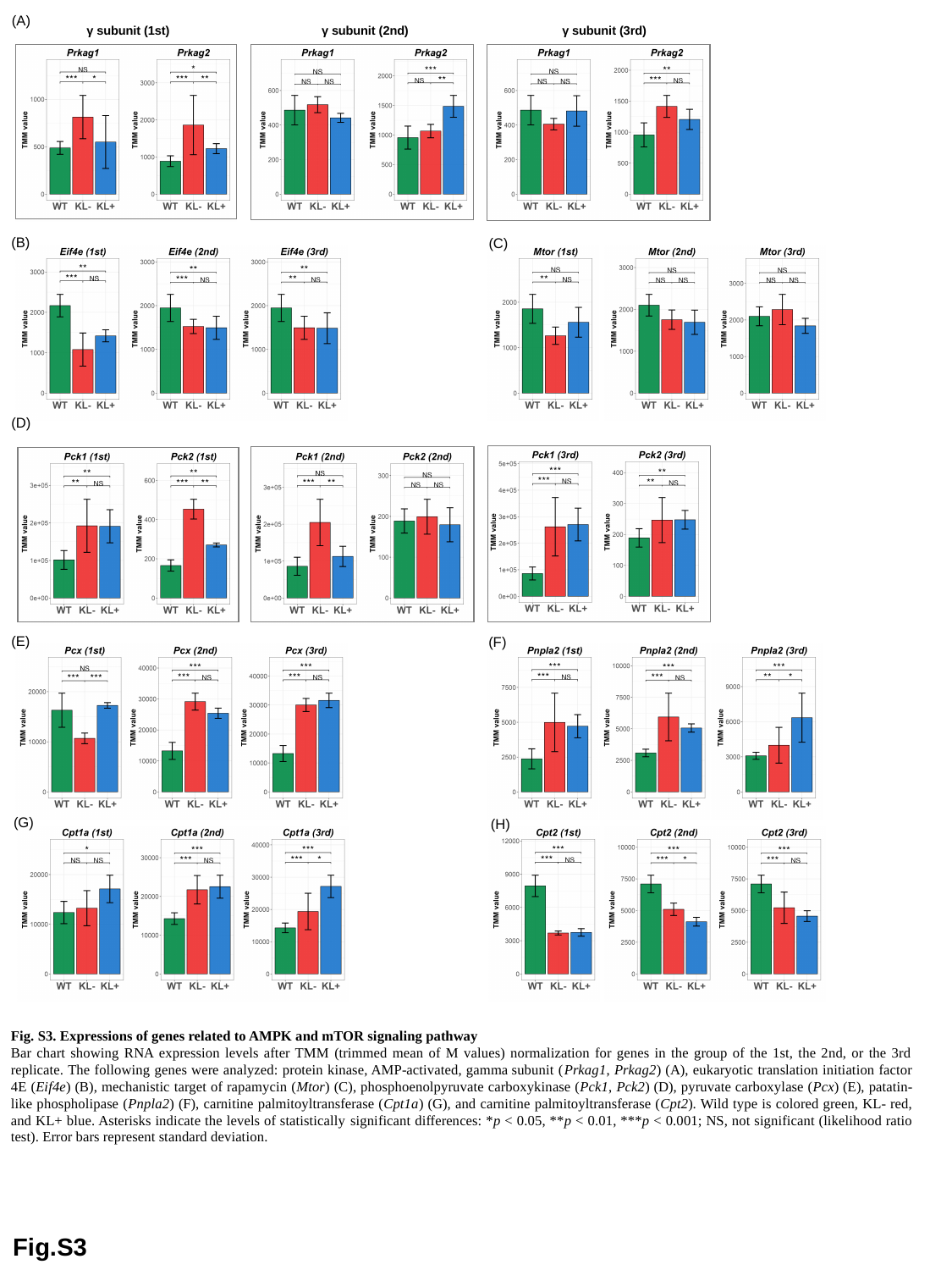

(A)
γ subunit (1st)
γ subunit (2nd)
γ subunit (3rd)
(B)
(C)
(D)
(E)
(F)
(G)
(H)
Fig. S3. Expressions of genes related to AMPK and mTOR signaling pathway
Bar chart showing RNA expression levels after TMM (trimmed mean of M values) normalization for genes in the group of the 1st, the 2nd, or the 3rd replicate. The following genes were analyzed: protein kinase, AMP-activated, gamma subunit (Prkag1, Prkag2) (A), eukaryotic translation initiation factor 4E (Eif4e) (B), mechanistic target of rapamycin (Mtor) (C), phosphoenolpyruvate carboxykinase (Pck1, Pck2) (D), pyruvate carboxylase (Pcx) (E), patatin-like phospholipase (Pnpla2) (F), carnitine palmitoyltransferase (Cpt1a) (G), and carnitine palmitoyltransferase (Cpt2). Wild type is colored green, KL- red, and KL+ blue. Asterisks indicate the levels of statistically significant differences: *p < 0.05, **p < 0.01, ***p < 0.001; NS, not significant (likelihood ratio test). Error bars represent standard deviation.
Fig.S3

### Slide 4
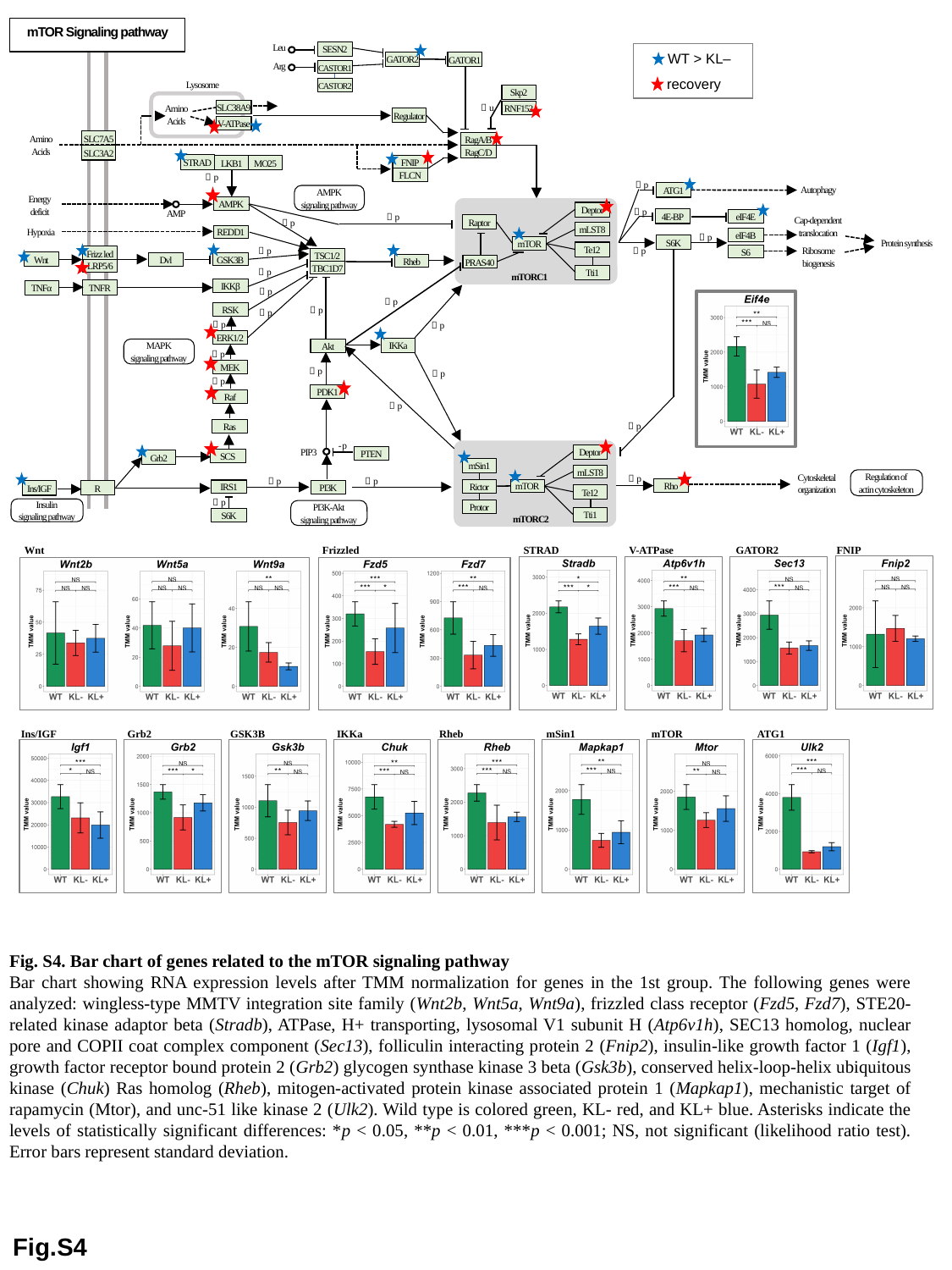

mTOR Signaling pathway
SESN2
Leu
GATOR2
GATOR1
CASTOR1
Arg
CASTOR2
Lysosome
Skp2
SLC38A9
RNF152
＋u
Amino
Acids
Regulator
V-ATPase
SLC7A5
RagA/B
Amino
Acids
RagC/D
SLC3A2
STRAD
LKB1
MO25
FNIP
FLCN
＋p
Autophagy
＋p
ATG1
AMPK
signaling pathway
Energy
deficit
AMPK
Deptor
＋p
AMP
4E-BP
eIF4E
＋p
Raptor
Cap-dependent
translocation
＋p
mLST8
REDD1
Hypoxia
eIF4B
＋p
S6K
Protein synthesis
mTOR
Te12
S6
＋p
＋p
Frizz led
Ribosome
biogenesis
TSC1/2
Wnt
GSK3B
Dvl
Rheb
PRAS40
LRP5/6
TBC1D7
Tti1
＋p
mTORC1
IKKβ
TNFR
TNFα
＋p
#### Chart
| Category |
|---|＋p
RSK
＋p
＋p
＋p
＋p
ERK1/2
IKKa
MAPK
signaling pathway
Akt
＋p
MEK
＋p
＋p
＋p
PDK1
Raf
＋p
Ras
＋p
- p
Deptor
PTEN
PIP3
SCS
Grb2
mSin1
Rictor
Protor
mLST8
Regulation of
actin cytoskeleton
Cytoskeletal
organization
＋p
＋p
＋p
Rho
mTOR
IRS1
PI3K
R
Ins/IGF
Te12
＋p
Insulin
signaling pathway
PI3K-Akt
signaling pathway
Tti1
S6K
mTORC2
WT > KL–
recovery
Wnt
Frizzled
STRAD
V-ATPase
GATOR2
FNIP
Ins/IGF
Grb2
GSK3B
IKKa
Rheb
mSin1
mTOR
ATG1
Fig. S4. Bar chart of genes related to the mTOR signaling pathway
Bar chart showing RNA expression levels after TMM normalization for genes in the 1st group. The following genes were analyzed: wingless-type MMTV integration site family (Wnt2b, Wnt5a, Wnt9a), frizzled class receptor (Fzd5, Fzd7), STE20-related kinase adaptor beta (Stradb), ATPase, H+ transporting, lysosomal V1 subunit H (Atp6v1h), SEC13 homolog, nuclear pore and COPII coat complex component (Sec13), folliculin interacting protein 2 (Fnip2), insulin-like growth factor 1 (Igf1), growth factor receptor bound protein 2 (Grb2) glycogen synthase kinase 3 beta (Gsk3b), conserved helix-loop-helix ubiquitous kinase (Chuk) Ras homolog (Rheb), mitogen-activated protein kinase associated protein 1 (Mapkap1), mechanistic target of rapamycin (Mtor), and unc-51 like kinase 2 (Ulk2). Wild type is colored green, KL- red, and KL+ blue. Asterisks indicate the levels of statistically significant differences: *p < 0.05, **p < 0.01, ***p < 0.001; NS, not significant (likelihood ratio test). Error bars represent standard deviation.
Fig.S4

### Slide 5
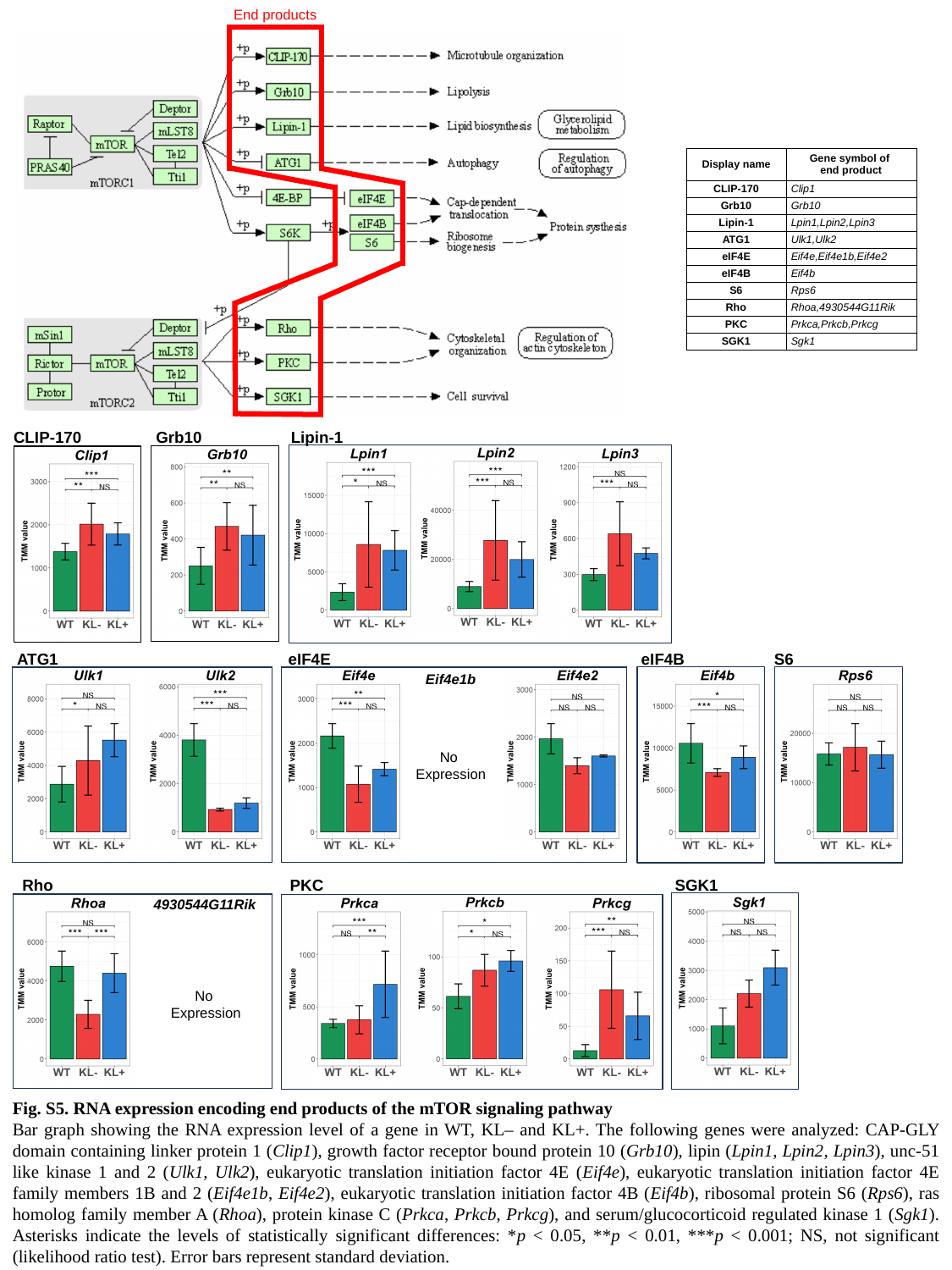

End products
| Display name | Gene symbol of end product |
| --- | --- |
| CLIP-170 | Clip1 |
| Grb10 | Grb10 |
| Lipin-1 | Lpin1,Lpin2,Lpin3 |
| ATG1 | Ulk1,Ulk2 |
| eIF4E | Eif4e,Eif4e1b,Eif4e2 |
| eIF4B | Eif4b |
| S6 | Rps6 |
| Rho | Rhoa,4930544G11Rik |
| PKC | Prkca,Prkcb,Prkcg |
| SGK1 | Sgk1 |
CLIP-170
Grb10
Lipin-1
ATG1
eIF4E
eIF4B
S6
Eif4e1b
No
Expression
Rho
PKC
SGK1
4930544G11Rik
No
Expression
Fig. S5. RNA expression encoding end products of the mTOR signaling pathway
Bar graph showing the RNA expression level of a gene in WT, KL– and KL+. The following genes were analyzed: CAP-GLY domain containing linker protein 1 (Clip1), growth factor receptor bound protein 10 (Grb10), lipin (Lpin1, Lpin2, Lpin3), unc-51 like kinase 1 and 2 (Ulk1, Ulk2), eukaryotic translation initiation factor 4E (Eif4e), eukaryotic translation initiation factor 4E family members 1B and 2 (Eif4e1b, Eif4e2), eukaryotic translation initiation factor 4B (Eif4b), ribosomal protein S6 (Rps6), ras homolog family member A (Rhoa), protein kinase C (Prkca, Prkcb, Prkcg), and serum/glucocorticoid regulated kinase 1 (Sgk1). Asterisks indicate the levels of statistically significant differences: *p < 0.05, **p < 0.01, ***p < 0.001; NS, not significant (likelihood ratio test). Error bars represent standard deviation.

### Slide 6
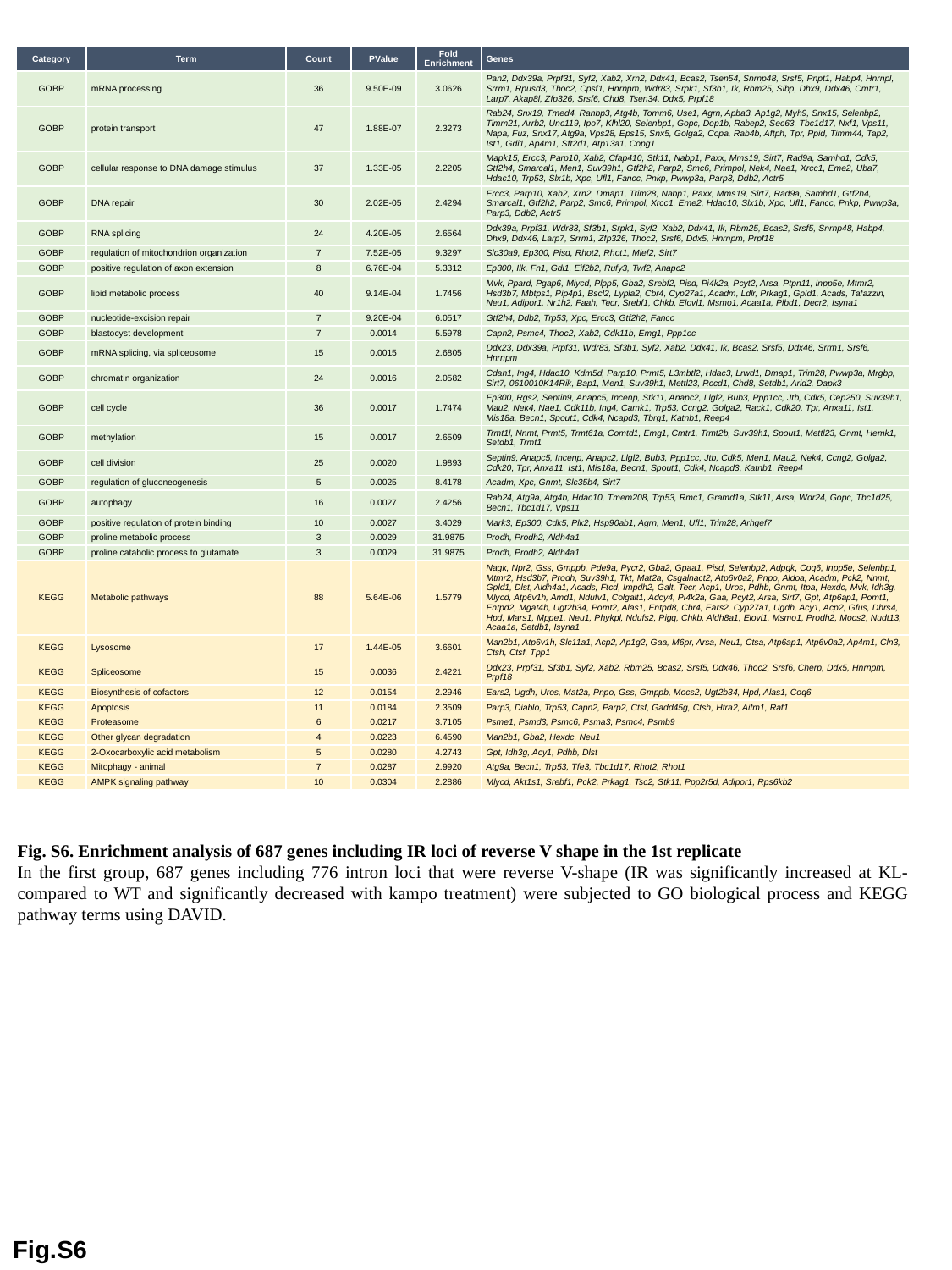

| Category | Term | Count | PValue | Fold Enrichment | Genes |
| --- | --- | --- | --- | --- | --- |
| GOBP | mRNA processing | 36 | 9.50E-09 | 3.0626 | Pan2, Ddx39a, Prpf31, Syf2, Xab2, Xrn2, Ddx41, Bcas2, Tsen54, Snrnp48, Srsf5, Pnpt1, Habp4, Hnrnpl, Srrm1, Rpusd3, Thoc2, Cpsf1, Hnrnpm, Wdr83, Srpk1, Sf3b1, Ik, Rbm25, Slbp, Dhx9, Ddx46, Cmtr1, Larp7, Akap8l, Zfp326, Srsf6, Chd8, Tsen34, Ddx5, Prpf18 |
| GOBP | protein transport | 47 | 1.88E-07 | 2.3273 | Rab24, Snx19, Tmed4, Ranbp3, Atg4b, Tomm6, Use1, Agrn, Apba3, Ap1g2, Myh9, Snx15, Selenbp2, Timm21, Arrb2, Unc119, Ipo7, Klhl20, Selenbp1, Gopc, Dop1b, Rabep2, Sec63, Tbc1d17, Nxf1, Vps11, Napa, Fuz, Snx17, Atg9a, Vps28, Eps15, Snx5, Golga2, Copa, Rab4b, Aftph, Tpr, Ppid, Timm44, Tap2, Ist1, Gdi1, Ap4m1, Sft2d1, Atp13a1, Copg1 |
| GOBP | cellular response to DNA damage stimulus | 37 | 1.33E-05 | 2.2205 | Mapk15, Ercc3, Parp10, Xab2, Cfap410, Stk11, Nabp1, Paxx, Mms19, Sirt7, Rad9a, Samhd1, Cdk5, Gtf2h4, Smarcal1, Men1, Suv39h1, Gtf2h2, Parp2, Smc6, Primpol, Nek4, Nae1, Xrcc1, Eme2, Uba7, Hdac10, Trp53, Slx1b, Xpc, Ufl1, Fancc, Pnkp, Pwwp3a, Parp3, Ddb2, Actr5 |
| GOBP | DNA repair | 30 | 2.02E-05 | 2.4294 | Ercc3, Parp10, Xab2, Xrn2, Dmap1, Trim28, Nabp1, Paxx, Mms19, Sirt7, Rad9a, Samhd1, Gtf2h4, Smarcal1, Gtf2h2, Parp2, Smc6, Primpol, Xrcc1, Eme2, Hdac10, Slx1b, Xpc, Ufl1, Fancc, Pnkp, Pwwp3a, Parp3, Ddb2, Actr5 |
| GOBP | RNA splicing | 24 | 4.20E-05 | 2.6564 | Ddx39a, Prpf31, Wdr83, Sf3b1, Srpk1, Syf2, Xab2, Ddx41, Ik, Rbm25, Bcas2, Srsf5, Snrnp48, Habp4, Dhx9, Ddx46, Larp7, Srrm1, Zfp326, Thoc2, Srsf6, Ddx5, Hnrnpm, Prpf18 |
| GOBP | regulation of mitochondrion organization | 7 | 7.52E-05 | 9.3297 | Slc30a9, Ep300, Pisd, Rhot2, Rhot1, Mief2, Sirt7 |
| GOBP | positive regulation of axon extension | 8 | 6.76E-04 | 5.3312 | Ep300, Ilk, Fn1, Gdi1, Eif2b2, Rufy3, Twf2, Anapc2 |
| GOBP | lipid metabolic process | 40 | 9.14E-04 | 1.7456 | Mvk, Ppard, Pgap6, Mlycd, Plpp5, Gba2, Srebf2, Pisd, Pi4k2a, Pcyt2, Arsa, Ptpn11, Inpp5e, Mtmr2, Hsd3b7, Mbtps1, Pip4p1, Bscl2, Lypla2, Cbr4, Cyp27a1, Acadm, Ldlr, Prkag1, Gpld1, Acads, Tafazzin, Neu1, Adipor1, Nr1h2, Faah, Tecr, Srebf1, Chkb, Elovl1, Msmo1, Acaa1a, Plbd1, Decr2, Isyna1 |
| GOBP | nucleotide-excision repair | 7 | 9.20E-04 | 6.0517 | Gtf2h4, Ddb2, Trp53, Xpc, Ercc3, Gtf2h2, Fancc |
| GOBP | blastocyst development | 7 | 0.0014 | 5.5978 | Capn2, Psmc4, Thoc2, Xab2, Cdk11b, Emg1, Ppp1cc |
| GOBP | mRNA splicing, via spliceosome | 15 | 0.0015 | 2.6805 | Ddx23, Ddx39a, Prpf31, Wdr83, Sf3b1, Syf2, Xab2, Ddx41, Ik, Bcas2, Srsf5, Ddx46, Srrm1, Srsf6, Hnrnpm |
| GOBP | chromatin organization | 24 | 0.0016 | 2.0582 | Cdan1, Ing4, Hdac10, Kdm5d, Parp10, Prmt5, L3mbtl2, Hdac3, Lrwd1, Dmap1, Trim28, Pwwp3a, Mrgbp, Sirt7, 0610010K14Rik, Bap1, Men1, Suv39h1, Mettl23, Rccd1, Chd8, Setdb1, Arid2, Dapk3 |
| GOBP | cell cycle | 36 | 0.0017 | 1.7474 | Ep300, Rgs2, Septin9, Anapc5, Incenp, Stk11, Anapc2, Llgl2, Bub3, Ppp1cc, Jtb, Cdk5, Cep250, Suv39h1, Mau2, Nek4, Nae1, Cdk11b, Ing4, Camk1, Trp53, Ccng2, Golga2, Rack1, Cdk20, Tpr, Anxa11, Ist1, Mis18a, Becn1, Spout1, Cdk4, Ncapd3, Tbrg1, Katnb1, Reep4 |
| GOBP | methylation | 15 | 0.0017 | 2.6509 | Trmt1l, Nnmt, Prmt5, Trmt61a, Comtd1, Emg1, Cmtr1, Trmt2b, Suv39h1, Spout1, Mettl23, Gnmt, Hemk1, Setdb1, Trmt1 |
| GOBP | cell division | 25 | 0.0020 | 1.9893 | Septin9, Anapc5, Incenp, Anapc2, Llgl2, Bub3, Ppp1cc, Jtb, Cdk5, Men1, Mau2, Nek4, Ccng2, Golga2, Cdk20, Tpr, Anxa11, Ist1, Mis18a, Becn1, Spout1, Cdk4, Ncapd3, Katnb1, Reep4 |
| GOBP | regulation of gluconeogenesis | 5 | 0.0025 | 8.4178 | Acadm, Xpc, Gnmt, Slc35b4, Sirt7 |
| GOBP | autophagy | 16 | 0.0027 | 2.4256 | Rab24, Atg9a, Atg4b, Hdac10, Tmem208, Trp53, Rmc1, Gramd1a, Stk11, Arsa, Wdr24, Gopc, Tbc1d25, Becn1, Tbc1d17, Vps11 |
| GOBP | positive regulation of protein binding | 10 | 0.0027 | 3.4029 | Mark3, Ep300, Cdk5, Plk2, Hsp90ab1, Agrn, Men1, Ufl1, Trim28, Arhgef7 |
| GOBP | proline metabolic process | 3 | 0.0029 | 31.9875 | Prodh, Prodh2, Aldh4a1 |
| GOBP | proline catabolic process to glutamate | 3 | 0.0029 | 31.9875 | Prodh, Prodh2, Aldh4a1 |
| KEGG | Metabolic pathways | 88 | 5.64E-06 | 1.5779 | Nagk, Npr2, Gss, Gmppb, Pde9a, Pycr2, Gba2, Gpaa1, Pisd, Selenbp2, Adpgk, Coq6, Inpp5e, Selenbp1, Mtmr2, Hsd3b7, Prodh, Suv39h1, Tkt, Mat2a, Csgalnact2, Atp6v0a2, Pnpo, Aldoa, Acadm, Pck2, Nnmt, Gpld1, Dlst, Aldh4a1, Acads, Ftcd, Impdh2, Galt, Tecr, Acp1, Uros, Pdhb, Gnmt, Itpa, Hexdc, Mvk, Idh3g, Mlycd, Atp6v1h, Amd1, Ndufv1, Colgalt1, Adcy4, Pi4k2a, Gaa, Pcyt2, Arsa, Sirt7, Gpt, Atp6ap1, Pomt1, Entpd2, Mgat4b, Ugt2b34, Pomt2, Alas1, Entpd8, Cbr4, Ears2, Cyp27a1, Ugdh, Acy1, Acp2, Gfus, Dhrs4, Hpd, Mars1, Mppe1, Neu1, Phykpl, Ndufs2, Pigq, Chkb, Aldh8a1, Elovl1, Msmo1, Prodh2, Mocs2, Nudt13, Acaa1a, Setdb1, Isyna1 |
| KEGG | Lysosome | 17 | 1.44E-05 | 3.6601 | Man2b1, Atp6v1h, Slc11a1, Acp2, Ap1g2, Gaa, M6pr, Arsa, Neu1, Ctsa, Atp6ap1, Atp6v0a2, Ap4m1, Cln3, Ctsh, Ctsf, Tpp1 |
| KEGG | Spliceosome | 15 | 0.0036 | 2.4221 | Ddx23, Prpf31, Sf3b1, Syf2, Xab2, Rbm25, Bcas2, Srsf5, Ddx46, Thoc2, Srsf6, Cherp, Ddx5, Hnrnpm, Prpf18 |
| KEGG | Biosynthesis of cofactors | 12 | 0.0154 | 2.2946 | Ears2, Ugdh, Uros, Mat2a, Pnpo, Gss, Gmppb, Mocs2, Ugt2b34, Hpd, Alas1, Coq6 |
| KEGG | Apoptosis | 11 | 0.0184 | 2.3509 | Parp3, Diablo, Trp53, Capn2, Parp2, Ctsf, Gadd45g, Ctsh, Htra2, Aifm1, Raf1 |
| KEGG | Proteasome | 6 | 0.0217 | 3.7105 | Psme1, Psmd3, Psmc6, Psma3, Psmc4, Psmb9 |
| KEGG | Other glycan degradation | 4 | 0.0223 | 6.4590 | Man2b1, Gba2, Hexdc, Neu1 |
| KEGG | 2-Oxocarboxylic acid metabolism | 5 | 0.0280 | 4.2743 | Gpt, Idh3g, Acy1, Pdhb, Dlst |
| KEGG | Mitophagy - animal | 7 | 0.0287 | 2.9920 | Atg9a, Becn1, Trp53, Tfe3, Tbc1d17, Rhot2, Rhot1 |
| KEGG | AMPK signaling pathway | 10 | 0.0304 | 2.2886 | Mlycd, Akt1s1, Srebf1, Pck2, Prkag1, Tsc2, Stk11, Ppp2r5d, Adipor1, Rps6kb2 |
Fig. S6. Enrichment analysis of 687 genes including IR loci of reverse V shape in the 1st replicate
In the first group, 687 genes including 776 intron loci that were reverse V-shape (IR was significantly increased at KL- compared to WT and significantly decreased with kampo treatment) were subjected to GO biological process and KEGG pathway terms using DAVID.
Fig.S6

### Slide 7
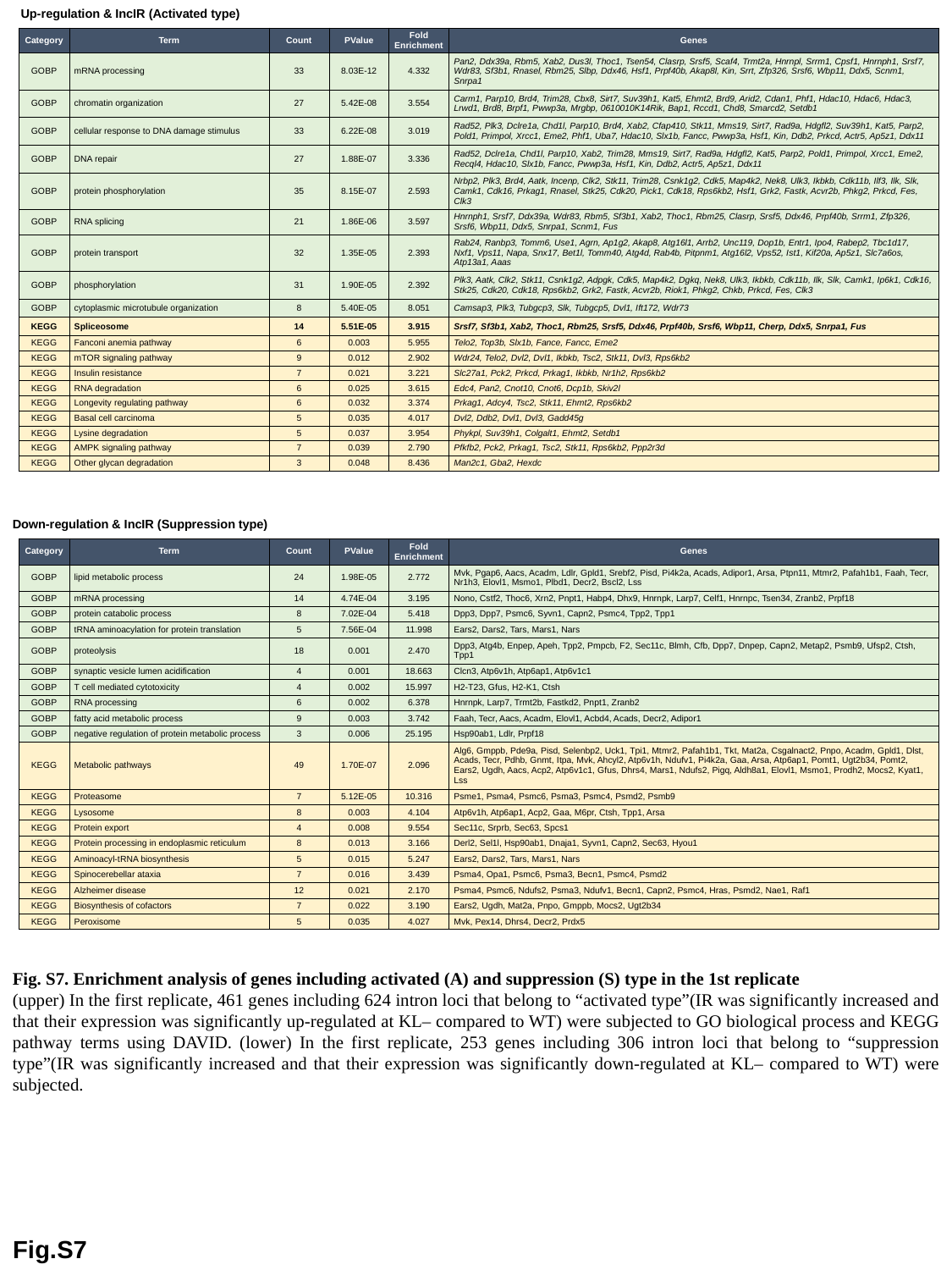

Up-regulation & IncIR (Activated type)
| Category | Term | Count | PValue | Fold Enrichment | Genes |
| --- | --- | --- | --- | --- | --- |
| GOBP | mRNA processing | 33 | 8.03E-12 | 4.332 | Pan2, Ddx39a, Rbm5, Xab2, Dus3l, Thoc1, Tsen54, Clasrp, Srsf5, Scaf4, Trmt2a, Hnrnpl, Srrm1, Cpsf1, Hnrnph1, Srsf7, Wdr83, Sf3b1, Rnasel, Rbm25, Slbp, Ddx46, Hsf1, Prpf40b, Akap8l, Kin, Srrt, Zfp326, Srsf6, Wbp11, Ddx5, Scnm1, Snrpa1 |
| GOBP | chromatin organization | 27 | 5.42E-08 | 3.554 | Carm1, Parp10, Brd4, Trim28, Cbx8, Sirt7, Suv39h1, Kat5, Ehmt2, Brd9, Arid2, Cdan1, Phf1, Hdac10, Hdac6, Hdac3, Lrwd1, Brd8, Brpf1, Pwwp3a, Mrgbp, 0610010K14Rik, Bap1, Rccd1, Chd8, Smarcd2, Setdb1 |
| GOBP | cellular response to DNA damage stimulus | 33 | 6.22E-08 | 3.019 | Rad52, Plk3, Dclre1a, Chd1l, Parp10, Brd4, Xab2, Cfap410, Stk11, Mms19, Sirt7, Rad9a, Hdgfl2, Suv39h1, Kat5, Parp2, Pold1, Primpol, Xrcc1, Eme2, Phf1, Uba7, Hdac10, Slx1b, Fancc, Pwwp3a, Hsf1, Kin, Ddb2, Prkcd, Actr5, Ap5z1, Ddx11 |
| GOBP | DNA repair | 27 | 1.88E-07 | 3.336 | Rad52, Dclre1a, Chd1l, Parp10, Xab2, Trim28, Mms19, Sirt7, Rad9a, Hdgfl2, Kat5, Parp2, Pold1, Primpol, Xrcc1, Eme2, Recql4, Hdac10, Slx1b, Fancc, Pwwp3a, Hsf1, Kin, Ddb2, Actr5, Ap5z1, Ddx11 |
| GOBP | protein phosphorylation | 35 | 8.15E-07 | 2.593 | Nrbp2, Plk3, Brd4, Aatk, Incenp, Clk2, Stk11, Trim28, Csnk1g2, Cdk5, Map4k2, Nek8, Ulk3, Ikbkb, Cdk11b, Ilf3, Ilk, Slk, Camk1, Cdk16, Prkag1, Rnasel, Stk25, Cdk20, Pick1, Cdk18, Rps6kb2, Hsf1, Grk2, Fastk, Acvr2b, Phkg2, Prkcd, Fes, Clk3 |
| GOBP | RNA splicing | 21 | 1.86E-06 | 3.597 | Hnrnph1, Srsf7, Ddx39a, Wdr83, Rbm5, Sf3b1, Xab2, Thoc1, Rbm25, Clasrp, Srsf5, Ddx46, Prpf40b, Srrm1, Zfp326, Srsf6, Wbp11, Ddx5, Snrpa1, Scnm1, Fus |
| GOBP | protein transport | 32 | 1.35E-05 | 2.393 | Rab24, Ranbp3, Tomm6, Use1, Agrn, Ap1g2, Akap8, Atg16l1, Arrb2, Unc119, Dop1b, Entr1, Ipo4, Rabep2, Tbc1d17, Nxf1, Vps11, Napa, Snx17, Bet1l, Tomm40, Atg4d, Rab4b, Pitpnm1, Atg16l2, Vps52, Ist1, Kif20a, Ap5z1, Slc7a6os, Atp13a1, Aaas |
| GOBP | phosphorylation | 31 | 1.90E-05 | 2.392 | Plk3, Aatk, Clk2, Stk11, Csnk1g2, Adpgk, Cdk5, Map4k2, Dgkq, Nek8, Ulk3, Ikbkb, Cdk11b, Ilk, Slk, Camk1, Ip6k1, Cdk16, Stk25, Cdk20, Cdk18, Rps6kb2, Grk2, Fastk, Acvr2b, Riok1, Phkg2, Chkb, Prkcd, Fes, Clk3 |
| GOBP | cytoplasmic microtubule organization | 8 | 5.40E-05 | 8.051 | Camsap3, Plk3, Tubgcp3, Slk, Tubgcp5, Dvl1, Ift172, Wdr73 |
| KEGG | Spliceosome | 14 | 5.51E-05 | 3.915 | Srsf7, Sf3b1, Xab2, Thoc1, Rbm25, Srsf5, Ddx46, Prpf40b, Srsf6, Wbp11, Cherp, Ddx5, Snrpa1, Fus |
| KEGG | Fanconi anemia pathway | 6 | 0.003 | 5.955 | Telo2, Top3b, Slx1b, Fance, Fancc, Eme2 |
| KEGG | mTOR signaling pathway | 9 | 0.012 | 2.902 | Wdr24, Telo2, Dvl2, Dvl1, Ikbkb, Tsc2, Stk11, Dvl3, Rps6kb2 |
| KEGG | Insulin resistance | 7 | 0.021 | 3.221 | Slc27a1, Pck2, Prkcd, Prkag1, Ikbkb, Nr1h2, Rps6kb2 |
| KEGG | RNA degradation | 6 | 0.025 | 3.615 | Edc4, Pan2, Cnot10, Cnot6, Dcp1b, Skiv2l |
| KEGG | Longevity regulating pathway | 6 | 0.032 | 3.374 | Prkag1, Adcy4, Tsc2, Stk11, Ehmt2, Rps6kb2 |
| KEGG | Basal cell carcinoma | 5 | 0.035 | 4.017 | Dvl2, Ddb2, Dvl1, Dvl3, Gadd45g |
| KEGG | Lysine degradation | 5 | 0.037 | 3.954 | Phykpl, Suv39h1, Colgalt1, Ehmt2, Setdb1 |
| KEGG | AMPK signaling pathway | 7 | 0.039 | 2.790 | Pfkfb2, Pck2, Prkag1, Tsc2, Stk11, Rps6kb2, Ppp2r3d |
| KEGG | Other glycan degradation | 3 | 0.048 | 8.436 | Man2c1, Gba2, Hexdc |
Down-regulation & IncIR (Suppression type)
| Category | Term | Count | PValue | Fold Enrichment | Genes |
| --- | --- | --- | --- | --- | --- |
| GOBP | lipid metabolic process | 24 | 1.98E-05 | 2.772 | Mvk, Pgap6, Aacs, Acadm, Ldlr, Gpld1, Srebf2, Pisd, Pi4k2a, Acads, Adipor1, Arsa, Ptpn11, Mtmr2, Pafah1b1, Faah, Tecr, Nr1h3, Elovl1, Msmo1, Plbd1, Decr2, Bscl2, Lss |
| GOBP | mRNA processing | 14 | 4.74E-04 | 3.195 | Nono, Cstf2, Thoc6, Xrn2, Pnpt1, Habp4, Dhx9, Hnrnpk, Larp7, Celf1, Hnrnpc, Tsen34, Zranb2, Prpf18 |
| GOBP | protein catabolic process | 8 | 7.02E-04 | 5.418 | Dpp3, Dpp7, Psmc6, Syvn1, Capn2, Psmc4, Tpp2, Tpp1 |
| GOBP | tRNA aminoacylation for protein translation | 5 | 7.56E-04 | 11.998 | Ears2, Dars2, Tars, Mars1, Nars |
| GOBP | proteolysis | 18 | 0.001 | 2.470 | Dpp3, Atg4b, Enpep, Apeh, Tpp2, Pmpcb, F2, Sec11c, Blmh, Cfb, Dpp7, Dnpep, Capn2, Metap2, Psmb9, Ufsp2, Ctsh, Tpp1 |
| GOBP | synaptic vesicle lumen acidification | 4 | 0.001 | 18.663 | Clcn3, Atp6v1h, Atp6ap1, Atp6v1c1 |
| GOBP | T cell mediated cytotoxicity | 4 | 0.002 | 15.997 | H2-T23, Gfus, H2-K1, Ctsh |
| GOBP | RNA processing | 6 | 0.002 | 6.378 | Hnrnpk, Larp7, Trmt2b, Fastkd2, Pnpt1, Zranb2 |
| GOBP | fatty acid metabolic process | 9 | 0.003 | 3.742 | Faah, Tecr, Aacs, Acadm, Elovl1, Acbd4, Acads, Decr2, Adipor1 |
| GOBP | negative regulation of protein metabolic process | 3 | 0.006 | 25.195 | Hsp90ab1, Ldlr, Prpf18 |
| KEGG | Metabolic pathways | 49 | 1.70E-07 | 2.096 | Alg6, Gmppb, Pde9a, Pisd, Selenbp2, Uck1, Tpi1, Mtmr2, Pafah1b1, Tkt, Mat2a, Csgalnact2, Pnpo, Acadm, Gpld1, Dlst, Acads, Tecr, Pdhb, Gnmt, Itpa, Mvk, Ahcyl2, Atp6v1h, Ndufv1, Pi4k2a, Gaa, Arsa, Atp6ap1, Pomt1, Ugt2b34, Pomt2, Ears2, Ugdh, Aacs, Acp2, Atp6v1c1, Gfus, Dhrs4, Mars1, Ndufs2, Pigq, Aldh8a1, Elovl1, Msmo1, Prodh2, Mocs2, Kyat1, Lss |
| KEGG | Proteasome | 7 | 5.12E-05 | 10.316 | Psme1, Psma4, Psmc6, Psma3, Psmc4, Psmd2, Psmb9 |
| KEGG | Lysosome | 8 | 0.003 | 4.104 | Atp6v1h, Atp6ap1, Acp2, Gaa, M6pr, Ctsh, Tpp1, Arsa |
| KEGG | Protein export | 4 | 0.008 | 9.554 | Sec11c, Srprb, Sec63, Spcs1 |
| KEGG | Protein processing in endoplasmic reticulum | 8 | 0.013 | 3.166 | Derl2, Sel1l, Hsp90ab1, Dnaja1, Syvn1, Capn2, Sec63, Hyou1 |
| KEGG | Aminoacyl-tRNA biosynthesis | 5 | 0.015 | 5.247 | Ears2, Dars2, Tars, Mars1, Nars |
| KEGG | Spinocerebellar ataxia | 7 | 0.016 | 3.439 | Psma4, Opa1, Psmc6, Psma3, Becn1, Psmc4, Psmd2 |
| KEGG | Alzheimer disease | 12 | 0.021 | 2.170 | Psma4, Psmc6, Ndufs2, Psma3, Ndufv1, Becn1, Capn2, Psmc4, Hras, Psmd2, Nae1, Raf1 |
| KEGG | Biosynthesis of cofactors | 7 | 0.022 | 3.190 | Ears2, Ugdh, Mat2a, Pnpo, Gmppb, Mocs2, Ugt2b34 |
| KEGG | Peroxisome | 5 | 0.035 | 4.027 | Mvk, Pex14, Dhrs4, Decr2, Prdx5 |
Fig. S7. Enrichment analysis of genes including activated (A) and suppression (S) type in the 1st replicate
(upper) In the first replicate, 461 genes including 624 intron loci that belong to “activated type”(IR was significantly increased and that their expression was significantly up-regulated at KL– compared to WT) were subjected to GO biological process and KEGG pathway terms using DAVID. (lower) In the first replicate, 253 genes including 306 intron loci that belong to “suppression type”(IR was significantly increased and that their expression was significantly down-regulated at KL– compared to WT) were subjected.
Fig.S7

### Slide 8
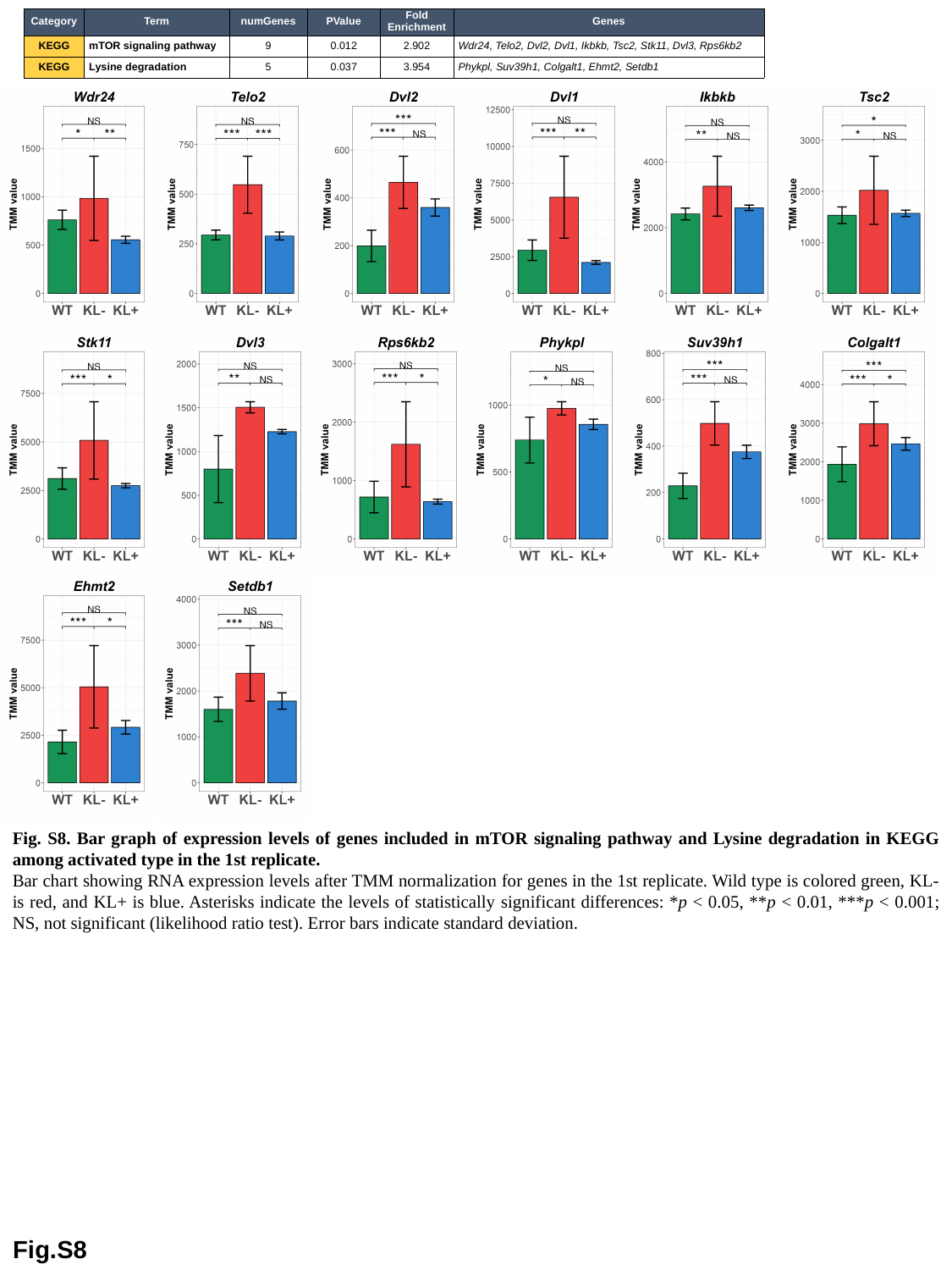

| Category | Term | numGenes | PValue | Fold Enrichment | Genes |
| --- | --- | --- | --- | --- | --- |
| KEGG | mTOR signaling pathway | 9 | 0.012 | 2.902 | Wdr24, Telo2, Dvl2, Dvl1, Ikbkb, Tsc2, Stk11, Dvl3, Rps6kb2 |
| KEGG | Lysine degradation | 5 | 0.037 | 3.954 | Phykpl, Suv39h1, Colgalt1, Ehmt2, Setdb1 |
Fig. S8. Bar graph of expression levels of genes included in mTOR signaling pathway and Lysine degradation in KEGG among activated type in the 1st replicate.
Bar chart showing RNA expression levels after TMM normalization for genes in the 1st replicate. Wild type is colored green, KL- is red, and KL+ is blue. Asterisks indicate the levels of statistically significant differences: *p < 0.05, **p < 0.01, ***p < 0.001; NS, not significant (likelihood ratio test). Error bars indicate standard deviation.
Fig.S8

### Slide 9
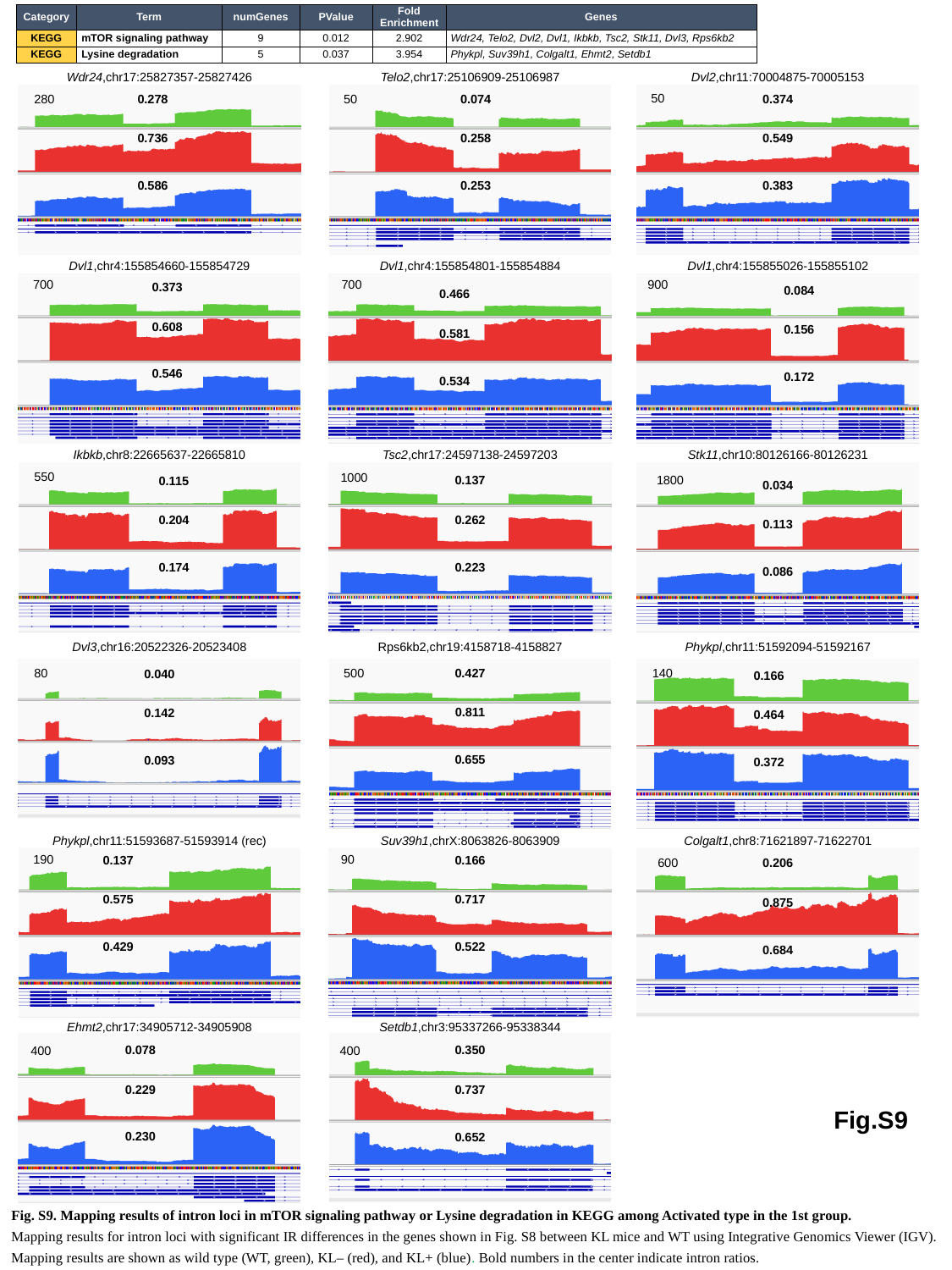

| Category | Term | numGenes | PValue | Fold Enrichment | Genes |
| --- | --- | --- | --- | --- | --- |
| KEGG | mTOR signaling pathway | 9 | 0.012 | 2.902 | Wdr24, Telo2, Dvl2, Dvl1, Ikbkb, Tsc2, Stk11, Dvl3, Rps6kb2 |
| KEGG | Lysine degradation | 5 | 0.037 | 3.954 | Phykpl, Suv39h1, Colgalt1, Ehmt2, Setdb1 |
Wdr24,chr17:25827357-25827426
Telo2,chr17:25106909-25106987
Dvl2,chr11:70004875-70005153
50
0.278
0.074
0.374
280
50
0.736
0.258
0.549
0.586
0.253
0.383
Dvl1,chr4:155854660-155854729
Dvl1,chr4:155854801-155854884
Dvl1,chr4:155855026-155855102
700
700
900
0.373
0.084
0.466
0.608
0.156
0.581
0.546
0.172
0.534
Ikbkb,chr8:22665637-22665810
Tsc2,chr17:24597138-24597203
Stk11,chr10:80126166-80126231
550
1000
1800
0.137
0.115
0.034
0.262
0.204
0.113
0.223
0.174
0.086
Dvl3,chr16:20522326-20523408
Rps6kb2,chr19:4158718-4158827
Phykpl,chr11:51592094-51592167
80
500
140
0.427
0.040
0.166
0.811
0.142
0.464
0.655
0.093
0.372
Phykpl,chr11:51593687-51593914 (rec)
Suv39h1,chrX:8063826-8063909
Colgalt1,chr8:71621897-71622701
90
190
0.166
0.137
600
0.206
0.717
0.575
0.875
0.522
0.429
0.684
Ehmt2,chr17:34905712-34905908
Setdb1,chr3:95337266-95338344
0.078
0.350
400
400
0.229
0.737
Fig.S9
0.230
0.652
Fig. S9. Mapping results of intron loci in mTOR signaling pathway or Lysine degradation in KEGG among Activated type in the 1st group.
Mapping results for intron loci with significant IR differences in the genes shown in Fig. S8 between KL mice and WT using Integrative Genomics Viewer (IGV). Mapping results are shown as wild type (WT, green), KL– (red), and KL+ (blue). Bold numbers in the center indicate intron ratios.

### Slide 10
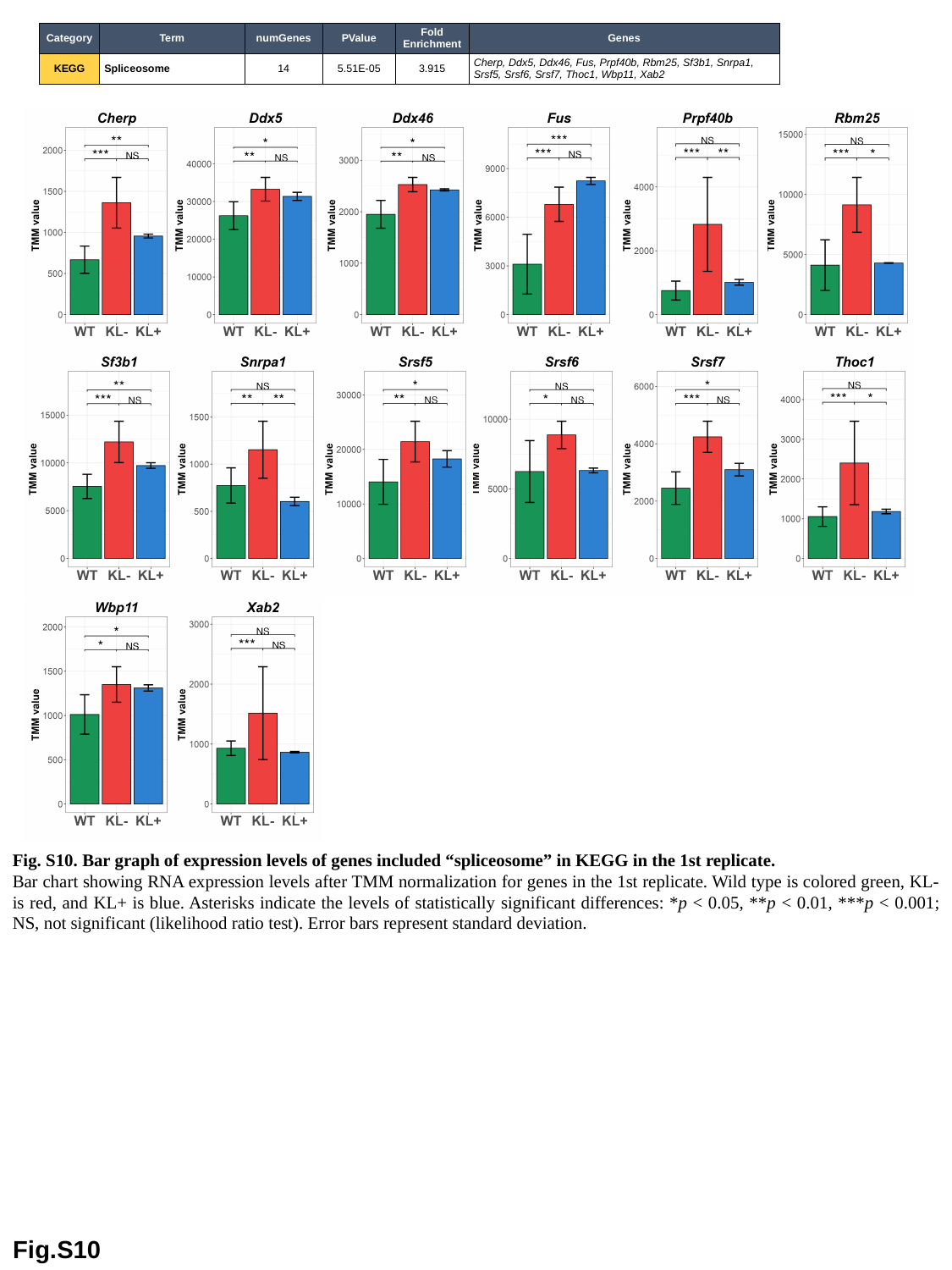

| Category | Term | numGenes | PValue | Fold Enrichment | Genes |
| --- | --- | --- | --- | --- | --- |
| KEGG | Spliceosome | 14 | 5.51E-05 | 3.915 | Cherp, Ddx5, Ddx46, Fus, Prpf40b, Rbm25, Sf3b1, Snrpa1, Srsf5, Srsf6, Srsf7, Thoc1, Wbp11, Xab2 |
Fig. S10. Bar graph of expression levels of genes included “spliceosome” in KEGG in the 1st replicate.
Bar chart showing RNA expression levels after TMM normalization for genes in the 1st replicate. Wild type is colored green, KL- is red, and KL+ is blue. Asterisks indicate the levels of statistically significant differences: *p < 0.05, **p < 0.01, ***p < 0.001; NS, not significant (likelihood ratio test). Error bars represent standard deviation.
Fig.S10

### Slide 11
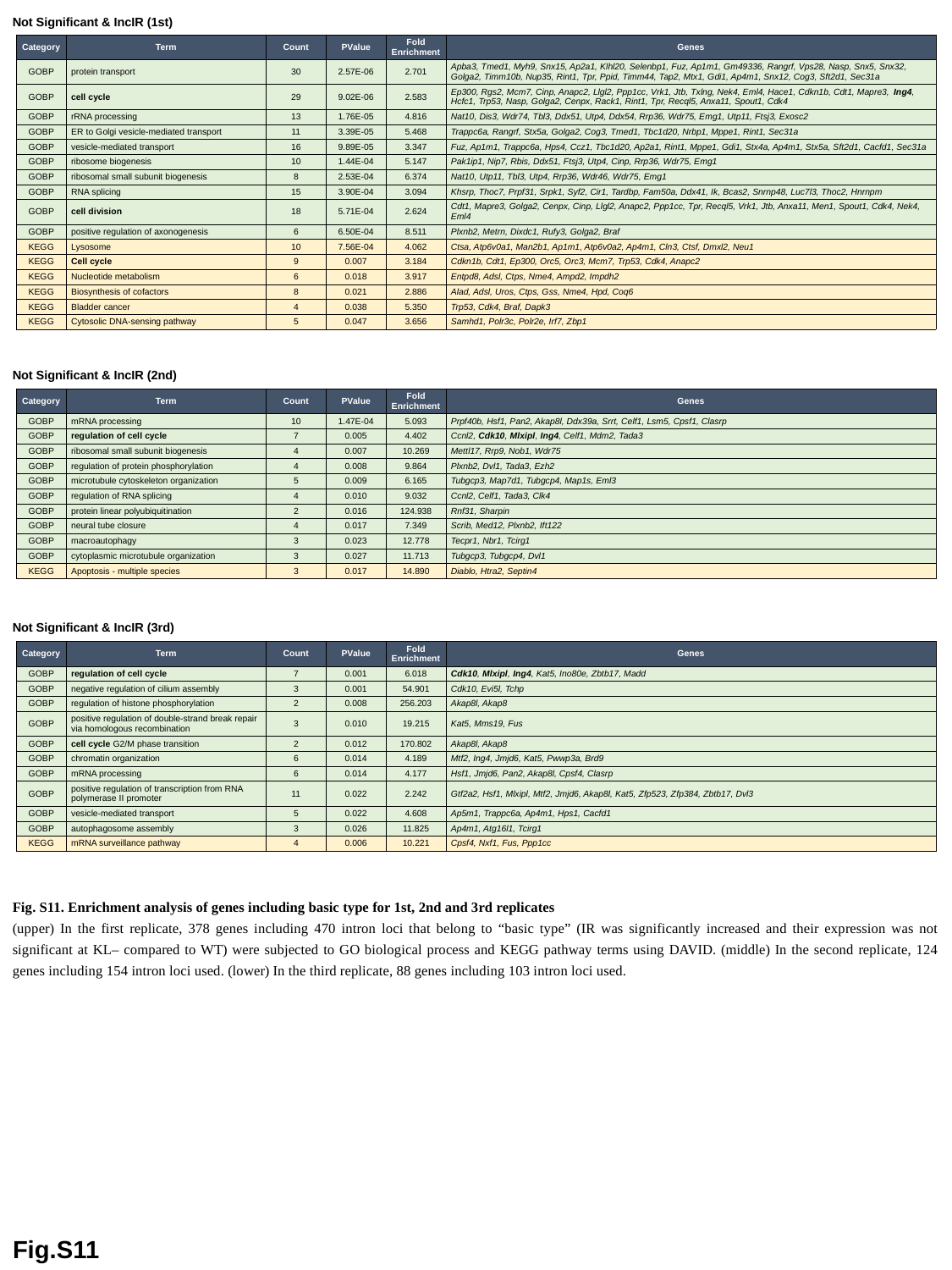

Not Significant & IncIR (1st)
| Category | Term | Count | PValue | Fold Enrichment | Genes |
| --- | --- | --- | --- | --- | --- |
| GOBP | protein transport | 30 | 2.57E-06 | 2.701 | Apba3, Tmed1, Myh9, Snx15, Ap2a1, Klhl20, Selenbp1, Fuz, Ap1m1, Gm49336, Rangrf, Vps28, Nasp, Snx5, Snx32, Golga2, Timm10b, Nup35, Rint1, Tpr, Ppid, Timm44, Tap2, Mtx1, Gdi1, Ap4m1, Snx12, Cog3, Sft2d1, Sec31a |
| GOBP | cell cycle | 29 | 9.02E-06 | 2.583 | Ep300, Rgs2, Mcm7, Cinp, Anapc2, Llgl2, Ppp1cc, Vrk1, Jtb, Txlng, Nek4, Eml4, Hace1, Cdkn1b, Cdt1, Mapre3, Ing4, Hcfc1, Trp53, Nasp, Golga2, Cenpx, Rack1, Rint1, Tpr, Recql5, Anxa11, Spout1, Cdk4 |
| GOBP | rRNA processing | 13 | 1.76E-05 | 4.816 | Nat10, Dis3, Wdr74, Tbl3, Ddx51, Utp4, Ddx54, Rrp36, Wdr75, Emg1, Utp11, Ftsj3, Exosc2 |
| GOBP | ER to Golgi vesicle-mediated transport | 11 | 3.39E-05 | 5.468 | Trappc6a, Rangrf, Stx5a, Golga2, Cog3, Tmed1, Tbc1d20, Nrbp1, Mppe1, Rint1, Sec31a |
| GOBP | vesicle-mediated transport | 16 | 9.89E-05 | 3.347 | Fuz, Ap1m1, Trappc6a, Hps4, Ccz1, Tbc1d20, Ap2a1, Rint1, Mppe1, Gdi1, Stx4a, Ap4m1, Stx5a, Sft2d1, Cacfd1, Sec31a |
| GOBP | ribosome biogenesis | 10 | 1.44E-04 | 5.147 | Pak1ip1, Nip7, Rbis, Ddx51, Ftsj3, Utp4, Cinp, Rrp36, Wdr75, Emg1 |
| GOBP | ribosomal small subunit biogenesis | 8 | 2.53E-04 | 6.374 | Nat10, Utp11, Tbl3, Utp4, Rrp36, Wdr46, Wdr75, Emg1 |
| GOBP | RNA splicing | 15 | 3.90E-04 | 3.094 | Khsrp, Thoc7, Prpf31, Srpk1, Syf2, Cir1, Tardbp, Fam50a, Ddx41, Ik, Bcas2, Snrnp48, Luc7l3, Thoc2, Hnrnpm |
| GOBP | cell division | 18 | 5.71E-04 | 2.624 | Cdt1, Mapre3, Golga2, Cenpx, Cinp, Llgl2, Anapc2, Ppp1cc, Tpr, Recql5, Vrk1, Jtb, Anxa11, Men1, Spout1, Cdk4, Nek4, Eml4 |
| GOBP | positive regulation of axonogenesis | 6 | 6.50E-04 | 8.511 | Plxnb2, Metrn, Dixdc1, Rufy3, Golga2, Braf |
| KEGG | Lysosome | 10 | 7.56E-04 | 4.062 | Ctsa, Atp6v0a1, Man2b1, Ap1m1, Atp6v0a2, Ap4m1, Cln3, Ctsf, Dmxl2, Neu1 |
| KEGG | Cell cycle | 9 | 0.007 | 3.184 | Cdkn1b, Cdt1, Ep300, Orc5, Orc3, Mcm7, Trp53, Cdk4, Anapc2 |
| KEGG | Nucleotide metabolism | 6 | 0.018 | 3.917 | Entpd8, Adsl, Ctps, Nme4, Ampd2, Impdh2 |
| KEGG | Biosynthesis of cofactors | 8 | 0.021 | 2.886 | Alad, Adsl, Uros, Ctps, Gss, Nme4, Hpd, Coq6 |
| KEGG | Bladder cancer | 4 | 0.038 | 5.350 | Trp53, Cdk4, Braf, Dapk3 |
| KEGG | Cytosolic DNA-sensing pathway | 5 | 0.047 | 3.656 | Samhd1, Polr3c, Polr2e, Irf7, Zbp1 |
Not Significant & IncIR (2nd)
| Category | Term | Count | PValue | Fold Enrichment | Genes |
| --- | --- | --- | --- | --- | --- |
| GOBP | mRNA processing | 10 | 1.47E-04 | 5.093 | Prpf40b, Hsf1, Pan2, Akap8l, Ddx39a, Srrt, Celf1, Lsm5, Cpsf1, Clasrp |
| GOBP | regulation of cell cycle | 7 | 0.005 | 4.402 | Ccnl2, Cdk10, Mlxipl, Ing4, Celf1, Mdm2, Tada3 |
| GOBP | ribosomal small subunit biogenesis | 4 | 0.007 | 10.269 | Mettl17, Rrp9, Nob1, Wdr75 |
| GOBP | regulation of protein phosphorylation | 4 | 0.008 | 9.864 | Plxnb2, Dvl1, Tada3, Ezh2 |
| GOBP | microtubule cytoskeleton organization | 5 | 0.009 | 6.165 | Tubgcp3, Map7d1, Tubgcp4, Map1s, Eml3 |
| GOBP | regulation of RNA splicing | 4 | 0.010 | 9.032 | Ccnl2, Celf1, Tada3, Clk4 |
| GOBP | protein linear polyubiquitination | 2 | 0.016 | 124.938 | Rnf31, Sharpin |
| GOBP | neural tube closure | 4 | 0.017 | 7.349 | Scrib, Med12, Plxnb2, Ift122 |
| GOBP | macroautophagy | 3 | 0.023 | 12.778 | Tecpr1, Nbr1, Tcirg1 |
| GOBP | cytoplasmic microtubule organization | 3 | 0.027 | 11.713 | Tubgcp3, Tubgcp4, Dvl1 |
| KEGG | Apoptosis - multiple species | 3 | 0.017 | 14.890 | Diablo, Htra2, Septin4 |
Not Significant & IncIR (3rd)
| Category | Term | Count | PValue | Fold Enrichment | Genes |
| --- | --- | --- | --- | --- | --- |
| GOBP | regulation of cell cycle | 7 | 0.001 | 6.018 | Cdk10, Mlxipl, Ing4, Kat5, Ino80e, Zbtb17, Madd |
| GOBP | negative regulation of cilium assembly | 3 | 0.001 | 54.901 | Cdk10, Evi5l, Tchp |
| GOBP | regulation of histone phosphorylation | 2 | 0.008 | 256.203 | Akap8l, Akap8 |
| GOBP | positive regulation of double-strand break repair via homologous recombination | 3 | 0.010 | 19.215 | Kat5, Mms19, Fus |
| GOBP | cell cycle G2/M phase transition | 2 | 0.012 | 170.802 | Akap8l, Akap8 |
| GOBP | chromatin organization | 6 | 0.014 | 4.189 | Mtf2, Ing4, Jmjd6, Kat5, Pwwp3a, Brd9 |
| GOBP | mRNA processing | 6 | 0.014 | 4.177 | Hsf1, Jmjd6, Pan2, Akap8l, Cpsf4, Clasrp |
| GOBP | positive regulation of transcription from RNA polymerase II promoter | 11 | 0.022 | 2.242 | Gtf2a2, Hsf1, Mlxipl, Mtf2, Jmjd6, Akap8l, Kat5, Zfp523, Zfp384, Zbtb17, Dvl3 |
| GOBP | vesicle-mediated transport | 5 | 0.022 | 4.608 | Ap5m1, Trappc6a, Ap4m1, Hps1, Cacfd1 |
| GOBP | autophagosome assembly | 3 | 0.026 | 11.825 | Ap4m1, Atg16l1, Tcirg1 |
| KEGG | mRNA surveillance pathway | 4 | 0.006 | 10.221 | Cpsf4, Nxf1, Fus, Ppp1cc |
Fig. S11. Enrichment analysis of genes including basic type for 1st, 2nd and 3rd replicates
(upper) In the first replicate, 378 genes including 470 intron loci that belong to “basic type” (IR was significantly increased and their expression was not significant at KL– compared to WT) were subjected to GO biological process and KEGG pathway terms using DAVID. (middle) In the second replicate, 124 genes including 154 intron loci used. (lower) In the third replicate, 88 genes including 103 intron loci used.
Fig.S11

### Slide 12
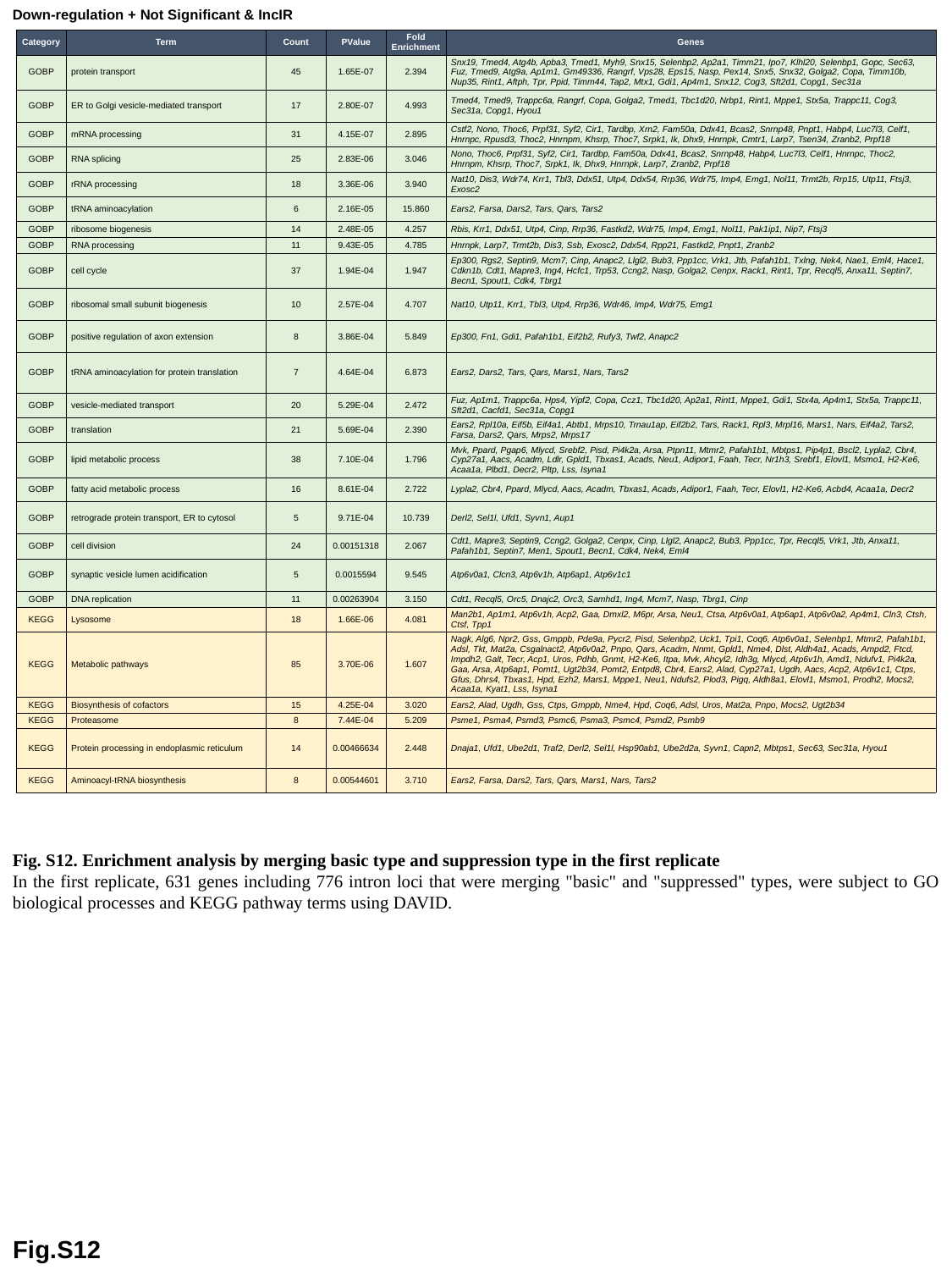

Down-regulation + Not Significant & IncIR
| Category | Term | Count | PValue | Fold Enrichment | Genes |
| --- | --- | --- | --- | --- | --- |
| GOBP | protein transport | 45 | 1.65E-07 | 2.394 | Snx19, Tmed4, Atg4b, Apba3, Tmed1, Myh9, Snx15, Selenbp2, Ap2a1, Timm21, Ipo7, Klhl20, Selenbp1, Gopc, Sec63, Fuz, Tmed9, Atg9a, Ap1m1, Gm49336, Rangrf, Vps28, Eps15, Nasp, Pex14, Snx5, Snx32, Golga2, Copa, Timm10b, Nup35, Rint1, Aftph, Tpr, Ppid, Timm44, Tap2, Mtx1, Gdi1, Ap4m1, Snx12, Cog3, Sft2d1, Copg1, Sec31a |
| GOBP | ER to Golgi vesicle-mediated transport | 17 | 2.80E-07 | 4.993 | Tmed4, Tmed9, Trappc6a, Rangrf, Copa, Golga2, Tmed1, Tbc1d20, Nrbp1, Rint1, Mppe1, Stx5a, Trappc11, Cog3, Sec31a, Copg1, Hyou1 |
| GOBP | mRNA processing | 31 | 4.15E-07 | 2.895 | Cstf2, Nono, Thoc6, Prpf31, Syf2, Cir1, Tardbp, Xrn2, Fam50a, Ddx41, Bcas2, Snrnp48, Pnpt1, Habp4, Luc7l3, Celf1, Hnrnpc, Rpusd3, Thoc2, Hnrnpm, Khsrp, Thoc7, Srpk1, Ik, Dhx9, Hnrnpk, Cmtr1, Larp7, Tsen34, Zranb2, Prpf18 |
| GOBP | RNA splicing | 25 | 2.83E-06 | 3.046 | Nono, Thoc6, Prpf31, Syf2, Cir1, Tardbp, Fam50a, Ddx41, Bcas2, Snrnp48, Habp4, Luc7l3, Celf1, Hnrnpc, Thoc2, Hnrnpm, Khsrp, Thoc7, Srpk1, Ik, Dhx9, Hnrnpk, Larp7, Zranb2, Prpf18 |
| GOBP | rRNA processing | 18 | 3.36E-06 | 3.940 | Nat10, Dis3, Wdr74, Krr1, Tbl3, Ddx51, Utp4, Ddx54, Rrp36, Wdr75, Imp4, Emg1, Nol11, Trmt2b, Rrp15, Utp11, Ftsj3, Exosc2 |
| GOBP | tRNA aminoacylation | 6 | 2.16E-05 | 15.860 | Ears2, Farsa, Dars2, Tars, Qars, Tars2 |
| GOBP | ribosome biogenesis | 14 | 2.48E-05 | 4.257 | Rbis, Krr1, Ddx51, Utp4, Cinp, Rrp36, Fastkd2, Wdr75, Imp4, Emg1, Nol11, Pak1ip1, Nip7, Ftsj3 |
| GOBP | RNA processing | 11 | 9.43E-05 | 4.785 | Hnrnpk, Larp7, Trmt2b, Dis3, Ssb, Exosc2, Ddx54, Rpp21, Fastkd2, Pnpt1, Zranb2 |
| GOBP | cell cycle | 37 | 1.94E-04 | 1.947 | Ep300, Rgs2, Septin9, Mcm7, Cinp, Anapc2, Llgl2, Bub3, Ppp1cc, Vrk1, Jtb, Pafah1b1, Txlng, Nek4, Nae1, Eml4, Hace1, Cdkn1b, Cdt1, Mapre3, Ing4, Hcfc1, Trp53, Ccng2, Nasp, Golga2, Cenpx, Rack1, Rint1, Tpr, Recql5, Anxa11, Septin7, Becn1, Spout1, Cdk4, Tbrg1 |
| GOBP | ribosomal small subunit biogenesis | 10 | 2.57E-04 | 4.707 | Nat10, Utp11, Krr1, Tbl3, Utp4, Rrp36, Wdr46, Imp4, Wdr75, Emg1 |
| GOBP | positive regulation of axon extension | 8 | 3.86E-04 | 5.849 | Ep300, Fn1, Gdi1, Pafah1b1, Eif2b2, Rufy3, Twf2, Anapc2 |
| GOBP | tRNA aminoacylation for protein translation | 7 | 4.64E-04 | 6.873 | Ears2, Dars2, Tars, Qars, Mars1, Nars, Tars2 |
| GOBP | vesicle-mediated transport | 20 | 5.29E-04 | 2.472 | Fuz, Ap1m1, Trappc6a, Hps4, Yipf2, Copa, Ccz1, Tbc1d20, Ap2a1, Rint1, Mppe1, Gdi1, Stx4a, Ap4m1, Stx5a, Trappc11, Sft2d1, Cacfd1, Sec31a, Copg1 |
| GOBP | translation | 21 | 5.69E-04 | 2.390 | Ears2, Rpl10a, Eif5b, Eif4a1, Abtb1, Mrps10, Trnau1ap, Eif2b2, Tars, Rack1, Rpl3, Mrpl16, Mars1, Nars, Eif4a2, Tars2, Farsa, Dars2, Qars, Mrps2, Mrps17 |
| GOBP | lipid metabolic process | 38 | 7.10E-04 | 1.796 | Mvk, Ppard, Pgap6, Mlycd, Srebf2, Pisd, Pi4k2a, Arsa, Ptpn11, Mtmr2, Pafah1b1, Mbtps1, Pip4p1, Bscl2, Lypla2, Cbr4, Cyp27a1, Aacs, Acadm, Ldlr, Gpld1, Tbxas1, Acads, Neu1, Adipor1, Faah, Tecr, Nr1h3, Srebf1, Elovl1, Msmo1, H2-Ke6, Acaa1a, Plbd1, Decr2, Pltp, Lss, Isyna1 |
| GOBP | fatty acid metabolic process | 16 | 8.61E-04 | 2.722 | Lypla2, Cbr4, Ppard, Mlycd, Aacs, Acadm, Tbxas1, Acads, Adipor1, Faah, Tecr, Elovl1, H2-Ke6, Acbd4, Acaa1a, Decr2 |
| GOBP | retrograde protein transport, ER to cytosol | 5 | 9.71E-04 | 10.739 | Derl2, Sel1l, Ufd1, Syvn1, Aup1 |
| GOBP | cell division | 24 | 0.00151318 | 2.067 | Cdt1, Mapre3, Septin9, Ccng2, Golga2, Cenpx, Cinp, Llgl2, Anapc2, Bub3, Ppp1cc, Tpr, Recql5, Vrk1, Jtb, Anxa11, Pafah1b1, Septin7, Men1, Spout1, Becn1, Cdk4, Nek4, Eml4 |
| GOBP | synaptic vesicle lumen acidification | 5 | 0.0015594 | 9.545 | Atp6v0a1, Clcn3, Atp6v1h, Atp6ap1, Atp6v1c1 |
| GOBP | DNA replication | 11 | 0.00263904 | 3.150 | Cdt1, Recql5, Orc5, Dnajc2, Orc3, Samhd1, Ing4, Mcm7, Nasp, Tbrg1, Cinp |
| KEGG | Lysosome | 18 | 1.66E-06 | 4.081 | Man2b1, Ap1m1, Atp6v1h, Acp2, Gaa, Dmxl2, M6pr, Arsa, Neu1, Ctsa, Atp6v0a1, Atp6ap1, Atp6v0a2, Ap4m1, Cln3, Ctsh, Ctsf, Tpp1 |
| KEGG | Metabolic pathways | 85 | 3.70E-06 | 1.607 | Nagk, Alg6, Npr2, Gss, Gmppb, Pde9a, Pycr2, Pisd, Selenbp2, Uck1, Tpi1, Coq6, Atp6v0a1, Selenbp1, Mtmr2, Pafah1b1, Adsl, Tkt, Mat2a, Csgalnact2, Atp6v0a2, Pnpo, Qars, Acadm, Nnmt, Gpld1, Nme4, Dlst, Aldh4a1, Acads, Ampd2, Ftcd, Impdh2, Galt, Tecr, Acp1, Uros, Pdhb, Gnmt, H2-Ke6, Itpa, Mvk, Ahcyl2, Idh3g, Mlycd, Atp6v1h, Amd1, Ndufv1, Pi4k2a, Gaa, Arsa, Atp6ap1, Pomt1, Ugt2b34, Pomt2, Entpd8, Cbr4, Ears2, Alad, Cyp27a1, Ugdh, Aacs, Acp2, Atp6v1c1, Ctps, Gfus, Dhrs4, Tbxas1, Hpd, Ezh2, Mars1, Mppe1, Neu1, Ndufs2, Plod3, Pigq, Aldh8a1, Elovl1, Msmo1, Prodh2, Mocs2, Acaa1a, Kyat1, Lss, Isyna1 |
| KEGG | Biosynthesis of cofactors | 15 | 4.25E-04 | 3.020 | Ears2, Alad, Ugdh, Gss, Ctps, Gmppb, Nme4, Hpd, Coq6, Adsl, Uros, Mat2a, Pnpo, Mocs2, Ugt2b34 |
| KEGG | Proteasome | 8 | 7.44E-04 | 5.209 | Psme1, Psma4, Psmd3, Psmc6, Psma3, Psmc4, Psmd2, Psmb9 |
| KEGG | Protein processing in endoplasmic reticulum | 14 | 0.00466634 | 2.448 | Dnaja1, Ufd1, Ube2d1, Traf2, Derl2, Sel1l, Hsp90ab1, Ube2d2a, Syvn1, Capn2, Mbtps1, Sec63, Sec31a, Hyou1 |
| KEGG | Aminoacyl-tRNA biosynthesis | 8 | 0.00544601 | 3.710 | Ears2, Farsa, Dars2, Tars, Qars, Mars1, Nars, Tars2 |
Fig. S12. Enrichment analysis by merging basic type and suppression type in the first replicate
In the first replicate, 631 genes including 776 intron loci that were merging "basic" and "suppressed" types, were subject to GO biological processes and KEGG pathway terms using DAVID.
Fig.S12

### Slide 13
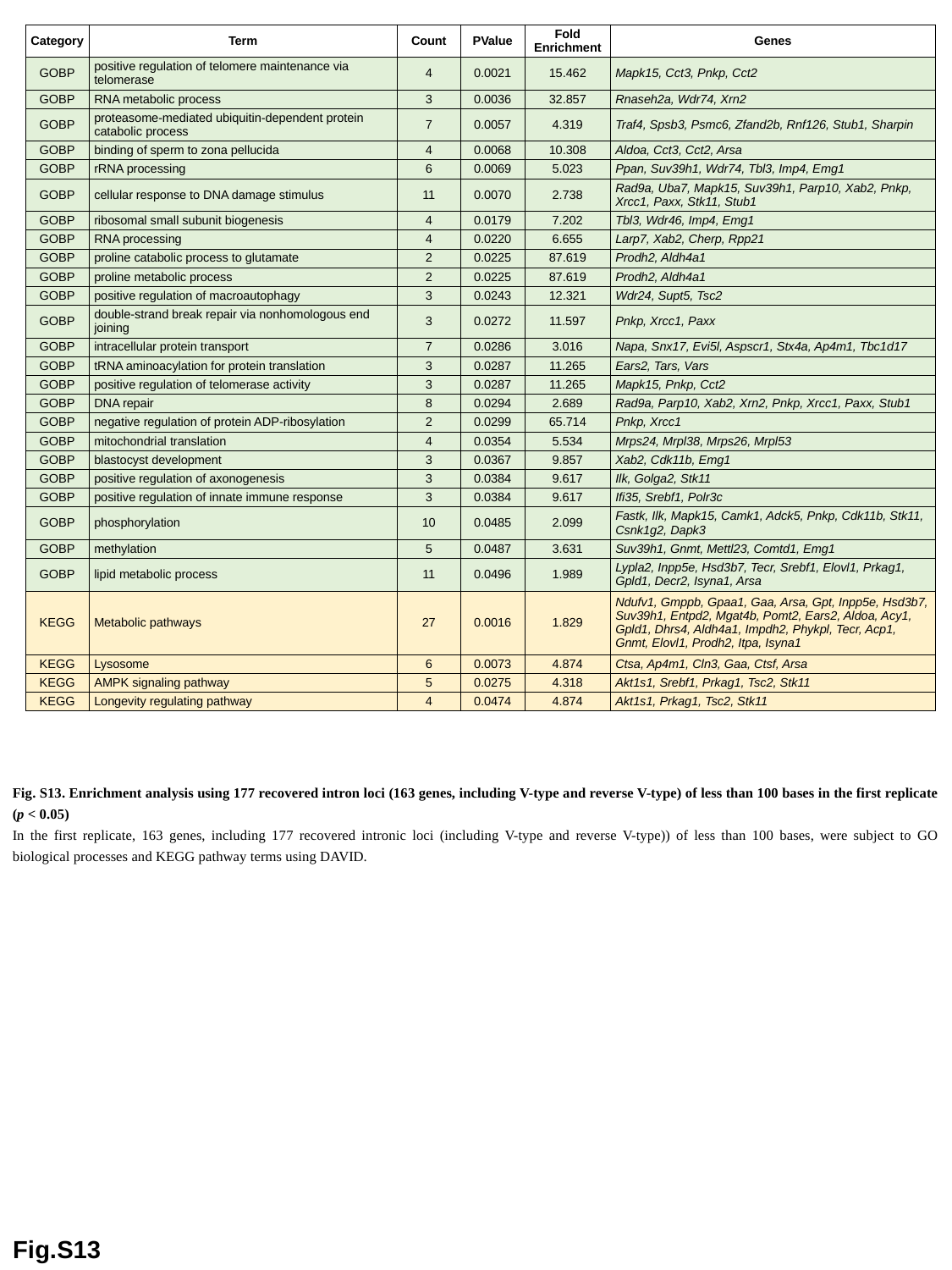

| Category | Term | Count | PValue | Fold Enrichment | Genes |
| --- | --- | --- | --- | --- | --- |
| GOBP | positive regulation of telomere maintenance via telomerase | 4 | 0.0021 | 15.462 | Mapk15, Cct3, Pnkp, Cct2 |
| GOBP | RNA metabolic process | 3 | 0.0036 | 32.857 | Rnaseh2a, Wdr74, Xrn2 |
| GOBP | proteasome-mediated ubiquitin-dependent protein catabolic process | 7 | 0.0057 | 4.319 | Traf4, Spsb3, Psmc6, Zfand2b, Rnf126, Stub1, Sharpin |
| GOBP | binding of sperm to zona pellucida | 4 | 0.0068 | 10.308 | Aldoa, Cct3, Cct2, Arsa |
| GOBP | rRNA processing | 6 | 0.0069 | 5.023 | Ppan, Suv39h1, Wdr74, Tbl3, Imp4, Emg1 |
| GOBP | cellular response to DNA damage stimulus | 11 | 0.0070 | 2.738 | Rad9a, Uba7, Mapk15, Suv39h1, Parp10, Xab2, Pnkp, Xrcc1, Paxx, Stk11, Stub1 |
| GOBP | ribosomal small subunit biogenesis | 4 | 0.0179 | 7.202 | Tbl3, Wdr46, Imp4, Emg1 |
| GOBP | RNA processing | 4 | 0.0220 | 6.655 | Larp7, Xab2, Cherp, Rpp21 |
| GOBP | proline catabolic process to glutamate | 2 | 0.0225 | 87.619 | Prodh2, Aldh4a1 |
| GOBP | proline metabolic process | 2 | 0.0225 | 87.619 | Prodh2, Aldh4a1 |
| GOBP | positive regulation of macroautophagy | 3 | 0.0243 | 12.321 | Wdr24, Supt5, Tsc2 |
| GOBP | double-strand break repair via nonhomologous end joining | 3 | 0.0272 | 11.597 | Pnkp, Xrcc1, Paxx |
| GOBP | intracellular protein transport | 7 | 0.0286 | 3.016 | Napa, Snx17, Evi5l, Aspscr1, Stx4a, Ap4m1, Tbc1d17 |
| GOBP | tRNA aminoacylation for protein translation | 3 | 0.0287 | 11.265 | Ears2, Tars, Vars |
| GOBP | positive regulation of telomerase activity | 3 | 0.0287 | 11.265 | Mapk15, Pnkp, Cct2 |
| GOBP | DNA repair | 8 | 0.0294 | 2.689 | Rad9a, Parp10, Xab2, Xrn2, Pnkp, Xrcc1, Paxx, Stub1 |
| GOBP | negative regulation of protein ADP-ribosylation | 2 | 0.0299 | 65.714 | Pnkp, Xrcc1 |
| GOBP | mitochondrial translation | 4 | 0.0354 | 5.534 | Mrps24, Mrpl38, Mrps26, Mrpl53 |
| GOBP | blastocyst development | 3 | 0.0367 | 9.857 | Xab2, Cdk11b, Emg1 |
| GOBP | positive regulation of axonogenesis | 3 | 0.0384 | 9.617 | Ilk, Golga2, Stk11 |
| GOBP | positive regulation of innate immune response | 3 | 0.0384 | 9.617 | Ifi35, Srebf1, Polr3c |
| GOBP | phosphorylation | 10 | 0.0485 | 2.099 | Fastk, Ilk, Mapk15, Camk1, Adck5, Pnkp, Cdk11b, Stk11, Csnk1g2, Dapk3 |
| GOBP | methylation | 5 | 0.0487 | 3.631 | Suv39h1, Gnmt, Mettl23, Comtd1, Emg1 |
| GOBP | lipid metabolic process | 11 | 0.0496 | 1.989 | Lypla2, Inpp5e, Hsd3b7, Tecr, Srebf1, Elovl1, Prkag1, Gpld1, Decr2, Isyna1, Arsa |
| KEGG | Metabolic pathways | 27 | 0.0016 | 1.829 | Ndufv1, Gmppb, Gpaa1, Gaa, Arsa, Gpt, Inpp5e, Hsd3b7, Suv39h1, Entpd2, Mgat4b, Pomt2, Ears2, Aldoa, Acy1, Gpld1, Dhrs4, Aldh4a1, Impdh2, Phykpl, Tecr, Acp1, Gnmt, Elovl1, Prodh2, Itpa, Isyna1 |
| KEGG | Lysosome | 6 | 0.0073 | 4.874 | Ctsa, Ap4m1, Cln3, Gaa, Ctsf, Arsa |
| KEGG | AMPK signaling pathway | 5 | 0.0275 | 4.318 | Akt1s1, Srebf1, Prkag1, Tsc2, Stk11 |
| KEGG | Longevity regulating pathway | 4 | 0.0474 | 4.874 | Akt1s1, Prkag1, Tsc2, Stk11 |
Fig. S13. Enrichment analysis using 177 recovered intron loci (163 genes, including V-type and reverse V-type) of less than 100 bases in the first replicate (p < 0.05)
In the first replicate, 163 genes, including 177 recovered intronic loci (including V-type and reverse V-type)) of less than 100 bases, were subject to GO biological processes and KEGG pathway terms using DAVID.
Fig.S13

### Slide 14
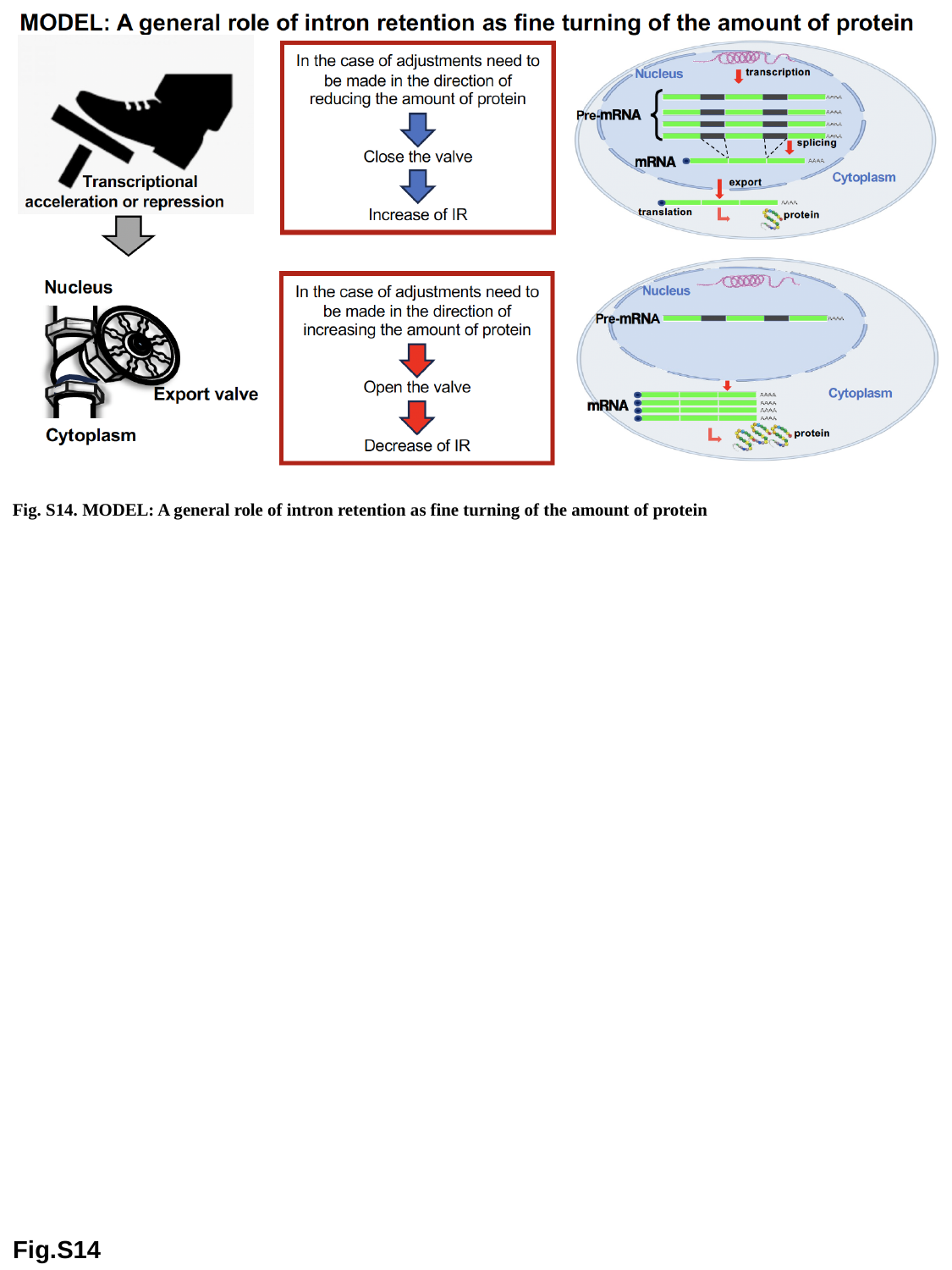

Fig. S14. MODEL: A general role of intron retention as fine turning of the amount of protein
Fig.S14

### Slide 15
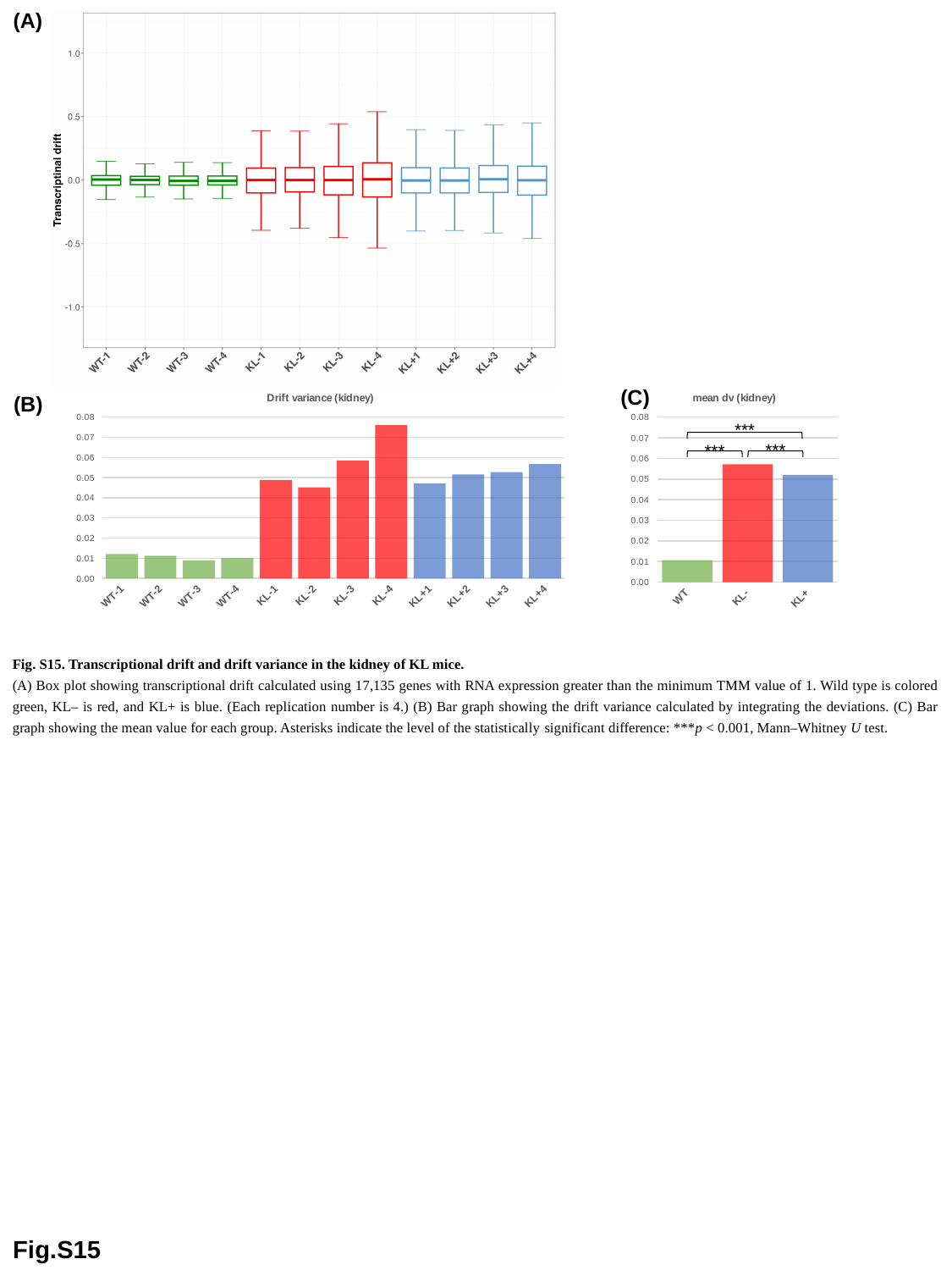

(A)
(C)
#### Chart: Drift variance (kidney)
| Category | dv |
|---|---|
| WT-1 | 0.012084050607975608 |
| WT-2 | 0.011260649637477227 |
| WT-3 | 0.008915806171838713 |
| WT-4 | 0.01017687116051383 |
| KL-1 | 0.04889947827432891 |
| KL-2 | 0.045153659983213176 |
| KL-3 | 0.05853005179741295 |
| KL-4 | 0.07608130814084763 |
| KL+1 | 0.04714301474018496 |
| KL+2 | 0.0515853691796437 |
| KL+3 | 0.05267973524999228 |
| KL+4 | 0.05675334799005152 |
#### Chart: mean dv (kidney)
| Category | dv |
|---|---|
| WT | 0.010609344394451386 |
| KL- | 0.05716612454895064 |
| KL+ | 0.05204036678996808 |(B)
***
***
***
Fig. S15. Transcriptional drift and drift variance in the kidney of KL mice.
(A) Box plot showing transcriptional drift calculated using 17,135 genes with RNA expression greater than the minimum TMM value of 1. Wild type is colored green, KL– is red, and KL+ is blue. (Each replication number is 4.) (B) Bar graph showing the drift variance calculated by integrating the deviations. (C) Bar graph showing the mean value for each group. Asterisks indicate the level of the statistically significant difference: ***p < 0.001, Mann–Whitney U test.
Fig.S15

### Slide 16
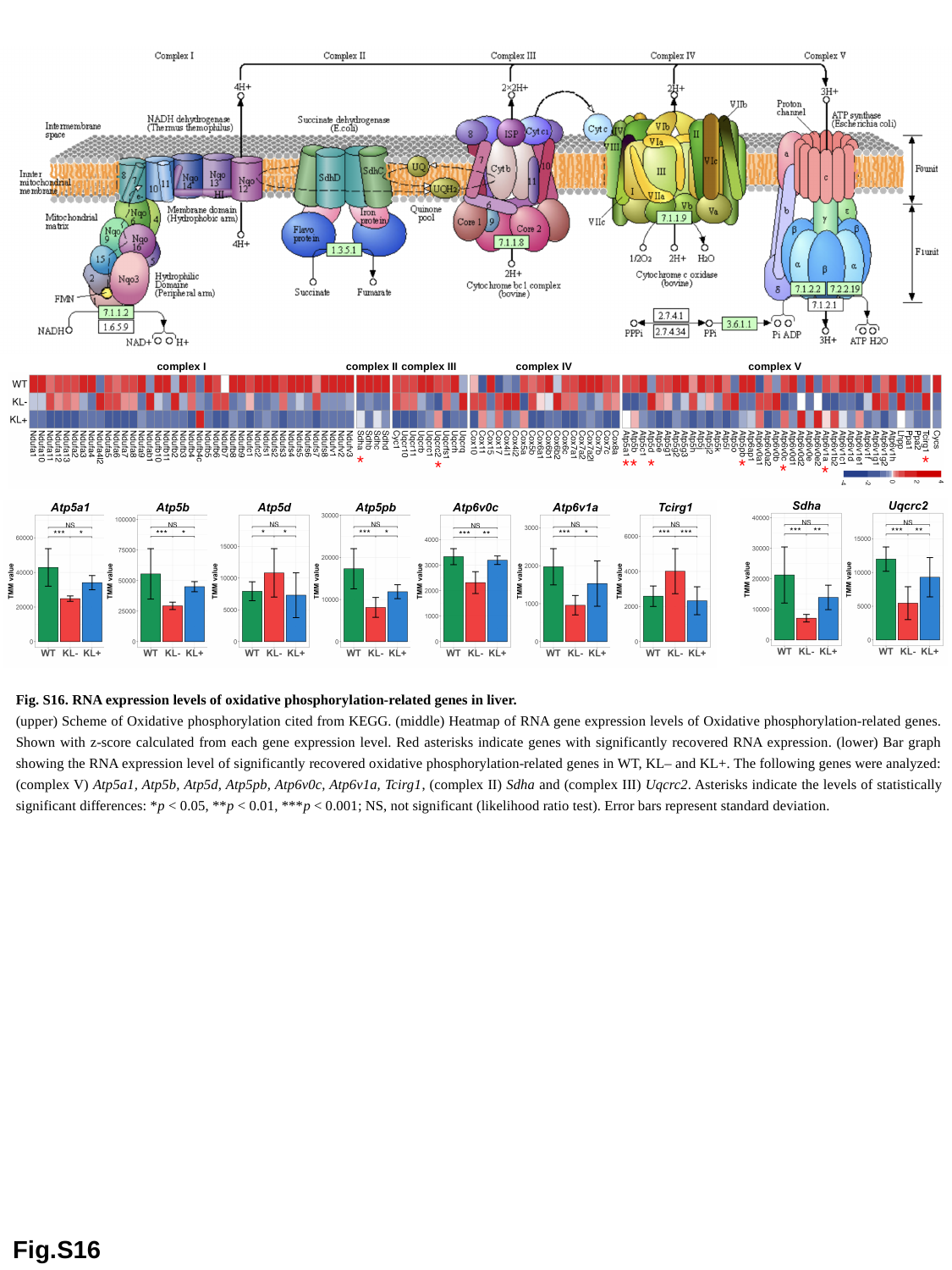

complex I
complex II
complex III
complex IV
complex V
*
*
*
*
*
*
*
*
*
Fig. S16. RNA expression levels of oxidative phosphorylation-related genes in liver.
(upper) Scheme of Oxidative phosphorylation cited from KEGG. (middle) Heatmap of RNA gene expression levels of Oxidative phosphorylation-related genes. Shown with z-score calculated from each gene expression level. Red asterisks indicate genes with significantly recovered RNA expression. (lower) Bar graph showing the RNA expression level of significantly recovered oxidative phosphorylation-related genes in WT, KL– and KL+. The following genes were analyzed: (complex V) Atp5a1, Atp5b, Atp5d, Atp5pb, Atp6v0c, Atp6v1a, Tcirg1, (complex II) Sdha and (complex III) Uqcrc2. Asterisks indicate the levels of statistically significant differences: *p < 0.05, **p < 0.01, ***p < 0.001; NS, not significant (likelihood ratio test). Error bars represent standard deviation.
Fig.S16
